# Discovery of knock-down resistance in the major African malaria vector *Anopheles funestus*

**DOI:** 10.1101/2024.03.13.584754

**Authors:** Joel O. Odero, Tristan P. W. Dennis, Brian Polo, Joachim Nwezeobi, Marilou Boddé, Sanjay C. Nagi, Anastasia Hernandez-Koutoucheva, Ismail H. Nambunga, Hamis Bwanary, Gustav Mkandawile, Nicodem J Govella, Emmanuel W. Kaindoa, Heather M. Ferguson, Eric Ochomo, Chris S. Clarkson, Alistair Miles, Mara K. N. Lawniczak, David Weetman, Francesco Baldini, Fredros O. Okumu

## Abstract

A major mechanism of insecticide resistance in insect pests is knock-down resistance (*kdr*) caused by mutations in the voltage-gated sodium channel (*Vgsc*) gene. Despite being common in most malaria *Anopheles* vector species, *kdr* mutations have never been observed in *Anopheles funestus*, the principal malaria vector in Eastern and Southern Africa. While monitoring 10 populations of *An. funestus* in Tanzania, we unexpectedly found resistance to DDT, a banned insecticide, in one location. Through whole-genome sequencing of 333 *An. funestus* samples from these populations, we found 8 novel amino acid substitutions in the *Vgsc* gene, including the *kdr* variant, L976F (L1014F in *An. gambiae*), in tight linkage disequilibrium with another (P1842S). The mutants were found only at high frequency in one region, with a significant decline between 2017 and 2023. Notably, *kdr* L976F was strongly associated with survivorship to the exposure to DDT insecticide, while no clear association was noted with a pyrethroid insecticide (deltamethrin). Further study is necessary to identify the origin and spread of *kdr* in *An. funestus*, and the potential threat to current insecticide-based vector control in Africa.

**Significance:** Knock-down resistance (kdr) mutations confer resistance to malaria vector control insecticides and pose a grave threat to malaria control. Here, we report the first discovery of kdr in *An. funestus*, the principal malaria vector in East and Southern Africa. Kdr in *An. funestus* conferred resistance to DDT but not deltamethrin. Based on extensive DDT contamination and unofficial usage in Tanzania, it is possible that kdr emerged because of widespread organic pollution as opposed to through public health efforts. Regardless of origin, the discovery of kdr in *An. funestus* is an alarming development that warrants immediate, extensive follow-up and close surveillance to establish the origin, and extent to which it may threaten malaria control in *An. funestus*.

## Introduction

Chemical insecticides are central to the control of agricultural pests and disease vectors. The control of *Anopheles* mosquitoes through the distribution of over 2.9 billion insecticide-treated bed nets (ITNs) has helped avert an estimated 633 million cases of malaria (*1*) - a disease that still kills 600,000 yearly (*2*). However, the widespread use of insecticides for agricultural pest and disease vector control also has detrimental consequences, including direct lethal and sub-lethal effects on human and animal health (*3, 4*) and destabilizing effects on ecosystem structure and function. For example, insecticide exposure is a key stressor affecting the population decline of pollinators, essential for ecosystem health and food production (*5, 6*).

A key obstacle to the sustainable control of malaria is the evolutionary arms race between mosquitoes and insecticide-based mosquito control. Strong selection pressures generated by insecticide-based agricultural pest and disease vector control activities have resulted in the independent evolution of a diverse range of mechanisms that confer insecticide resistance (IR) phenotypes in numerous insect species (*7*). One of the earliest described IR mechanisms was the emergence of knock-down resistance (*kdr*), mediated by mutations in the target site of pyrethroid and organochlorine insecticides, located in the voltage-gated sodium channel gene (*Vgsc*), an essential component of the nervous system (*8*). These *kdr-*driven resistance phenotypes appeared rapidly after the introduction of the organochlorine dichloro-diphenyl-trichloroethane (DDT) spraying for insect control in the mid-20th century (*9*) and eventually evolving to confer resistance to pyrethroids (*10, 11*), the key ingredient in ITNs - the first line of defence against malaria. In an era of stalling gains in malaria control (*2*), and concerted efforts both to develop a new generation of ITN and IRS products (*12, 13*) and proactively manage the deployment of existing insecticides to maximise efficacy, intensified surveillance, including genomic surveillance (*14, 15*), of malaria vector populations is critical for providing real-time warning of insecticide resistance emergence.

As part of phenotypic and genomic surveillance done in Tanzania to understand the evolution and spread of insecticide resistance in *Anopheles funestus -* the dominant malaria vector in Eastern and Southern Africa (*16*), we report the first discovery of *kdr* mutations in *An. funestus*. We discover that this mutation confers resistance to DDT, but not deltamethrin, despite a complete ban on DDT use for agriculture and vector control in Tanzania since 2008 (*17*). We suggest environmental contamination from extensive DDT stockpiles (*18*), or unofficial agricultural use, as possible causes. The emergence of *kdr*, which threatens the control of major crop pests and vectors of disease, such as *An. gambiae* and *Aedes aegypti* (*19*), highlights the potential of chemical insecticide contamination or unofficial use to exert unexpected and potentially harmful impacts on public health.

## Results

Resistance to all major classes of insecticide is common in *An. funestus* and is primarily mediated through the increased activity of enzymes that bind to and metabolise insecticides (metabolic resistance) (*20, 21*). This contrasts with another major vector *An. gambiae* where resistance is mostly conferred by a combination of metabolic and target-site resistance (*7*). As part of an insecticide resistance surveillance study in 10 sites across Tanzania (**Fig. 1A**) (*22*), we investigated phenotypic resistance (as measured by mosquito survival 24 hours following insecticide exposure) in *An. funestus* to the discriminating doses of DDT, deltamethrin (type II pyrethroid), or deltamethrin together with the piperonyl butoxide (PBO) synergist, which is increasingly used on ITNs (*23*) to restore susceptibility in pyrethroid-resistant populations in Tanzania. The mosquitoes were phenotypically resistant to deltamethrin in all regions, but PBO ubiquitously restored susceptibility. Unexpectedly, resistance to DDT was recorded in the Morogoro region (68%, CI 57.8 - 77.9), but not in other regions (**Fig. 1B**).

**Fig. 1:**
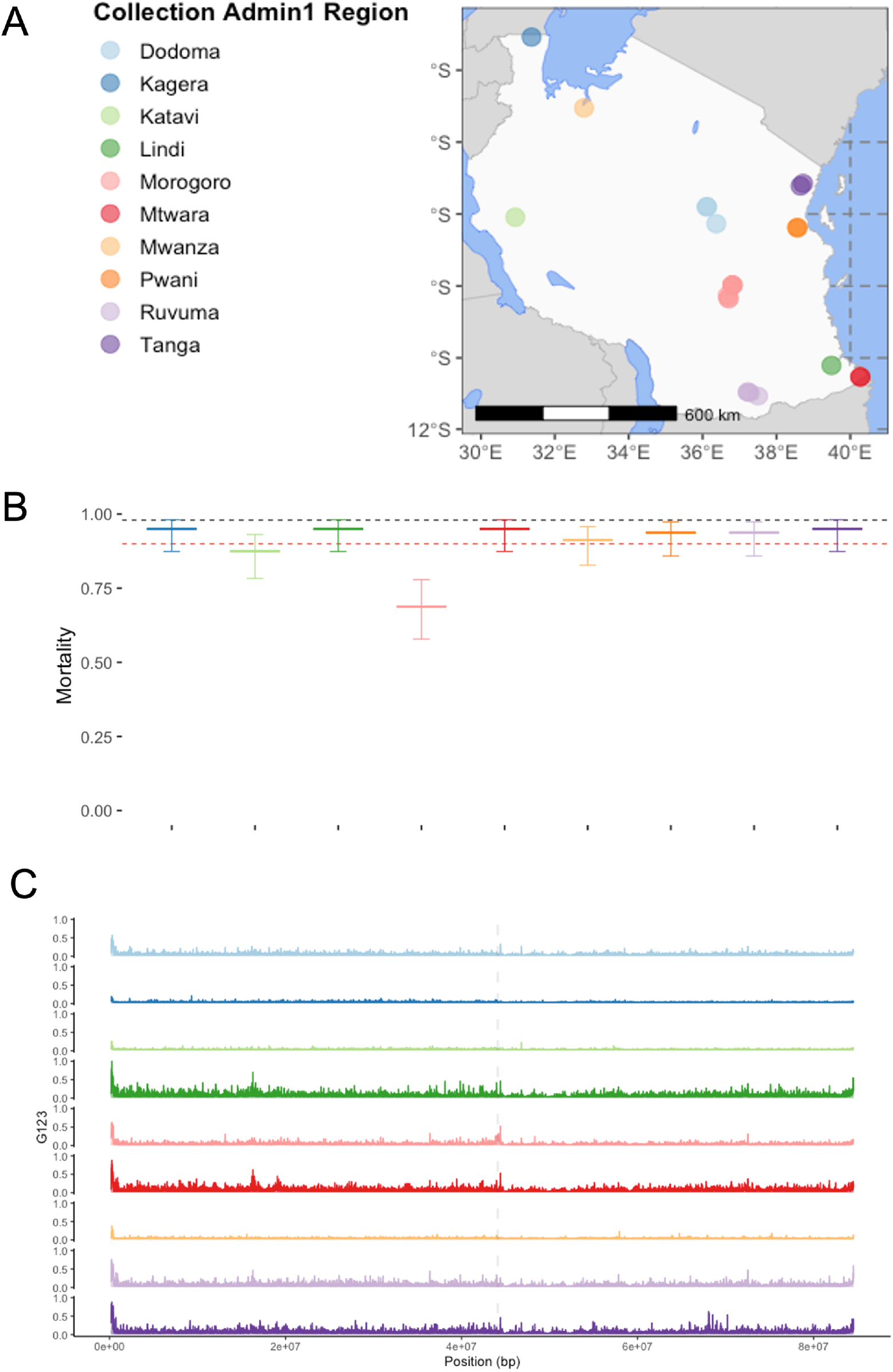
**(A)** Map of *An. funestus* collection locations. Points indicate sample collection locations. The point colour indicates the administrative region from which samples were collected. **(B)**: Phenotypic insecticide resistance profile of *An. funestus* to DDT using bioassay data adopted from our recent surveillance (*22*). The colours represent the various regions where the bioassays were conducted, and the error bars are 95% confidence interval. The black and red dotted lines on the y-axis represent the 98 and 90% mortality thresholds. (**C**) G123 selection scans of *An. funestus* chromosome 3RL, coloured and windowed by sample collection region (where n>20 – see **Supp Table 2**). X-axis indicates the position (in base-pairs (bp)), Y-axis indicates the selection statistic G123. The Grey dotted line indicates the location of the *Vgsc* gene. Note Mwanza region is absent from panel **C** as there were too few samples (n<20) to perform a selection scan.

To understand the genetic bases of resistance, we analysed whole-genome-sequencing (WGS) data from 333 mosquitoes sampled from 10 sites across Tanzania (**Fig. 1A**). We performed genome-wide selection scans (GWSS) with the G123 statistic to test for evidence of selective sweeps in the *An. funestus* genome associated with known or novel IR loci (**Fig. 1C**; grouping samples by administrative region (see **Supp. Table 1** for per-group sample numbers). We detected a clear signal of elevated G123 in the region containing the *Vgsc* gene in samples from the Morogoro region in the southeastern part of the country (**Fig. 1C**). Notably, *Vgsc* encodes for the voltage-gated sodium channel, where DDT binds in mosquitoes, and where mutations are strongly linked to resistance in *An. gambiae* (*11*). In Kagera, Katavi, and Mwanza regions, there was no visible sign of a selective sweep at or near the *Vgsc* region. In Dodoma, Lindi, Ruvuma, and Tanga, there were peaks of elevated G123 near to *Vgsc*, but these appeared within the context of relatively high G123 across the chromosome **(Fig. 1C)**.

**Table 1:**
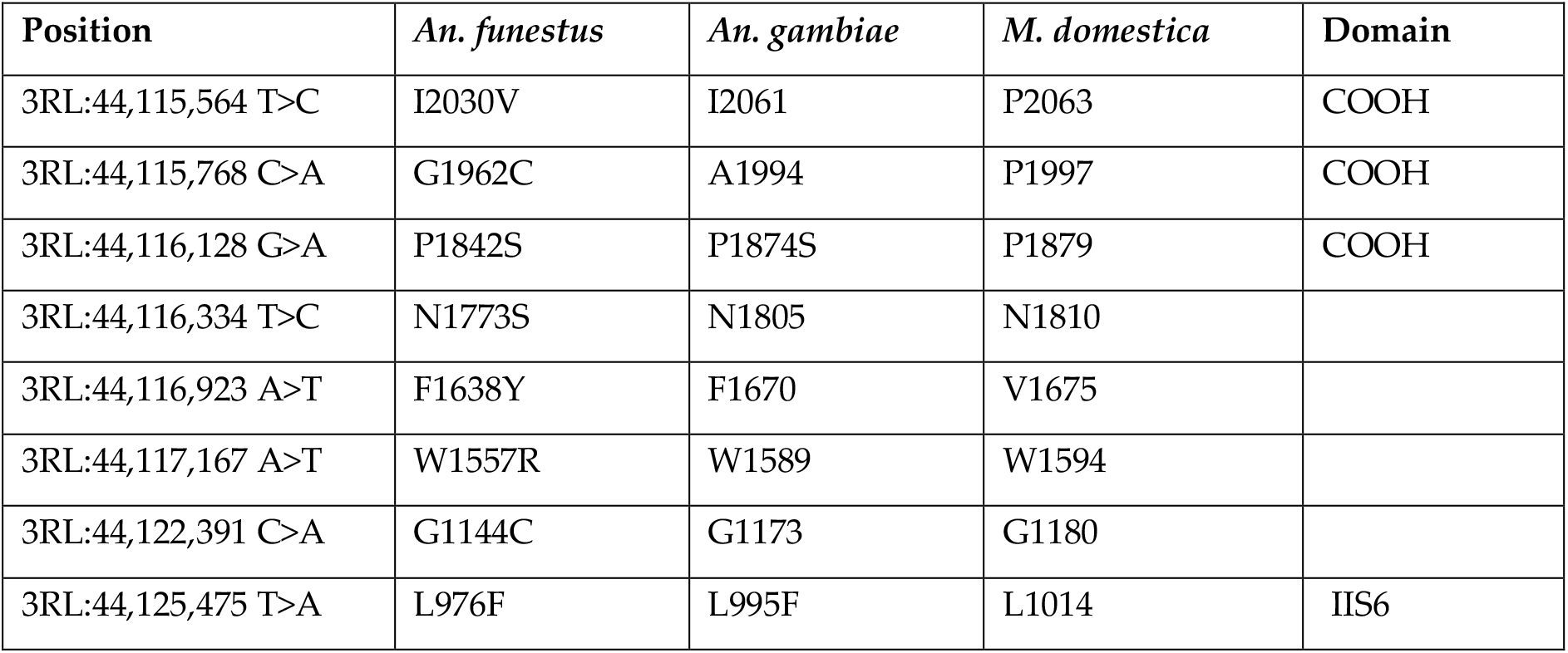
Comparative non-synonymous nucleotide variation in the voltage-gated sodium channel gene. The position is relative to the *Anopheles funestus* strain FUMOZ reference, chromosome arm 3RL. Codon numbering according to *Anopheles funestus Vgsc* transcript AFUN2_008728.R15290, *Anopheles gambiae* transcript AGAP004707-RD in gene set AgamP4.12, and *Musca domestica* EMBL accession X96668 Williamson *et al*. (*29*).

| Position | *An. funestus* | *An. gambiae* | *M. domestica* | Domain |
| --- | --- | --- | --- | --- |
| 3RL:44,115,564 T>C | I2030V | I2061 | P2063 | COOH |
| 3RL:44,115,768 C>A | G1962C | A1994 | P1997 | COOH |
| 3RL:44,116,128 G>A | P1842S | P1874S | P1879 | COOH |
| 3RL:44,116,334 T>C | N1773S | N1805 | N1810 |  |
| 3RL:44,116,923 A>T | F1638Y | F1670 | V1675 |  |
| 3RL:44,117,167 A>T | W1557R | W1589 | W1594 |  |
| 3RL:44,122,391 C>A | G1144C | G1173 | G1180 |  |
| 3RL:44,125,475 T>A | L976F | L995F | L1014 | IIS6 |

Mutations in *Vgsc* confer *kdr* in numerous pest and insect taxa (*24*). While not previously reported in *An. funestus*(*25*), *kdr* mutations in major malaria vectors within the *An. gambiae* complex are subject to intense selection (*26, 27*) and confer resistance to pyrethroid and organochlorine insecticides used in ITNs and insecticide sprays (*11, 24*). We searched our data for mutations in the *Vgsc* gene and found 8 amino acid substitutions occurring at frequencies greater than 5% (**Fig. 2A**). Of these, two alleles, L976F and P1842S occurred at the highest frequency (**Fig. 2A**). The frequencies of P1824S and L976F were highest in samples collected from Morogoro in 2017 (0.75 and 0.90 respectively) **(Fig. 2A)** and declined yearly, reaching their lowest frequency in samples collected in 2023 (0.48 and 0.56 respectively; χ^2^ = 12.15, p=0.0005; **Fig. 2B)**. These mutations occurred at very low frequencies or were absent in all other locations (**Fig. 2A)**. To understand their function, we aligned the *An. funestus Vgsc* sequence (Gene ID: AFUN2_008728.R15290) with that of *Musca domestica* (Gene ID: X96668) and *An. gambiae* (AGAP004707-RD AgamP4.12 gene set) (*27*). We found that the amino acid change at *An. funestus* L976F corresponded to L1014F in *M. domestica* and L995F in *An. gambiae* in domain II subunit 6 (IIS6) of the *Vgsc* gene (**Table 1**), which in *An. gambiae* species complex drastically increases IR to DDT and pyrethroids (*11, 28*). The second variant P1842S corresponded to P1874S in *An. gambiae* and P1879 in *M. domestica* and were all in the C-terminal domain (**Table 1**).

**Fig. 2:**
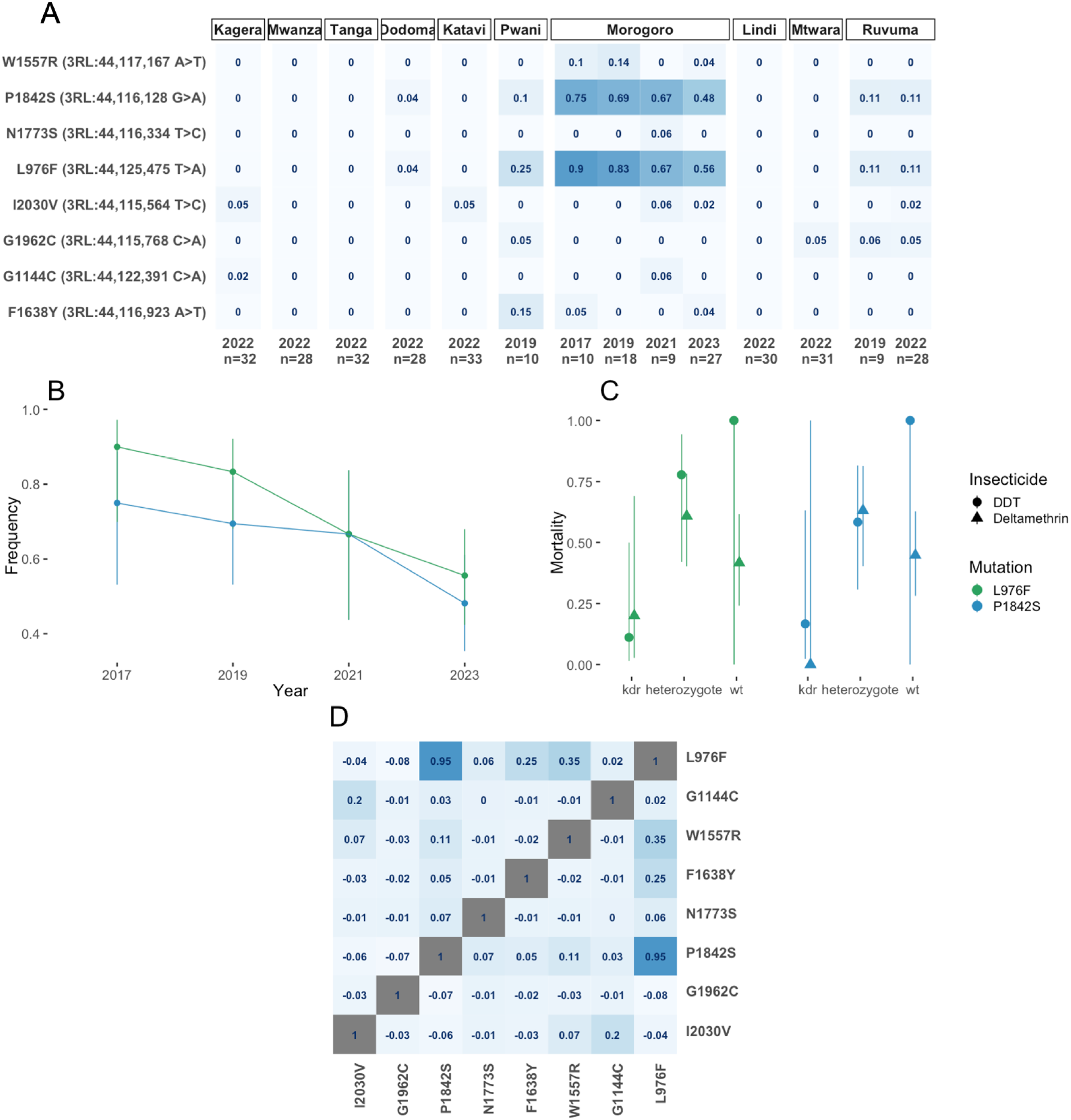
**(A)** Heatmap of *Vgsc* allele frequencies. Y-axis labels indicate mutation effect, chromosome position, and nucleotide change. X-axis labels indicate the collection date and heatmap intensity indicates frequency where darker = higher, with frequency labelled in each heatmap facet. The heatmap is panelled by the sample collection region. (**B)** L976F and 1842S frequencies, in the Morogoro region, over time. The y-axis indicates allele frequency, X-axis indicates the date. Line and point colour refer to mutation, specified in the legend. Bars indicate 95% confidence intervals. (**C**) Denotes the association of L976F and P1842S with resistance to Deltamethrin and DDT. Colour and panelling are by mutation, the x-axis indicates genotype, the y-axis indicates mortality, the point shape indicates the mean for each insecticide and the line indicates the 95% CI based on generalised mixed model prediction. (**D)** Heatmap of linkage disequilibrium (LD) (Rogers and Huff R) between nonsynonymous variants in the *Vgsc* gene at frequency > 5%. LD is indicated by fill colour. SNP effects and positions are labelled on the X and Y axes.

To explore the possible association between L976F and P1842S alleles with DDT and deltamethrin resistance, we genotyped surviving (resistant) and dead (susceptible) mosquitoes from IR bioassays for both L976Fand P1842S loci). Neither locus was associated with deltamethrin resistance: L976F (χ^2^ = 0.04, p = 0.84) and P1842S (χ^2^ = 0.59, p = 0.44). (**Fig. 2C**). We found a strong association with DDT resistance in mosquitoes carrying the mutant allele of L976F (χ^2^ = 9.23, odds ratio = 11.0, p = 0.0024) and a marginally non-significant positive association for P1842S (χ^2^ = 3.75, p = 0.0528) (**Fig. 2C**).

In *An. gambiae*, multiple *kdr* haplotypes have evolved independently (*30, 31*). To elucidate *Vgsc* haplotype structure in *An. funestus*, we computed pairwise linkage disequilibrium (LD) using the Rogers and Huff method (*32*), between nonsynonymous variants occurring at a frequency of > 5% in Tanzanian *An. funestus* **(Fig. 2C**). We found that P1824S occurred in tight LD with L976F (*D’*=0.95) **(Fig. 2D)**.

Of other non-synonymous polymorphisms, F1638Y and W1557R exhibited only weak LD with L976F (**Fig. 2D)**. We constructed a haplotype clustering dendrogram from haplotypes in all 333 individuals, from the *Vgsc* gene **(Fig. 3)**. The clustering dendrogram disclosed three major clades and three main combinations of the four most prevalent *Vgsc* alleles **(Fig. 3)**. The most striking signal was a subclade of identical, or near-identical haplotypes containing both L976F and P1842S (**Fig. 3)**, indicating a selective sweep on a combined L976F/P1842S haplotype. This combined haplotype was present at higher frequencies in the Morogoro region relative to the neighbouring regions of Pwani, Ruvuma, and Dodoma (**Fig. 3**). Most amino acid substitutions were present in a single clade in samples from Pwani, Dodoma, Ruvuma, and especially Morogoro (**Fig. 3**). This extremely restricted geographic hotspot of *kdr* is in stark contrast to its distribution in *An. gambiae* where *kdr* mutations spread rapidly across vast spatial scales in response to strong selection pressure (*26, 33*).

**Fig. 3:**
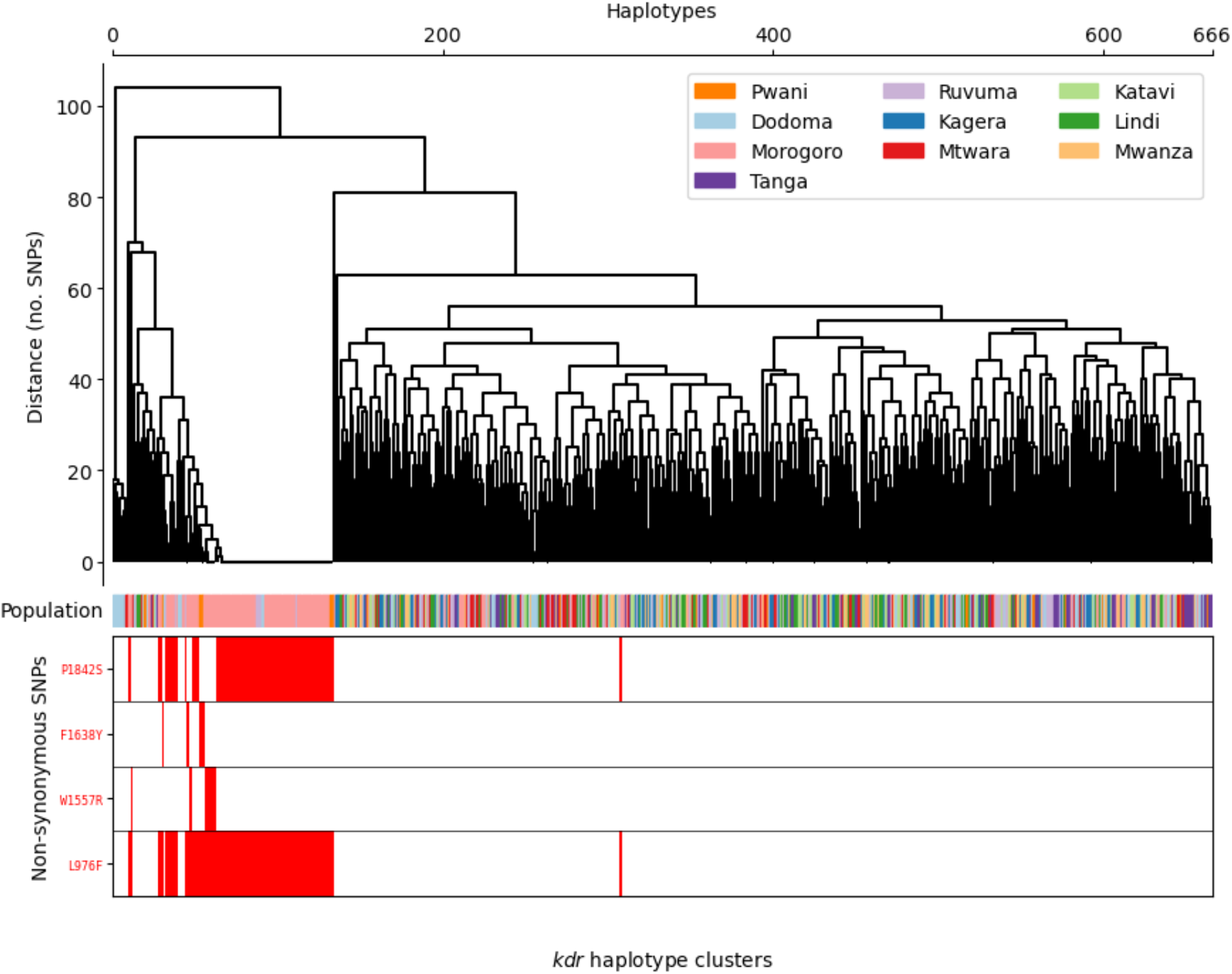
Clustering of haplotypes at the *Vgsc* gene (LOC125769886, 3RL:44105643-44156624). The dendrogram branch length corresponds to no. SNPs difference (y-axis). Tips correspond to individual haplotypes (x-axis). The coloured Population bar denotes the administrative region of origin (as described by the legend). Red blocks at the bottom indicate the presence of the labelled non-synonymous SNPs in the *Vgsc* gene.

DDT is a largely obsolete, banned, pesticide that is no longer widely used for vector control in Tanzania, or in Africa as a whole, due to its bio-accumulative and toxic properties. However, the presence of DDT resistance phenotypes in *An*.

*funestus* in Morogoro in 2015 (*34*) suggests *kdr*-mediated resistance to DDT has been present at least since then, and the emergence of *kdr* resistance to DDT suggests that future use of DDT for IRS may become even less favoured. The lack of association of *kdr* with pyrethroid resistance might be due to the strong metabolic resistance shown to pyrethroids in *An. funestus*, reducing the benefit of *kdr* (*35*). However, since our testing included only type II pyrethroids, additional data are required to assess the potential impacts of *kdr* on other pyrethroids and commercial products. The association of *kdr* with resistance to DDT but not pyrethroids, combined with selection signals and recently declining *kdr* allele frequencies where we have time series, suggests recent-past, rather than contemporary selection, perhaps due to factors other than the current use of public health pesticides.

## Discussion

In a genomic surveillance study in Tanzanian *An. funestus*, we discovered eight novel *Vgsc* mutations. Two of these, L976F and P1842S, confer *knockdown resistance* (*kdr*), occurring in tight linkage disequilibrium and at high frequencies (up to 90%) in the Morogoro region over 4 years, with limited spread to neighbouring regions. The mutation L976F showed an association with resistance to DDT, but not to pyrethroid insecticides. The role of *kdr* in pyrethroid resistance phenotypes in other *Aedes, Culex* and *Anopheles* vectors, makes the discovery of *kdr* in *An. funestus* a significant and unwelcome development that has the potential to pose a new threat to vector control in the region. Reassuringly, a lack of association between *kdr* and deltamethrin resistance indicates that the emergence of *kdr* is not linked to, nor is presently likely to threaten, the mass rollout of PBO-pyrethroid bed nets currently underway in Tanzania as a response to IR(*36*). However, this does not preclude a role for *kdr* in the *An. funestus* IR armamentarium in the future, and an urgent follow-up study is required to determine whether they confer *kdr* resistance phenotypes to other widely used pyrethroids, such as permethrin, and alpha-cypermethrin, as well as other insecticide families, especially PBO and pyrrole formulations currently being rolled out in new ITN products across the African continent (*37*).

This discovery raises intriguing questions about the conditions that have enabled the emergence of *kdr* in *An. funestus*. Our data suggests that *Vgsc* mutation in *An. funestus* do not confer target-site resistance to pyrethroids, indicating a possible explanation as to why, despite extreme selection pressures imposed by pyrethroid control have facilitated widespread propagation of resistant *Vgsc* haplotypes across the African continent in *An. gambiae (27)*, the emergence of *kdr* in Tanzanian *An. funestus* remains relatively localised. Mechanistic studies, including expression studies of mutant *Vgsc* proteins in *Xenopus* oocytes (*38*), will enable comparisons between taxa that will elucidate this further.

If the ubiquitous use of pyrethroids in vector control did not select for the emergence of *kdr*, from whence came *kdr?* Even more curiously, the apparent decline of *kdr* allele frequencies between 2017 and 2023 suggests that the selection pressure causing the emergence of *kdr* has eased (although non-uniform sample sizes per time-point make confident assertion of this difficult). There is no record of DDT use in the last decade for agriculture or vector control in the Morogoro region, or Tanzania as a whole, where the production, importation, and usage of DDT have been banned since 2009 (*17*), except for limited use in malaria vector control. In 2008, Tanzania rolled out an ambitious malaria vector control strategy relying on large-scale use of DDT for indoor residual spraying (IRS), implemented in 60 districts across the country(**Fig. 4A**), and discontinued in 2010 (*39*). Morogoro, where we detected *kdr*, was not part of this expanded campaign. Before the ban, Tanzania imported large stockpiles of DDT mostly for agricultural pest control (**Fig. 4B**).

**Fig. 4:**
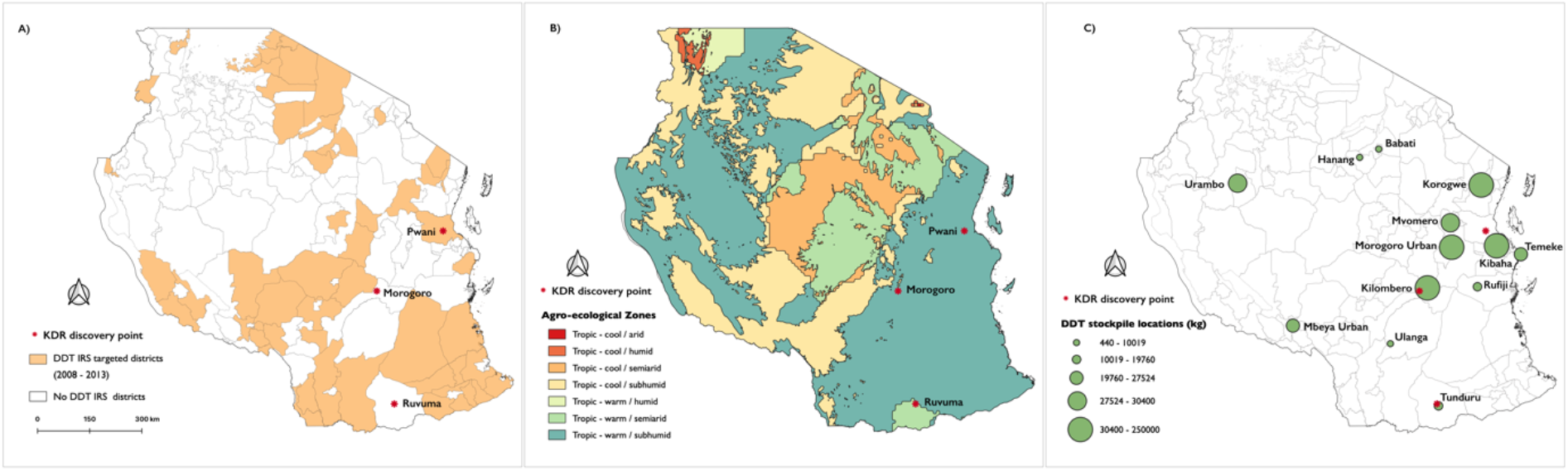
(A) Agro-ecological zones in Tanzania with colours on the map denoting the different categories indicated in the figure key. (B) Tanzanian National Malaria Control Programme (NMCP) indoor residual spraying strategy 2008 – 2012. The colours indicating districts where DDT spraying was planned. (C) DDT stockpile locations with the size of the circle indicating the stockpile quantities.

Following the ban, there have been anecdotal reports of continued illegal use of DDT amongst farmers to date (*40*). The Africa Stockpiles Programme (ASP) was launched in 2005 to eliminate stockpiles of obsolete pesticides, including DDT. At this time, it was estimated that Tanzania still possessed approx. 1,500 tonnes of obsolete pesticides (*41*), including a DDT stockpile of 30 tons (as of 2012) (*42*), approximately 50 km away from where DDT-resistant *An. funestus* were detected in this study, and 156 tons were in Morogoro town (**Fig. 4C**) (*18*). The ASP and the Tanzanian Government eliminated 100% of inventoried publicly held DDT stockpiles and conducted extensive environmental remediation by programme closed in 2013 (*43*). However, extensive DDT contamination remains (*44*), and DDT remains in widespread use by private individuals (*40*). The coincident proximal location of high levels of *kdr* in *An. funestus* with large past DDT stockpiles as well as the presence of widespread DDT contamination and private usage, leads us to hypothesise that the two most likely scenarios of *kdr* emergence in *An. funestus* are contamination of local larval breeding sites from agricultural or stockpiled DDT (**Fig. 4B, C**). The removal of DDT stockpiles by the ASP, and ongoing environmental remediation, may have contributed to reduced selection pressure on *kdr*, evident from declining frequency in Morogoro. Continued monitoring of allele frequencies and future studies of *kdr* frequencies targeted towards sites of known DDT contamination will establish whether this hypothesis is correct.

In *Silent Spring* (1962), Rachel Carson brought for the first time into the public eye the unpredictable and often remote impacts of anti-insect chemical agents on human health and nature “On one hand delicate and destructible, on the other miraculously tough and resilient, and capable of striking back in unexpected ways” (*45*). Further study of *kdr* in *An. funestus* will enable the identification of the origin of this mutation and make clear the full implications of its presence in the population for vector control. Whether the emergence of *kdr* in *An. funestus* is caused by vector control, unlicensed DDT usage in agriculture, or exposure to stockpiled DDT, our findings underscore the legacy of *Silent Spring* by reinforcing the potential for pesticides and organic pollutants to exert inadvertent influences on animal biology that may have profound and unfortunate consequences for public health.

## Materials and Methods

All scripts and Jupyter Notebooks used to analyse genotype and haplotype data, and produce figures and tables are available from the GitHub repository: https://github.com/tristanpwdennis/kdr_funestus_report_2023

### Mosquito collection

*Anopheles funestus* samples analyzed in this study were collected from ten administrative regions in Tanzania: Dodoma, Kagera, Katavi, Lindi, Morogoro, Mtwara, Mwanza, Pwani, Ruvuma, and Tanga (**Figure. 1A**). The collections were done as part of a countrywide *Anopheles funestus* surveillance project in Tanzania and were subsequently incorporated into the MalariaGEN *Anopheles funestus* genomic surveillance project database (https://www.malariagen.net/projects/anopheles-funestus-genomic-surveillance-project). Mosquitoes were collected in households between 2017 and 2023 using CDC light traps and mechanical aspirators. They were sorted by sex and taxa and *An. funestus* group mosquitoes preserved individually in 96-well plates containing 80% ethanol.

### Whole genome sequencing and analysis

The samples were processed as part of the *Anopheles funestus* genomics surveillance MalariaGEN Vector Observatory (VObs) project (https://www.malariagen.net/mosquito).Briefly, the mosquitoes were individually whole-genome-sequenced on an Illumina NovaSeq 6000s instrument. Reads were aligned to the *An. funestus* reference genome AfunGA1 (*46*) with Burrows-Wheeler Aligner (BWA) version v0.7.15. Indel realignment was performed using Genome Analysis Toolkit (GATK) version 3.7-0 RealignerTargetCreator and IndelRealigner. Single nucleotide polymorphisms were called using GATK version 3.7-0 UnifiedGenotyper. Genotypes were called for each sample independently, in genotyping mode, given all possible alleles at all genomic sites where the reference base was not “N”.

Complete specifications of the alignment and genotyping pipelines are available from the malariagen/pipelines GitHub repository (https://github.com/malariagen/pipelines/). The aligned sequences in BAM format were stored in the European Nucleotide Archive (ENA).

The identification of high-quality SNPs and haplotypes were conducted using BWA version 0.7.15 and GATK version 3.7-0. Quality control involved removal of samples with low mean coverage, removing cross-contaminated samples, running PCA to identify and remove population outliers, and sex confirmation by calling the sex of all samples based on the modal coverage ratio between the X chromosome and the autosomal chromosome arm 3R. Full quality control methods are available on the MalariaGEN vector data user guide (https://malariagen.github.io/vector-data/ag3/methods.html).

We used decision-tree filters that identify genomic sites where SNP calling and genotyping is likely to be less reliable. More information on site filters can be found on the MalariaGEN vector data user guide.

Genotypes at biallelic SNPs that passed the decision-tree site filtering process were phased into haplotypes using a combination of read-backed and statistical phasing. Read-backed phasing was performed for each sample using WhatsHap version 1.0 [https://whatshap.readthedocs.io/]. Statistical phasing was then performed using SHAPEIT4 version 4.2.1 [https://odelaneau.github.io/shapeit4/].

Complete specifications of the haplotype phasing pipeline are available from the malariagen/pipelines GitHub repository (https://github.com/malariagen/pipelines/tree/master/pipelines/phasing-vector).

### Identification of SNPs on *Vgsc*

To identify the *An. funestus Vgsc* gene and the variant that confers target-site resistance we performed alignments between the *An. gambiae* VGSC transcript AGAP004707-RD in AgamP4.12 geneset from the Ag1000 phase 3 data resource (https://www.malariagen.net/data/ag1000g-phase3-snp) and AFUN2_008728 from the *An. funestus* AfunF1.3 dataset. We extracted single nucleotide polymorphism (SNPs) altering the amino acid of VGSC protein from the *An. funestus* dataset and computed the allele frequency on the mosquito cohorts defined by the region and year of collection ((See **Supp. Table 1** for per region/year sample numbers)). Under selection pressure various alleles are expected to increase in frequency; we therefore filtered out variant alleles with a frequency lower than 5% resulting in a list of 8 variant alleles. Multiple sequence alignments of *An. funestus Vgsc* against *An. gambaie* and *M. domestica* were performed using MEGA v11.013.

### Population genetic analyses

We searched for signatures of selective sweeps on the *Vgsc* gene using the *G123* selection statistic (*47*). G123 selection scans were performed on *An. funestus* genotypes by collection region where sample *n>*20 [see **Figure 1A** and **Supp Table 2**] ] using the *g123_gwss* function in the malariagen_data python API (https://malariagen.github.io/malariagen-data-python/latest/Af1.html). Linkage disequilibrium (Rogers and Huff’s R-squared) (*32*) between the 8 *Vgsc* alleles was calculated using the *rogers_huff_r_between* in scikit-allel (https://zenodo.org/record/4759368). Haplotype clustering was performed by performing hierarchical clustering on a Hamming distance matrix, inferred from phased *An. funestus* haplotypes, using the Scipy library (https://scipy.org/citing-scipy/). Clustering dendrogram, and bar plot of amino acid substitutions, was plotted using the seaborn library (*48*).

### Association of L976F and P1842S alleles with insecticide resistance

To test for associations between the identified mutations with IR, we exposed wild non-blood-fed *An. funestus* mosquitoes of unknown ages to standard doses of deltamethrin and DDT insecticides following the WHO tube assays. For each insecticide, we randomly separated phenotypically resistant mosquitoes (i.e., alive 24 hours post-exposure) and susceptible (i.e., dead 24 hours post-exposure) and extracted DNA from individual mosquitoes using DNeasy Blood and Tissue kit (Qiagen, Germany). The mosquitoes were identified at the species level using species-specific primers that can distinguish *An. funestus* from the other members of the group (*49*). To establish if the two *kdr* variants are associated with insecticide resistance, we designed PCR primers from *An. funestus Vgsc* (Gene ID: LOC125769886) to amplify domain IIS6 (L976F) and C-terminal (P1842S) (see **Supp Table 3** for primer and thermocycler conditions). The DNA fragments were separated on a 1% agarose gel, cut, purified using PureLink™ Quick Gel Extraction Kit (Invitrogen), and commercially Sanger sequenced. Collectively, we sequenced 76 individual mosquitoes: 56 from deltamethrin and the rest from the DDT bioassays.

The frequencies of the wild type and mutant alleles were determined and correlated with phenotypes using generalised linear models in R-software v4.1.1.

## Data availability

The sequencing data generated in this study have been deposited in the European Nucleotide Archive (https://www.ebi.ac.uk/ena/browser/home) under study number PRJEB2141.

## Acknowledgements

This study was supported by funding support was received from the Bill and Melinda Gates Foundation (grant no. INV-002138) to FO, FB, HMF. Howard Hughes Medical Institute-Gates Foundation International Research Scholar Award (grant no. OPP 1099295) to FO, and the Academy Medical Science Springboard Award (ref: SBF007\100094) to FB. The findings and conclusions within this publication are those of the authors and do not necessarily reflect positions or policies of the HHMI, the BMGF or the AMSS.

This study was supported by the MalariaGEN Vector Observatory which is an international collaboration working to build capacity for malaria vector genomic research and surveillance and involves contributions by the following institutions and teams. Wellcome Sanger Institute: Lee Hart, Kelly Bennett, Anastasia Hernandez-Koutoucheva, Jon Brenas, Menelaos Ioannidis, Chris Clarkson, Alistair Miles, Julia Jeans, Paballo Chauke, Victoria Simpson, Eleanor Drury, Osama Mayet, Sónia Gonçalves, Katherine Figueroa, Tom Maddison, Kevin Howe, Mara Lawniczak; Liverpool School of Tropical Medicine: Eric Lucas, Sanjay Nagi, Martin Donnelly; Broad Institute of Harvard and MIT: Jessica Way, George Grant; Pan-African Mosquito Control Association: Jane Mwangi, Edward Lukyamuzi, Sonia Barasa, Ibra Lujumba, Elijah Juma. The authors would like to thank the staff of the Wellcome Sanger Genomic Surveillance unit and the Wellcome Sanger Institute Sample Logistics, Sequencing, and Informatics facilities for their contributions.

The MalariaGEN Vector Observatory is supported by multiple institutes and funders. The Wellcome Sanger Institute’s participation was supported by funding from Wellcome (220540/Z/20/A, ‘Wellcome Sanger Institute Quinquennial Review 2021-2026’) and the Bill & Melinda Gates Foundation (INV-001927). The Liverpool School of Tropical Medicine’s participation was supported by the National Institute of Allergy and Infectious Diseases ([NIAID] R01-AI116811), with additional support from the Medical Research Council (MR/P02520X/1). The latter grant is a UK-funded award and is part of the EDCTP2 programme supported by the European Union.

Martin Donnelly is supported by a Royal Society Wolfson Fellowship (RSWF\FT\180003). The Pan-African Mosquito Control Association’s participation was funded by the Bill and Melinda Gates Foundation (INV-031595).

The findings and conclusions within this publication are those of the authors and do not necessarily reflect the positions or policies of the funders listed above.

## Contributions

The project was conceived and supervised by FOO, FB, and DW. Field collection was performed by JOO, IHN, HB, and GM. Laboratory analysis, data acquisition and management, and preparing samples for whole genome sequencing were performed by JOO. Sequence QC, alignments, SNP calling, and haplotype phasing were performed by AHK, JN, CSC, and AM. JOO, BP, and TPWD analyzed the data and generated all figures and tables. The manuscript was drafted by JOO and TPWD and revised by all authors. Throughout the project, all authors have contributed key ideas that have shaped the work and the final paper.

---

**Supplementary Table 2:**
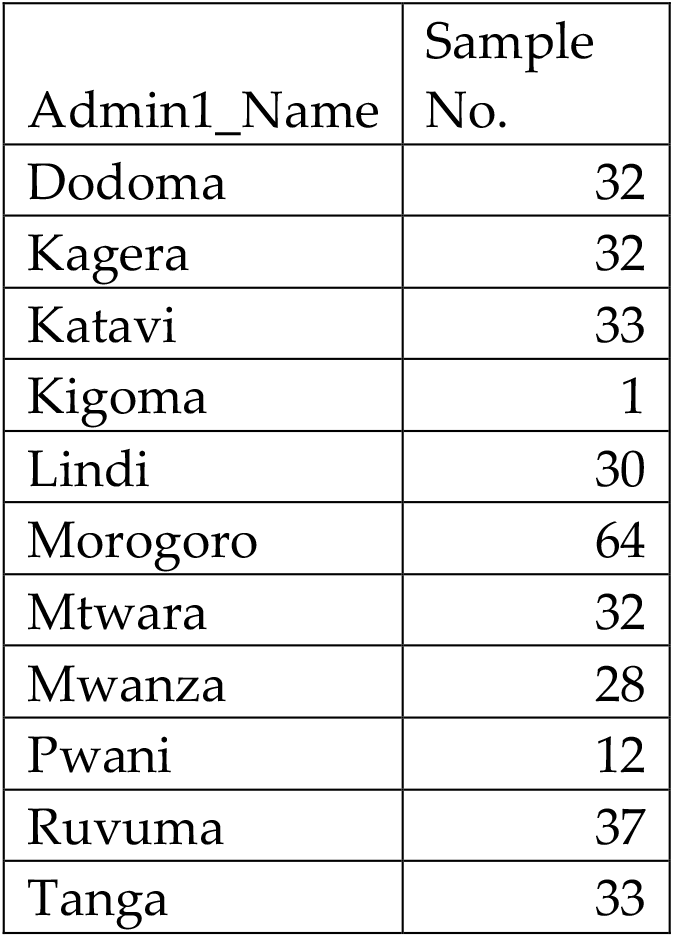
Samples location and number used in the G123 selection scans. Only locations with n>20

**Supplementary Table 3:**
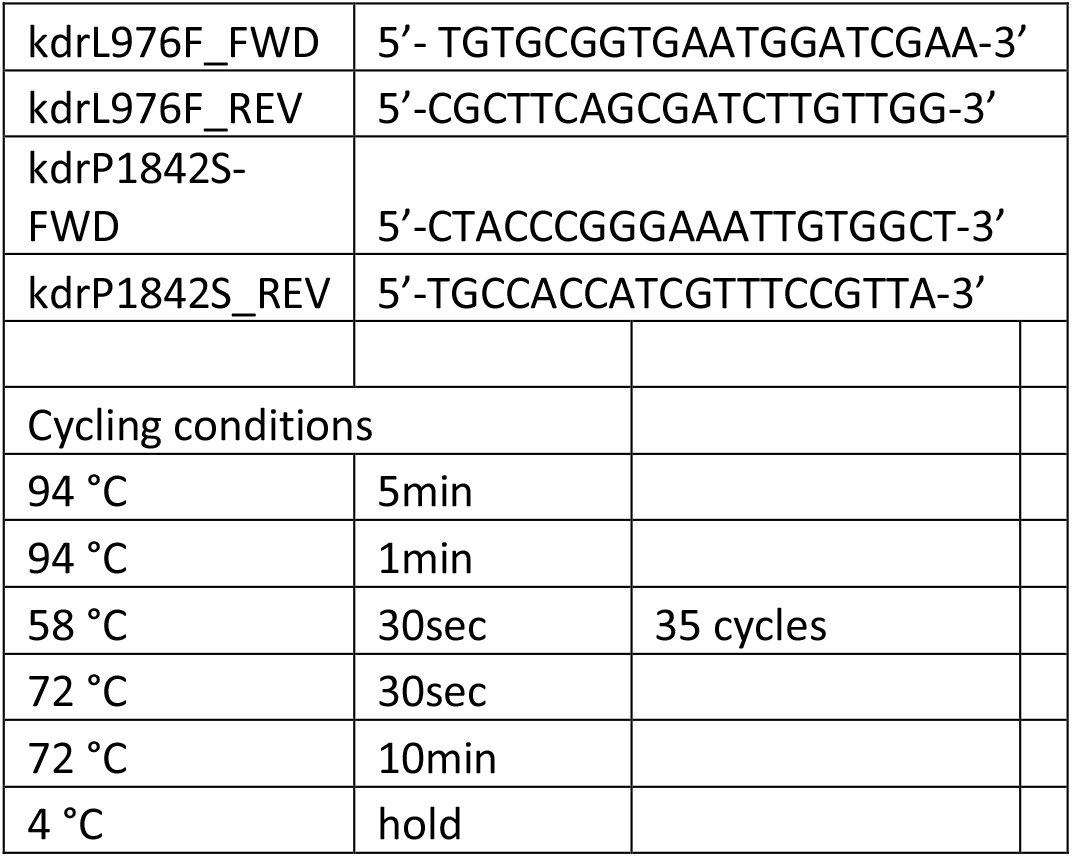
List of primer sets and cycling conditions used to amplify *Anopheles funestus* VGSC gene

