## Supplementary tables for "Discovery of knock-down resistance in the major African malaria vector *Anopheles funestus*"

**Supplementary Table 1: List of samples analysed in this study.**

| sample_id | partner_sample_id | contributor | country | location | year | month | latitude | longitude | sex_call | sample_set | release | quarter | country_iso | admin1_name | admin1_iso | admin2_name | taxon | cohort_admin1_year | cohort_admin1_month | cohort_admin1_quarter | cohort_admin2_year | cohort_admin2_month | cohort_admin2_quarter |
|---|---|---|---|---|---|---|---|---|---|---|---|---|---|---|---|---|---|---|---|---|---|---|---|
| OKFR_001_A1( | TZ-AF-L-009 | Fredros Okumu | Tanzania | Bagamoyo 4 | 2019 | 8 | -6.385 | 38.568 | F | 1236-VO-TZ-OKUMU-OKFR-TZ-2008 | 1.1 | 3 | TZA | Pwani | TZ-19 | Bagamoyo | funestus | TZ-19_fune_2019 | TZ-19_fune_2019_08 | TZ-19_fune_2019_Q3 | TZ-19_Bagamoyo_fune_2019 | TZ-19_Bagamoyo_fune_2019_08 | TZ-19_Bagamoyo_fune_2019_Q3 |
| OKFR_001_A5- | TZ-AF-L-004 | Fredros Okumu | Tanzania | Bagamoyo 4 | 2019 | 8 | -6.385 | 38.568 | F | 1236-VO-TZ-OKUMU-OKFR-TZ-2008 | 1.1 | 3 | TZA | Pwani | TZ-19 | Bagamoyo | funestus | TZ-19_fune_2019 | TZ-19_fune_2019_08 | TZ-19_fune_2019_Q3 | TZ-19_Bagamoyo_fune_2019 | TZ-19_Bagamoyo_fune_2019_08 | TZ-19_Bagamoyo_fune_2019_Q3 |
| OKFR_001_B4- | TZ-AF-A-006 | Fredros Okumu | Tanzania | Bagamoyo 1 | 2019 | 8 | -6.385 | 38.567 | F | 1236-VO-TZ-OKUMU-OKFR-TZ-2008 | 1.1 | 3 | TZA | Pwani | TZ-19 | Bagamoyo | funestus | TZ-19_fune_2019 | TZ-19_fune_2019_08 | TZ-19_fune_2019_Q3 | TZ-19_Bagamoyo_fune_2019 | TZ-19_Bagamoyo_fune_2019_08 | TZ-19_Bagamoyo_fune_2019_Q3 |
| OKFR_001_B5- | TZ-AF-A-007 | Fredros Okumu | Tanzania | Bagamoyo 1 | 2019 | 8 | -6.385 | 38.567 | F | 1236-VO-TZ-OKUMU-OKFR-TZ-2008 | 1.1 | 3 | TZA | Pwani | TZ-19 | Bagamoyo | funestus | TZ-19_fune_2019 | TZ-19_fune_2019_08 | TZ-19_fune_2019_Q3 | TZ-19_Bagamoyo_fune_2019 | TZ-19_Bagamoyo_fune_2019_08 | TZ-19_Bagamoyo_fune_2019_Q3 |
| OKFR_001_B6- | TZ-AF-A-008 | Fredros Okumu | Tanzania | Bagamoyo 1 | 2019 | 8 | -6.385 | 38.567 | F | 1236-VO-TZ-OKUMU-OKFR-TZ-2008 | 1.1 | 3 | TZA | Pwani | TZ-19 | Bagamoyo | funestus | TZ-19_fune_2019 | TZ-19_fune_2019_08 | TZ-19_fune_2019_Q3 | TZ-19_Bagamoyo_fune_2019 | TZ-19_Bagamoyo_fune_2019_08 | TZ-19_Bagamoyo_fune_2019_Q3 |
| OKFR_001_B7- | TZ-AF-A-009 | Fredros Okumu | Tanzania | Bagamoyo 1 | 2019 | 8 | -6.385 | 38.567 | F | 1236-VO-TZ-OKUMU-OKFR-TZ-2008 | 1.1 | 3 | TZA | Pwani | TZ-19 | Bagamoyo | funestus | TZ-19_fune_2019 | TZ-19_fune_2019_08 | TZ-19_fune_2019_Q3 | TZ-19_Bagamoyo_fune_2019 | TZ-19_Bagamoyo_fune_2019_08 | TZ-19_Bagamoyo_fune_2019_Q3 |
| OKFR_001_B8- | TZ-AF-A-010 | Fredros Okumu | Tanzania | Bagamoyo 1 | 2019 | 8 | -6.385 | 38.567 | F | 1236-VO-TZ-OKUMU-OKFR-TZ-2008 | 1.1 | 3 | TZA | Pwani | TZ-19 | Bagamoyo | funestus | TZ-19_fune_2019 | TZ-19_fune_2019_08 | TZ-19_fune_2019_Q3 | TZ-19_Bagamoyo_fune_2019 | TZ-19_Bagamoyo_fune_2019_08 | TZ-19_Bagamoyo_fune_2019_Q3 |
| OKFR_001_C4- | TZ-AF-A-018 | Fredros Okumu | Tanzania | Bagamoyo 2 | 2019 | 8 | -6.386 | 38.567 | F | 1236-VO-TZ-OKUMU-OKFR-TZ-2008 | 1.1 | 3 | TZA | Pwani | TZ-19 | Bagamoyo | funestus | TZ-19_fune_2019 | TZ-19_fune_2019_08 | TZ-19_fune_2019_Q3 | TZ-19_Bagamoyo_fune_2019 | TZ-19_Bagamoyo_fune_2019_08 | TZ-19_Bagamoyo_fune_2019_Q3 |
| OKFR_001_D4- | TZ-AF-A-030 | Fredros Okumu | Tanzania | Bagamoyo 3 | 2019 | 8 | -6.385 | 38.567 | F | 1236-VO-TZ-OKUMU-OKFR-TZ-2008 | 1.1 | 3 | TZA | Pwani | TZ-19 | Bagamoyo | funestus | TZ-19_fune_2019 | TZ-19_fune_2019_08 | TZ-19_fune_2019_Q3 | TZ-19_Bagamoyo_fune_2019 | TZ-19_Bagamoyo_fune_2019_08 | TZ-19_Bagamoyo_fune_2019_Q3 |
| OKFR_001_D7- | TZ-AF-A-033 | Fredros Okumu | Tanzania | Bagamoyo 3 | 2019 | 8 | -6.385 | 38.567 | F | 1236-VO-TZ-OKUMU-OKFR-TZ-2008 | 1.1 | 3 | TZA | Pwani | TZ-19 | Bagamoyo | funestus | TZ-19_fune_2019 | TZ-19_fune_2019_08 | TZ-19_fune_2019_Q3 | TZ-19_Bagamoyo_fune_2019 | TZ-19_Bagamoyo_fune_2019_08 | TZ-19_Bagamoyo_fune_2019_Q3 |
| OKFR_001_E2- | TZ-AF-A-040 | Fredros Okumu | Tanzania | Chamwino 1 | 2019 | 3 | -5.807 | 36.12 | F | 1236-VO-TZ-OKUMU-OKFR-TZ-2008 | 1.1 | 1 | TZA | Dodoma | TZ-03 | Chamwino | longipalpis | TZ-03_long_2019 | TZ-03_long_2019_03 | TZ-03_long_2019_Q1 | TZ-03_Chamwino_long_2019 | TZ-03_Chamwino_long_2019_03 | TZ-03_Chamwino_long_2019_Q1 |
| OKFR_001_E4- | TZ-AF-A-042 | Fredros Okumu | Tanzania | Chamwino 1 | 2019 | 3 | -5.807 | 36.12 | F | 1236-VO-TZ-OKUMU-OKFR-TZ-2008 | 1.1 | 1 | TZA | Dodoma | TZ-03 | Chamwino | longipalpis | TZ-03_long_2019 | TZ-03_long_2019_03 | TZ-03_long_2019_Q1 | TZ-03_Chamwino_long_2019 | TZ-03_Chamwino_long_2019_03 | TZ-03_Chamwino_long_2019_Q1 |
| OKFR_001_E7- | TZ-AF-A-045 | Fredros Okumu | Tanzania | Chamwino 2 | 2019 | 3 | -5.801 | 36.126 | F | 1236-VO-TZ-OKUMU-OKFR-TZ-2008 | 1.1 | 1 | TZA | Dodoma | TZ-03 | Chamwino | longipalpis | TZ-03_long_2019 | TZ-03_long_2019_03 | TZ-03_long_2019_Q1 | TZ-03_Chamwino_long_2019 | TZ-03_Chamwino_long_2019_03 | TZ-03_Chamwino_long_2019_Q1 |
| OKFR_001_E9- | TZ-AF-A-047 | Fredros Okumu | Tanzania | Chamwino 2 | 2019 | 3 | -5.801 | 36.126 | F | 1236-VO-TZ-OKUMU-OKFR-TZ-2008 | 1.1 | 1 | TZA | Dodoma | TZ-03 | Chamwino | longipalpis | TZ-03_long_2019 | TZ-03_long_2019_03 | TZ-03_long_2019_Q1 | TZ-03_Chamwino_long_2019 | TZ-03_Chamwino_long_2019_03 | TZ-03_Chamwino_long_2019_Q1 |
| OKFR_001_F1- | TZ-AF-L-011 | Fredros Okumu | Tanzania | Kilombero 1 | 2019 | 2 | -7.987 | 36.818 | F | 1236-VO-TZ-OKUMU-OKFR-TZ-2008 | 1.1 | 1 | TZA | Morogoro | TZ-16 | Kilombero | funestus | TZ-16_fune_2019 | TZ-16_fune_2019_02 | TZ-16_fune_2019_Q1 | TZ-16_Kilombero_fune_2019 | TZ-16_Kilombero_fune_2019_02 | TZ-16_Kilombero_fune_2019_Q1 |
| OKFR_001_F4- | TZ-AF-L-014 | Fredros Okumu | Tanzania | Kilombero 1 | 2019 | 2 | -7.987 | 36.818 | F | 1236-VO-TZ-OKUMU-OKFR-TZ-2008 | 1.1 | 1 | TZA | Morogoro | TZ-16 | Kilombero | funestus | TZ-16_fune_2019 | TZ-16_fune_2019_02 | TZ-16_fune_2019_Q1 | TZ-16_Kilombero_fune_2019 | TZ-16_Kilombero_fune_2019_02 | TZ-16_Kilombero_fune_2019_Q1 |
| OKFR_001_F5- | TZ-AF-L-015 | Fredros Okumu | Tanzania | Kilombero 1 | 2019 | 2 | -7.987 | 36.818 | F | 1236-VO-TZ-OKUMU-OKFR-TZ-2008 | 1.1 | 1 | TZA | Morogoro | TZ-16 | Kilombero | funestus | TZ-16_fune_2019 | TZ-16_fune_2019_02 | TZ-16_fune_2019_Q1 | TZ-16_Kilombero_fune_2019 | TZ-16_Kilombero_fune_2019_02 | TZ-16_Kilombero_fune_2019_Q1 |
| OKFR_001_F6- | TZ-AF-L-016 | Fredros Okumu | Tanzania | Kilombero 1 | 2019 | 2 | -7.987 | 36.818 | F | 1236-VO-TZ-OKUMU-OKFR-TZ-2008 | 1.1 | 1 | TZA | Morogoro | TZ-16 | Kilombero | funestus | TZ-16_fune_2019 | TZ-16_fune_2019_02 | TZ-16_fune_2019_Q1 | TZ-16_Kilombero_fune_2019 | TZ-16_Kilombero_fune_2019_02 | TZ-16_Kilombero_fune_2019_Q1 |
| OKFR_001_G1( | TZ-AF-A-062 | Fredros Okumu | Tanzania | Kilombero 2 | 2019 | 2 | -7.979 | 36.813 | F | 1236-VO-TZ-OKUMU-OKFR-TZ-2008 | 1.1 | 1 | TZA | Morogoro | TZ-16 | Kilombero | funestus | TZ-16_fune_2019 | TZ-16_fune_2019_02 | TZ-16_fune_2019_Q1 | TZ-16_Kilombero_fune_2019 | TZ-16_Kilombero_fune_2019_02 | TZ-16_Kilombero_fune_2019_Q1 |
| OKFR_001_G3- | TZ-AF-A-055 | Fredros Okumu | Tanzania | Kilombero 2 | 2019 | 2 | -7.979 | 36.813 | F | 1236-VO-TZ-OKUMU-OKFR-TZ-2008 | 1.1 | 1 | TZA | Morogoro | TZ-16 | Kilombero | funestus | TZ-16_fune_2019 | TZ-16_fune_2019_02 | TZ-16_fune_2019_Q1 | TZ-16_Kilombero_fune_2019 | TZ-16_Kilombero_fune_2019_02 | TZ-16_Kilombero_fune_2019_Q1 |
| OKFR_001_G5- | TZ-AF-A-057 | Fredros Okumu | Tanzania | Kilombero 2 | 2019 | 2 | -7.979 | 36.813 | F | 1236-VO-TZ-OKUMU-OKFR-TZ-2008 | 1.1 | 1 | TZA | Morogoro | TZ-16 | Kilombero | funestus | TZ-16_fune_2019 | TZ-16_fune_2019_02 | TZ-16_fune_2019_Q1 | TZ-16_Kilombero_fune_2019 | TZ-16_Kilombero_fune_2019_02 | TZ-16_Kilombero_fune_2019_Q1 |
| OKFR_001_G7- | TZ-AF-A-059 | Fredros Okumu | Tanzania | Kilombero 2 | 2019 | 2 | -7.979 | 36.813 | F | 1236-VO-TZ-OKUMU-OKFR-TZ-2008 | 1.1 | 1 | TZA | Morogoro | TZ-16 | Kilombero | funestus | TZ-16_fune_2019 | TZ-16_fune_2019_02 | TZ-16_fune_2019_Q1 | TZ-16_Kilombero_fune_2019 | TZ-16_Kilombero_fune_2019_02 | TZ-16_Kilombero_fune_2019_Q1 |
| OKFR_001_H2- | TZ-AF-A-066 | Fredros Okumu | Tanzania | Kilombero 2 | 2019 | 2 | -7.979 | 36.813 | F | 1236-VO-TZ-OKUMU-OKFR-TZ-2008 | 1.1 | 1 | TZA | Morogoro | TZ-16 | Kilombero | funestus | TZ-16_fune_2019 | TZ-16_fune_2019_02 | TZ-16_fune_2019_Q1 | TZ-16_Kilombero_fune_2019 | TZ-16_Kilombero_fune_2019_02 | TZ-16_Kilombero_fune_2019_Q1 |
| OKFR_002_A4- | TZ-AF-A-079 | Fredros Okumu | Tanzania | Kakonko | 2019 | 4 | -3.294 | 30.294 | F | 1236-VO-TZ-OKUMU-OKFR-TZ-2008 | 1.1 | 2 | TZA | Kigoma | TZ-08 | Kibondo | funestus | TZ-08_fune_2019 | TZ-08_fune_2019_04 | TZ-08_fune_2019_Q2 | TZ-08_Kibondo_fune_2019 | TZ-08_Kibondo_fune_2019_04 | TZ-08_Kibondo_fune_2019_Q2 |
| OKFR_002_B7- | TZ-AF-A-085 | Fredros Okumu | Tanzania | Muheza 2 | 2019 | 4 | -5.227 | 38.66 | F | 1236-VO-TZ-OKUMU-OKFR-TZ-2008 | 1.1 | 2 | TZA | Tanga | TZ-25 | Muheza | funestus | TZ-25_fune_2019 | TZ-25_fune_2019_04 | TZ-25_fune_2019_Q2 | TZ-25_Muheza_fune_2019 | TZ-25_Muheza_fune_2019_04 | TZ-25_Muheza_fune_2019_Q2 |
| OKFR_002_C1( | TZ-AF-A-090 | Fredros Okumu | Tanzania | Ulanga 1 | 2019 | 2 | -8.358 | 36.71 | F | 1236-VO-TZ-OKUMU-OKFR-TZ-2008 | 1.1 | 1 | TZA | Morogoro | TZ-16 | Ulanga | funestus | TZ-16_fune_2019 | TZ-16_fune_2019_02 | TZ-16_fune_2019_Q1 | TZ-16_Ulanga_fune_2019 | TZ-16_Ulanga_fune_2019_02 | TZ-16_Ulanga_fune_2019_Q1 |
| OKFR_002_C3- | TZ-AF-L-032 | Fredros Okumu | Tanzania | Ulanga 2 | 2019 | 2 | -8.354 | 36.706 | F | 1236-VO-TZ-OKUMU-OKFR-TZ-2008 | 1.1 | 1 | TZA | Morogoro | TZ-16 | Ulanga | funestus | TZ-16_fune_2019 | TZ-16_fune_2019_02 | TZ-16_fune_2019_Q1 | TZ-16_Ulanga_fune_2019 | TZ-16_Ulanga_fune_2019_02 | TZ-16_Ulanga_fune_2019_Q1 |
| OKFR_002_C9- | TZ-AF-L-038 | Fredros Okumu | Tanzania | Ulanga 2 | 2019 | 2 | -8.354 | 36.706 | F | 1236-VO-TZ-OKUMU-OKFR-TZ-2008 | 1.1 | 1 | TZA | Morogoro | TZ-16 | Ulanga | funestus | TZ-16_fune_2019 | TZ-16_fune_2019_02 | TZ-16_fune_2019_Q1 | TZ-16_Ulanga_fune_2019 | TZ-16_Ulanga_fune_2019_02 | TZ-16_Ulanga_fune_2019_Q1 |
| OKFR_002_D1( | TZ-AF-A-102 | Fredros Okumu | Tanzania | Ulanga 4 | 2019 | 2 | -8.359 | 36.71 | F | 1236-VO-TZ-OKUMU-OKFR-TZ-2008 | 1.1 | 1 | TZA | Morogoro | TZ-16 | Ulanga | funestus | TZ-16_fune_2019 | TZ-16_fune_2019_02 | TZ-16_fune_2019_Q1 | TZ-16_Ulanga_fune_2019 | TZ-16_Ulanga_fune_2019_02 | TZ-16_Ulanga_fune_2019_Q1 |
| OKFR_002_D2- | TZ-AF-A-094 | Fredros Okumu | Tanzania | Ulanga 1 | 2019 | 2 | -8.358 | 36.71 | F | 1236-VO-TZ-OKUMU-OKFR-TZ-2008 | 1.1 | 1 | TZA | Morogoro | TZ-16 | Ulanga | funestus | TZ-16_fune_2019 | TZ-16_fune_2019_02 | TZ-16_fune_2019_Q1 | TZ-16_Ulanga_fune_2019 | TZ-16_Ulanga_fune_2019_02 | TZ-16_Ulanga_fune_2019_Q1 |
| OKFR_002_D4- | TZ-AF-A-096 | Fredros Okumu | Tanzania | Ulanga 1 | 2019 | 2 | -8.358 | 36.71 | F | 1236-VO-TZ-OKUMU-OKFR-TZ-2008 | 1.1 | 1 | TZA | Morogoro | TZ-16 | Ulanga | funestus | TZ-16_fune_2019 | TZ-16_fune_2019_02 | TZ-16_fune_2019_Q1 | TZ-16_Ulanga_fune_2019 | TZ-16_Ulanga_fune_2019_02 | TZ-16_Ulanga_fune_2019_Q1 |
| OKFR_002_D7- | TZ-AF-A-099 | Fredros Okumu | Tanzania | Ulanga 3 | 2019 | 2 | -8.358 | 36.71 | F | 1236-VO-TZ-OKUMU-OKFR-TZ-2008 | 1.1 | 1 | TZA | Morogoro | TZ-16 | Ulanga | funestus | TZ-16_fune_2019 | TZ-16_fune_2019_02 | TZ-16_fune_2019_Q1 | TZ-16_Ulanga_fune_2019 | TZ-16_Ulanga_fune_2019_02 | TZ-16_Ulanga_fune_2019_Q1 |
| OKFR_002_D8- | TZ-AF-A-100 | Fredros Okumu | Tanzania | Ulanga 3 | 2019 | 2 | -8.358 | 36.71 | F | 1236-VO-TZ-OKUMU-OKFR-TZ-2008 | 1.1 | 1 | TZA | Morogoro | TZ-16 | Ulanga | funestus | TZ-16_fune_2019 | TZ-16_fune_2019_02 | TZ-16_fune_2019_Q1 | TZ-16_Ulanga_fune_2019 | TZ-16_Ulanga_fune_2019_02 | TZ-16_Ulanga_fune_2019_Q1 |
| OKFR_002_D9- | TZ-AF-A-101 | Fredros Okumu | Tanzania | Ulanga 3 | 2019 | 2 | -8.358 | 36.71 | F | 1236-VO-TZ-OKUMU-OKFR-TZ-2008 | 1.1 | 1 | TZA | Morogoro | TZ-16 | Ulanga | funestus | TZ-16_fune_2019 | TZ-16_fune_2019_02 | TZ-16_fune_2019_Q1 | TZ-16_Ulanga_fune_2019 | TZ-16_Ulanga_fune_2019_02 | TZ-16_Ulanga_fune_2019_Q1 |
| OKFR_002_E1( | TZ-AF-A-114 | Fredros Okumu | Tanzania | Tundulu 3 | 2019 | 5 | -10.957 | 37.239 | F | 1236-VO-TZ-OKUMU-OKFR-TZ-2008 | 1.1 | 2 | TZA | Ruvuma | TZ-21 | Tunduru | funestus | TZ-21_fune_2019 | TZ-21_fune_2019_05 | TZ-21_fune_2019_Q2 | TZ-21_Tunduru_fune_2019 | TZ-21_Tunduru_fune_2019_05 | TZ-21_Tunduru_fune_2019_Q2 |
| OKFR_002_E11 | TZ-AF-A-115 | Fredros Okumu | Tanzania | Tundulu 4 | 2019 | 5 | -10.958 | 37.238 | F | 1236-VO-TZ-OKUMU-OKFR-TZ-2008 | 1.1 | 2 | TZA | Ruvuma | TZ-21 | Tunduru | funestus | TZ-21_fune_2019 | TZ-21_fune_2019_05 | TZ-21_fune_2019_Q2 | TZ-21_Tunduru_fune_2019 | TZ-21_Tunduru_fune_2019_05 | TZ-21_Tunduru_fune_2019_Q2 |
| OKFR_002_E12 | TZ-AF-A-116 | Fredros Okumu | Tanzania | Tundulu 4 | 2019 | 5 | -10.958 | 37.238 | F | 1236-VO-TZ-OKUMU-OKFR-TZ-2008 | 1.1 | 2 | TZA | Ruvuma | TZ-21 | Tunduru | funestus | TZ-21_fune_2019 | TZ-21_fune_2019_05 | TZ-21_fune_2019_Q2 | TZ-21_Tunduru_fune_2019 | TZ-21_Tunduru_fune_2019_05 | TZ-21_Tunduru_fune_2019_Q2 |
| OKFR_002_E2- | TZ-AF-A-106 | Fredros Okumu | Tanzania | Tundulu 1 | 2019 | 5 | -10.959 | 37.237 | F | 1236-VO-TZ-OKUMU-OKFR-TZ-2008 | 1.1 | 2 | TZA | Ruvuma | TZ-21 | Tunduru | funestus | TZ-21_fune_2019 | TZ-21_fune_2019_05 | TZ-21_fune_2019_Q2 | TZ-21_Tunduru_fune_2019 | TZ-21_Tunduru_fune_2019_05 | TZ-21_Tunduru_fune_2019_Q2 |
| OKFR_002_E4- | TZ-AF-A-108 | Fredros Okumu | Tanzania | Tundulu 1 | 2019 | 5 | -10.959 | 37.237 | F | 1236-VO-TZ-OKUMU-OKFR-TZ-2008 | 1.1 | 2 | TZA | Ruvuma | TZ-21 | Tunduru | funestus | TZ-21_fune_2019 | TZ-21_fune_2019_05 | TZ-21_fune_2019_Q2 | TZ-21_Tunduru_fune_2019 | TZ-21_Tunduru_fune_2019_05 | TZ-21_Tunduru_fune_2019_Q2 |
| OKFR_002_E5- | TZ-AF-A-109 | Fredros Okumu | Tanzania | Tundulu 1 | 2019 | 5 | -10.959 | 37.237 | F | 1236-VO-TZ-OKUMU-OKFR-TZ-2008 | 1.1 | 2 | TZA | Ruvuma | TZ-21 | Tunduru | funestus | TZ-21_fune_2019 | TZ-21_fune_2019_05 | TZ-21_fune_2019_Q2 | TZ-21_Tunduru_fune_2019 | TZ-21_Tunduru_fune_2019_05 | TZ-21_Tunduru_fune_2019_Q2 |
| OKFR_002_E6- | TZ-AF-A-110 | Fredros Okumu | Tanzania | Tundulu 1 | 2019 | 5 | -10.959 | 37.237 | F | 1236-VO-TZ-OKUMU-OKFR-TZ-2008 | 1.1 | 2 | TZA | Ruvuma | TZ-21 | Tunduru | funestus | TZ-21_fune_2019 | TZ-21_fune_2019_05 | TZ-21_fune_2019_Q2 | TZ-21_Tunduru_fune_2019 | TZ-21_Tunduru_fune_2019_05 | TZ-21_Tunduru_fune_2019_Q2 |
| OKFR_002_F3- | TZ-AF-A-119 | Fredros Okumu | Tanzania | Tundulu 2 | 2019 | 5 | -11.071 | 37.502 | F | 1236-VO-TZ-OKUMU-OKFR-TZ-2008 | 1.1 | 2 | TZA | Ruvuma | TZ-21 | Tunduru | funestus | TZ-21_fune_2019 | TZ-21_fune_2019_05 | TZ-21_fune_2019_Q2 | TZ-21_Tunduru_fune_2019 | TZ-21_Tunduru_fune_2019_05 | TZ-21_Tunduru_fune_2019_Q2 |
| OKFR_002_F4- | TZ-AF-A-120 | Fredros Okumu | Tanzania | Tundulu 2 | 2019 | 5 | -11.071 | 37.502 | F | 1236-VO-TZ-OKUMU-OKFR-TZ-2008 | 1.1 | 2 | TZA | Ruvuma | TZ-21 | Tunduru | funestus | TZ-21_fune_2019 | TZ-21_fune_2019_05 | TZ-21_fune_2019_Q2 | TZ-21_Tunduru_fune_2019 | TZ-21_Tunduru_fune_2019_05 | TZ-21_Tunduru_fune_2019_Q2 |
| VBS23999 | 1236-TZ-A-TZF07 | Fredros Okumu | Tanzania | Minepa Village | 2017 | 8 | -8.273 | 36.681 | F | 1236-VO-TZ-OKUMU-VMF00090 | 1 | 3 | TZA | Morogoro | TZ-16 | Ulanga | funestus | TZ-16_fune_2017 | TZ-16_fune_2017_08 | TZ-16_fune_2017_Q3 | TZ-16_Ulanga_fune_2017 | TZ-16_Ulanga_fune_2017_08 | TZ-16_Ulanga_fune_2017_Q3 |
| VBS24001 | 1236-TZ-A-TZF09 | Fredros Okumu | Tanzania | Minepa Village | 2017 | 8 | -8.273 | 36.681 | F | 1236-VO-TZ-OKUMU-VMF00090 | 1 | 3 | TZA | Morogoro | TZ-16 | Ulanga | funestus | TZ-16_fune_2017 | TZ-16_fune_2017_08 | TZ-16_fune_2017_Q3 | TZ-16_Ulanga_fune_2017 | TZ-16_Ulanga_fune_2017_08 | TZ-16_Ulanga_fune_2017_Q3 |
| VBS24004 | 1236-TZ-A-TZF12 | Fredros Okumu | Tanzania | Minepa Village | 2017 | 8 | -8.273 | 36.681 | F | 1236-VO-TZ-OKUMU-VMF00090 | 1 | 3 | TZA | Morogoro | TZ-16 | Ulanga | funestus | TZ-16_fune_2017 | TZ-16_fune_2017_08 | TZ-16_fune_2017_Q3 | TZ-16_Ulanga_fune_2017 | TZ-16_Ulanga_fune_2017_08 | TZ-16_Ulanga_fune_2017_Q3 |
| VBS24005 | 1236-TZ-A-TZF13 | Fredros Okumu | Tanzania | Minepa Village | 2017 | 8 | -8.273 | 36.681 | F | 1236-VO-TZ-OKUMU-VMF00090 | 1 | 3 | TZA | Morogoro | TZ-16 | Ulanga | funestus | TZ-16_fune_2017 | TZ-16_fune_2017_08 | TZ-16_fune_2017_Q3 | TZ-16_Ulanga_fune_2017 | TZ-16_Ulanga_fune_2017_08 | TZ-16_Ulanga_fune_2017_Q3 |
| VBS24007 | 1236-TZ-A-TZF15 | Fredros Okumu | Tanzania | Minepa Village | 2017 | 8 | -8.273 | 36.681 | F | 1236-VO-TZ-OKUMU-VMF00090 | 1 | 3 | TZA | Morogoro | TZ-16 | Ulanga | funestus | TZ-16_fune_2017 | TZ-16_fune_2017_08 | TZ-16_fune_2017_Q3 | TZ-16_Ulanga_fune_2017 | TZ-16_Ulanga_fune_2017_08 | TZ-16_Ulanga_fune_2017_Q3 |
| VBS24012 | 1236-TZ-A-TZF20 | Fredros Okumu | Tanzania | Minepa Village | 2017 | 8 | -8.273 | 36.681 | F | 1236-VO-TZ-OKUMU-VMF00090 | 1 | 3 | TZA | Morogoro | TZ-16 | Ulanga | funestus | TZ-16_fune_2017 | TZ-16_fune_2017_08 | TZ-16_fune_2017_Q3 | TZ-16_Ulanga_fune_2017 | TZ-16_Ulanga_fune_2017_08 | TZ-16_Ulanga_fune_2017_Q3 |
| VBS24015 | 1236-TZ-A-TZF23 | Fredros Okumu | Tanzania | Minepa Village | 2017 | 8 | -8.273 | 36.681 | F | 1236-VO-TZ-OKUMU-VMF00090 | 1 | 3 | TZA | Morogoro | TZ-16 | Ulanga | funestus | TZ-16_fune_2017 | TZ-16_fune_2017_08 | TZ-16_fune_2017_Q3 | TZ-16_Ulanga_fune_2017 | TZ-16_Ulanga_fune_2017_08 | TZ-16_Ulanga_fune_2017_Q3 |
| VBS24020 | 1236-TZ-A-TZF28 | Fredros Okumu | Tanzania | Minepa Village | 2017 | 8 | -8.273 | 36.681 | F | 1236-VO-TZ-OKUMU-VMF00090 | 1 | 3 | TZA | Morogoro | TZ-16 | Ulanga | funestus | TZ-16_fune_2017 | TZ-16_fune_2017_08 | TZ-16_fune_2017_Q3 | TZ-16_Ulanga_fune_2017 | TZ-16_Ulanga_fune_2017_08 | TZ-16_Ulanga_fune_2017_Q3 |
| VBS24027 | 1236-TZ-A-TZF35 | Fredros Okumu | Tanzania | Minepa Village | 2017 | 8 | -8.273 | 36.681 | F | 1236-VO-TZ-OKUMU-VMF00090 | 1 | 3 | TZA | Morogoro | TZ-16 | Ulanga | funestus | TZ-16_fune_2017 | TZ-16_fune_2017_08 | TZ-16_fune_2017_Q3 | TZ-16_Ulanga_fune_2017 | TZ-16_Ulanga_fune_2017_08 | TZ-16_Ulanga_fune_2017_Q3 |
| VBS24032 | 1236-TZ-A-TZF40 | Fredros Okumu | Tanzania | Minepa Village | 2017 | 8 | -8.273 | 36.681 | F | 1236-VO-TZ-OKUMU-VMF00090 | 1 | 3 | TZA | Morogoro | TZ-16 | Ulanga | funestus | TZ-16_fune_2017 | TZ-16_fune_2017_08 | TZ-16_fune_2017_Q3 | TZ-16_Ulanga_fune_2017 | TZ-16_Ulanga_fune_2017_08 | TZ-16_Ulanga_fune_2017_Q3 |
| VBS81450 | TZ_MOR_08 | Joel Odero | Tanzania | Sululu | 2021 | -1 | -7.984 | 36.818 | F | 1236-VO-TZ-OKUMU-VMF00248 | 1.1 | -1 | TZA | Morogoro | TZ-16 | Kilombero | funestus | TZ-16_fune_2021 | TZ-16_fune_2021 | TZ-16_fune_2021 | TZ-16_Kilombero_fune_2021 | TZ-16_Kilombero_fune_2021 | TZ-16_Kilombero_fune_2021 |
| VBS81451 | TZ_KAG_01 | Joel Odero | Tanzania | Byeju | 2022 | -1 | -1.08 | 31.376 | F | 1236-VO-TZ-OKUMU-VMF00248 | 1.1 | -1 | TZA | Kagera | TZ-05 | Missenyi | funestus | TZ-05_fune_2022 | TZ-05_fune_2022 | TZ-05_fune_2022 | TZ-05_Missenyi_fune_2022 | TZ-05_Missenyi_fune_2022 | TZ-05_Missenyi_fune_2022 |
| VBS81452 | TZ_KAG_02 | Joel Odero | Tanzania | Byeju | 2022 | -1 | -1.08 | 31.376 | F | 1236-VO-TZ-OKUMU-VMF00248 | 1.1 | -1 | TZA | Kagera | TZ-05 | Missenyi | funestus | TZ-05_fune_2022 | TZ-05_fune_2022 | TZ-05_fune_2022 | TZ-05_Missenyi_fune_2022 | TZ-05_Missenyi_fune_2022 | TZ-05_Missenyi_fune_2022 |
| VBS81453 | TZ_KAG_09 | Joel Odero | Tanzania | Byeju | 2022 | -1 | -1.08 | 31.376 | F | 1236-VO-TZ-OKUMU-VMF00248 | 1.1 | -1 | TZA | Kagera | TZ-05 | Missenyi | funestus | TZ-05_fune_2022 | TZ-05_fune_2022 | TZ-05_fune_2022 | TZ-05_Missenyi_fune_2022 | TZ-05_Missenyi_fune_2022 | TZ-05_Missenyi_fune_2022 |
| VBS81454 | TZ_KAG_10 | Joel Odero | Tanzania | Byeju | 2022 | -1 | -1.08 | 31.376 | F | 1236-VO-TZ-OKUMU-VMF00248 | 1.1 | -1 | TZA | Kagera | TZ-05 | Missenyi | funestus | TZ-05_fune_2022 | TZ-05_fune_2022 | TZ-05_fune_2022 | TZ-05_Missenyi_fune_2022 | TZ-05_Missenyi_fune_2022 | TZ-05_Missenyi_fune_2022 |
| VBS81455 | TZ_MTW_02 | Joel Odero | Tanzania | Mahurunga | 2022 | -1 | -10.538 | 40.279 | F | 1236-VO-TZ-OKUMU-VMF00248 | 1.1 | -1 | TZA | Mtwara | TZ-17 | Mtwara | funestus | TZ-17_fune_2022 | TZ-17_fune_2022 | TZ-17_fune_2022 | TZ-17_Mtwara_fune_2022 | TZ-17_Mtwara_fune_2022 | TZ-17_Mtwara_fune_2022 |
| VBS81456 | TZ_MTW_09 | Joel Odero | Tanzania | Mahurunga | 2022 | -1 | -10.538 | 40.279 | F | 1236-VO-TZ-OKUMU-VMF00248 | 1.1 | -1 | TZA | Mtwara | TZ-17 | Mtwara | funestus | TZ-17_fune_2022 | TZ-17_fune_2022 | TZ-17_fune_2022 | TZ-17_Mtwara_fune_2022 | TZ-17_Mtwara_fune_2022 | TZ-17_Mtwara_fune_2022 |
| VBS81457 | TZ_MOR_11 | Joel Odero | Tanzania | Sululu | 2021 | -1 | -7.984 | 36.818 | F | 1236-VO-TZ-OKUMU-VMF00248 | 1.1 | -1 | TZA | Morogoro | TZ-16 | Kilombero | funestus | TZ-16_fune_2021 | TZ-16_fune_2021 | TZ-16_fune_2021 | TZ-16_Kilombero_fune_2021 | TZ-16_Kilombero_fune_2021 | TZ-16_Kilombero_fune_2021 |
| VBS81458 | TZ_MOR_12 | Joel Odero | Tanzania | Sululu | 2021 | -1 | -7.984 | 36.818 | F | 1236-VO-TZ-OKUMU-VMF00248 | 1.1 | -1 | TZA | Morogoro | TZ-16 | Kilombero | funestus | TZ-16_fune_2021 | TZ-16_fune_2021 | TZ-16_fune_2021 | TZ-16_Kilombero_fune_2021 | TZ-16_Kilombero_fune_2021 | TZ-16_Kilombero_fune_2021 |
| VBS81459 | TZ_MOR_13 | Joel Odero | Tanzania | Sululu | 2021 | -1 | -7.984 | 36.818 | F | 1236-VO-TZ-OKUMU-VMF00248 | 1.1 | -1 | TZA | Morogoro | TZ-16 | Kilombero | funestus | TZ-16_fune_2021 | TZ-16_fune_2021 | TZ-16_fune_2021 | TZ-16_Kilombero_fune_2021 | TZ-16_Kilombero_fune_2021 | TZ-16_Kilombero_fune_2021 |
| VBS81460 | TZ_MOR_14 | Joel Odero | Tanzania | Sululu | 2021 | -1 | -7.984 | 36.818 | F | 1236-VO-TZ-OKUMU-VMF00248 | 1.1 | -1 | TZA | Morogoro | TZ-16 | Kilombero | funestus | TZ-16_fune_2021 | TZ-16_fune_2021 | TZ-16_fune_2021 | TZ-16_Kilombero_fune_2021 | TZ-16_Kilombero_fune_2021 | TZ-16_Kilombero_fune_2021 |
| VBS81461 | TZ_MOR_15 | Joel Odero | Tanzania | Sululu | 2021 | -1 | -7.984 | 36.818 | F | 1236-VO-TZ-OKUMU-VMF00248 | 1.1 | -1 | TZA | Morogoro | TZ-16 | Kilombero | funestus | TZ-16_fune_2021 | TZ-16_fune_2021 | TZ-16_fune_2021 | TZ-16_Kilombero_fune_2021 | TZ-16_Kilombero_fune_2021 | TZ-16_Kilombero_fune_2021 |
| VBS81462 | TZ_MOR_16 | Joel Odero | Tanzania | Sululu | 2021 | -1 | -7.984 | 36.818 | F | 1236-VO-TZ-OKUMU-VMF00248 | 1.1 | -1 | TZA | Morogoro | TZ-16 | Kilombero | funestus | TZ-16_fune_2021 | TZ-16_fune_2021 | TZ-16_fune_2021 | TZ-16_Kilombero_fune_2021 | TZ-16_Kilombero_fune_2021 | TZ-16_Kilombero_fune_2021 |
| VBS81463 | TZ_MOR_18 | Joel Odero | Tanzania | Sululu | 2021 | -1 | -7.984 | 36.818 | F | 1236-VO-TZ-OKUMU-VMF00248 | 1.1 | -1 | TZA | Morogoro | TZ-16 | Kilombero | funestus | TZ-16_fune_2021 | TZ-16_fune_2021 | TZ-16_fune_2021 | TZ-16_Kilombero_fune_2021 | TZ-16_Kilombero_fune_2021 | TZ-16_Kilombero_fune_2021 |
| VBS81464 | TZ_MOR_19 | Joel Odero | Tanzania | Sululu | 2021 | -1 | -7.984 | 36.818 | F | 1236-VO-TZ-OKUMU-VMF00248 | 1.1 | -1 | TZA | Morogoro | TZ-16 | Kilombero | funestus | TZ-16_fune_2021 | TZ-16_fune_2021 | TZ-16_fune_2021 | TZ-16_Kilombero_fune_2021 | TZ-16_Kilombero_fune_2021 | TZ-16_Kilombero_fune_2021 |
| VBS81466 | TZ_KAG_21 | Joel Odero | Tanzania | Byeju | 2022 | -1 | -1.08 | 31.376 | F | 1236-VO-TZ-OKUMU-VMF00248 | 1.1 | -1 | TZA | Kagera | TZ-05 | Missenyi | funestus | TZ-05_fune_2022 | TZ-05_fune_2022 | TZ-05_fune_2022 | TZ-05_Missenyi_fune_2022 | TZ-05_Missenyi_fune_2022 | TZ-05_Missenyi_fune_2022 |
| VBS81467 | TZ_KAG_22 | Joel Odero | Tanzania | Byeju | 2022 | -1 | -1.08 | 31.376 | F | 1236-VO-TZ-OKUMU-VMF00248 | 1.1 | -1 | TZA | Kagera | TZ-05 | Missenyi | funestus | TZ-05_fune_2022 | TZ-05_fune_2022 | TZ-05_fune_2022 | TZ-05_Missenyi_fune_2022 | TZ-05_Missenyi_fune_2022 | TZ-05_Missenyi_fune_2022 |
| VBS81468 | TZ_KAG_23 | Joel Odero | Tanzania | Byeju | 2022 | -1 | -1.08 | 31.376 | F | 1236-VO-TZ-OKUMU-VMF00248 | 1.1 | -1 | TZA | Kagera | TZ-05 | Missenyi | funestus | TZ-05_fune_2022 | TZ-05_fune_2022 | TZ-05_fune_2022 | TZ-05_Missenyi_fune_2022 | TZ-05_Missenyi_fune_2022 | TZ-05_Missenyi_fune_2022 |
| VBS81469 | TZ_KAG_24 | Joel Odero | Tanzania | Byeju | 2022 | -1 | -1.08 | 31.376 | F | 1236-VO-TZ-OKUMU-VMF00248 | 1.1 | -1 | TZA | Kagera | TZ-05 | Missenyi | funestus | TZ-05_fune_2022 | TZ-05_fune_2022 | TZ-05_fune_2022 | TZ-05_Missenyi_fune_2022 | TZ-05_Missenyi_fune_2022 | TZ-05_Missenyi_fune_2022 |
| VBS81470 | TZ_KAG_26 | Joel Odero | Tanzania | Byeju | 2022 | -1 | -1.08 | 31.376 | F | 1236-VO-TZ-OKUMU-VMF00248 | 1.1 | -1 | TZA | Kagera | TZ-05 | Missenyi | funestus | TZ-05_fune_2022 | TZ-05_fune_2022 | TZ-05_fune_2022 | TZ-05_Missenyi_fune_2022 | TZ-05_Missenyi_fune_2022 | TZ-05_Missenyi_fune_2022 |
| VBS81471 | TZ_KAG_29 | Joel Odero | Tanzania | Byeju | 2022 | -1 | -1.08 | 31.376 | F | 1236-VO-TZ-OKUMU-VMF00248 | 1.1 | -1 | TZA | Kagera | TZ-05 | Missenyi | funestus | TZ-05_fune_2022 | TZ-05_fune_2022 | TZ-05_fune_2022 | TZ-05_Missenyi_fune_2022 | TZ-05_Missenyi_fune_2022 | TZ-05_Missenyi_fune_2022 |
| VBS81472 | TZ_MTW_11 | Joel Odero | Tanzania | Mahurunga | 2022 | -1 | -10.538 | 40.279 | F | 1236-VO-TZ-OKUMU-VMF00248 | 1.1 | -1 | TZA | Mtwara | TZ-17 | Mtwara | funestus | TZ-17_fune_2022 | TZ-17_fune_2022 | TZ-17_fune_2022 | TZ-17_Mtwara_fune_2022 | TZ-17_Mtwara_fune_2022 | TZ-17_Mtwara_fune_2022 |
| VBS81473 | TZ_MTW_14 | Joel Odero | Tanzania | Mahurunga | 2022 | -1 | -10.538 | 40.279 | F | 1236-VO-TZ-OKUMU-VMF00248 | 1.1 | -1 | TZA | Mtwara | TZ-17 | Mtwara | funestus | TZ-17_fune_2022 | TZ-17_fune_2022 | TZ-17_fune_2022 | TZ-17_Mtwara_fune_2022 | TZ-17_Mtwara_fune_2022 | TZ-17_Mtwara_fune_2022 |
| VBS81474 | TZ_MTW_15 | Joel Odero | Tanzania | Mahurunga | 2022 | -1 | -10.538 | 40.279 | F | 1236-VO-TZ-OKUMU-VMF00248 | 1.1 | -1 | TZA | Mtwara | TZ-17 | Mtwara | funestus | TZ-17_fune_2022 | TZ-17_fune_2022 | TZ-17_fune_2022 | TZ-17_Mtwara_fune_2022 | TZ-17_Mtwara_fune_2022 | TZ-17_Mtwara_fune_2022 |

| | | | | | | | | | | | | | | | | | | | | | | | |
|---|---|---|---|---|---|---|---|---|---|---|---|---|---|---|---|---|---|---|---|---|---|---|---|
| VBS81475 | TZ_MTW_16 | Joel Odero | Tanzania | Mahurunga | 2022 | -1 | -10.538 | 40.279 | F | 1236-VO-TZ-OKUMU-VMF00248 | 1.1 | -1 | TZA | Mtwara | TZ-17 | Mtwara | funestus | TZ-17_fune_2022 | TZ-17_fune_2022 | TZ-17_fune_2022 | TZ-17_Mtwara_fune_2022 | TZ-17_Mtwara_fune_2022 | TZ-17_Mtwara_fune_2022 |
| VBS81476 | TZ_MTW_17 | Joel Odero | Tanzania | Mahurunga | 2022 | -1 | -10.538 | 40.279 | F | 1236-VO-TZ-OKUMU-VMF00248 | 1.1 | -1 | TZA | Mtwara | TZ-17 | Mtwara | funestus | TZ-17_fune_2022 | TZ-17_fune_2022 | TZ-17_fune_2022 | TZ-17_Mtwara_fune_2022 | TZ-17_Mtwara_fune_2022 | TZ-17_Mtwara_fune_2022 |
| VBS81477 | TZ_MTW_18 | Joel Odero | Tanzania | Mahurunga | 2022 | -1 | -10.538 | 40.279 | F | 1236-VO-TZ-OKUMU-VMF00248 | 1.1 | -1 | TZA | Mtwara | TZ-17 | Mtwara | parensis | TZ-17_pare_2022 | TZ-17_pare_2022 | TZ-17_pare_2022 | TZ-17_Mtwara_pare_2022 | TZ-17_Mtwara_pare_2022 | TZ-17_Mtwara_pare_2022 |
| VBS81478 | TZ_MTW_19 | Joel Odero | Tanzania | Mahurunga | 2022 | -1 | -10.538 | 40.279 | F | 1236-VO-TZ-OKUMU-VMF00248 | 1.1 | -1 | TZA | Mtwara | TZ-17 | Mtwara | funestus | TZ-17_fune_2022 | TZ-17_fune_2022 | TZ-17_fune_2022 | TZ-17_Mtwara_fune_2022 | TZ-17_Mtwara_fune_2022 | TZ-17_Mtwara_fune_2022 |
| VBS81479 | TZ_MTW_20 | Joel Odero | Tanzania | Mahurunga | 2022 | -1 | -10.538 | 40.279 | F | 1236-VO-TZ-OKUMU-VMF00248 | 1.1 | -1 | TZA | Mtwara | TZ-17 | Mtwara | funestus | TZ-17_fune_2022 | TZ-17_fune_2022 | TZ-17_fune_2022 | TZ-17_Mtwara_fune_2022 | TZ-17_Mtwara_fune_2022 | TZ-17_Mtwara_fune_2022 |
| VBS82166 | TZ_KAT_2 | Joel Odero | Tanzania | Vikonge | 2022 | 3 | -6.091 | 30.936 | F | 1236-VO-TZ-OKUMU-VMF00252 | 1.2 | 1 | TZA | Katavi | TZ-28 | Mpanda | funestus | TZ-28_fune_2022 | TZ-28_fune_2022_03 | TZ-28_fune_2022_Q1 | TZ-28_Mpanda_fune_2022 | TZ-28_Mpanda_fune_2022_03 | TZ-28_Mpanda_fune_2022_Q1 |
| VBS82167 | TZ_KAT_3 | Joel Odero | Tanzania | Vikonge | 2022 | 3 | -6.091 | 30.936 | F | 1236-VO-TZ-OKUMU-VMF00252 | 1.2 | 1 | TZA | Katavi | TZ-28 | Mpanda | funestus | TZ-28_fune_2022 | TZ-28_fune_2022_03 | TZ-28_fune_2022_Q1 | TZ-28_Mpanda_fune_2022 | TZ-28_Mpanda_fune_2022_03 | TZ-28_Mpanda_fune_2022_Q1 |
| VBS82168 | TZ_KAT_5 | Joel Odero | Tanzania | Vikonge | 2022 | 3 | -6.091 | 30.936 | F | 1236-VO-TZ-OKUMU-VMF00252 | 1.2 | 1 | TZA | Katavi | TZ-28 | Mpanda | funestus | TZ-28_fune_2022 | TZ-28_fune_2022_03 | TZ-28_fune_2022_Q1 | TZ-28_Mpanda_fune_2022 | TZ-28_Mpanda_fune_2022_03 | TZ-28_Mpanda_fune_2022_Q1 |
| VBS82169 | TZ_KAT_8 | Joel Odero | Tanzania | Vikonge | 2022 | 3 | -6.091 | 30.936 | F | 1236-VO-TZ-OKUMU-VMF00252 | 1.2 | 1 | TZA | Katavi | TZ-28 | Mpanda | funestus | TZ-28_fune_2022 | TZ-28_fune_2022_03 | TZ-28_fune_2022_Q1 | TZ-28_Mpanda_fune_2022 | TZ-28_Mpanda_fune_2022_03 | TZ-28_Mpanda_fune_2022_Q1 |
| VBS82170 | TZ_KAT_9 | Joel Odero | Tanzania | Vikonge | 2022 | 3 | -6.091 | 30.936 | F | 1236-VO-TZ-OKUMU-VMF00252 | 1.2 | 1 | TZA | Katavi | TZ-28 | Mpanda | funestus | TZ-28_fune_2022 | TZ-28_fune_2022_03 | TZ-28_fune_2022_Q1 | TZ-28_Mpanda_fune_2022 | TZ-28_Mpanda_fune_2022_03 | TZ-28_Mpanda_fune_2022_Q1 |
| VBS82171 | TZ_KAT_11 | Joel Odero | Tanzania | Vikonge | 2022 | 3 | -6.091 | 30.936 | F | 1236-VO-TZ-OKUMU-VMF00252 | 1.2 | 1 | TZA | Katavi | TZ-28 | Mpanda | funestus | TZ-28_fune_2022 | TZ-28_fune_2022_03 | TZ-28_fune_2022_Q1 | TZ-28_Mpanda_fune_2022 | TZ-28_Mpanda_fune_2022_03 | TZ-28_Mpanda_fune_2022_Q1 |
| VBS82172 | TZ_KAT_14 | Joel Odero | Tanzania | Vikonge | 2022 | 3 | -6.091 | 30.936 | F | 1236-VO-TZ-OKUMU-VMF00252 | 1.2 | 1 | TZA | Katavi | TZ-28 | Mpanda | funestus | TZ-28_fune_2022 | TZ-28_fune_2022_03 | TZ-28_fune_2022_Q1 | TZ-28_Mpanda_fune_2022 | TZ-28_Mpanda_fune_2022_03 | TZ-28_Mpanda_fune_2022_Q1 |
| VBS82173 | TZ_KAT_15 | Joel Odero | Tanzania | Vikonge | 2022 | 3 | -6.091 | 30.936 | F | 1236-VO-TZ-OKUMU-VMF00252 | 1.2 | 1 | TZA | Katavi | TZ-28 | Mpanda | funestus | TZ-28_fune_2022 | TZ-28_fune_2022_03 | TZ-28_fune_2022_Q1 | TZ-28_Mpanda_fune_2022 | TZ-28_Mpanda_fune_2022_03 | TZ-28_Mpanda_fune_2022_Q1 |
| VBS82174 | TZ_KAT_19 | Joel Odero | Tanzania | Vikonge | 2022 | 3 | -6.091 | 30.936 | F | 1236-VO-TZ-OKUMU-VMF00252 | 1.2 | 1 | TZA | Katavi | TZ-28 | Mpanda | funestus | TZ-28_fune_2022 | TZ-28_fune_2022_03 | TZ-28_fune_2022_Q1 | TZ-28_Mpanda_fune_2022 | TZ-28_Mpanda_fune_2022_03 | TZ-28_Mpanda_fune_2022_Q1 |
| VBS82176 | TZ_RUV_8 | Joel Odero | Tanzania | Nandembo | 2022 | 3 | -10.983 | 37.283 | F | 1236-VO-TZ-OKUMU-VMF00252 | 1.2 | 1 | TZA | Ruvuma | TZ-21 | Tunduru | funestus | TZ-21_fune_2022 | TZ-21_fune_2022_03 | TZ-21_fune_2022_Q1 | TZ-21_Tunduru_fune_2022 | TZ-21_Tunduru_fune_2022_03 | TZ-21_Tunduru_fune_2022_Q1 |
| VBS82177 | TZ_RUV_9 | Joel Odero | Tanzania | Nandembo | 2022 | 3 | -10.983 | 37.283 | F | 1236-VO-TZ-OKUMU-VMF00252 | 1.2 | 1 | TZA | Ruvuma | TZ-21 | Tunduru | funestus | TZ-21_fune_2022 | TZ-21_fune_2022_03 | TZ-21_fune_2022_Q1 | TZ-21_Tunduru_fune_2022 | TZ-21_Tunduru_fune_2022_03 | TZ-21_Tunduru_fune_2022_Q1 |
| VBS82178 | TZ_RUV_13 | Joel Odero | Tanzania | Nandembo | 2022 | 3 | -10.983 | 37.283 | F | 1236-VO-TZ-OKUMU-VMF00252 | 1.2 | 1 | TZA | Ruvuma | TZ-21 | Tunduru | funestus | TZ-21_fune_2022 | TZ-21_fune_2022_03 | TZ-21_fune_2022_Q1 | TZ-21_Tunduru_fune_2022 | TZ-21_Tunduru_fune_2022_03 | TZ-21_Tunduru_fune_2022_Q1 |
| VBS82179 | TZ_RUV_18 | Joel Odero | Tanzania | Nandembo | 2022 | 3 | -10.983 | 37.283 | F | 1236-VO-TZ-OKUMU-VMF00252 | 1.2 | 1 | TZA | Ruvuma | TZ-21 | Tunduru | funestus | TZ-21_fune_2022 | TZ-21_fune_2022_03 | TZ-21_fune_2022_Q1 | TZ-21_Tunduru_fune_2022 | TZ-21_Tunduru_fune_2022_03 | TZ-21_Tunduru_fune_2022_Q1 |
| VBS82180 | TZ_DOM_1 | Joel Odero | Tanzania | Mbori | 2022 | 5 | -6.272 | 36.378 | F | 1236-VO-TZ-OKUMU-VMF00252 | 1.2 | 2 | TZA | Dodoma | TZ-03 | Mpwapwa | funestus | TZ-03_fune_2022 | TZ-03_fune_2022_05 | TZ-03_fune_2022_Q2 | TZ-03_Mpwapwa_fune_2022 | TZ-03_Mpwapwa_fune_2022_05 | TZ-03_Mpwapwa_fune_2022_Q2 |
| VBS82181 | TZ_DOM_2 | Joel Odero | Tanzania | Mbori | 2022 | 5 | -6.272 | 36.378 | F | 1236-VO-TZ-OKUMU-VMF00252 | 1.2 | 2 | TZA | Dodoma | TZ-03 | Mpwapwa | funestus | TZ-03_fune_2022 | TZ-03_fune_2022_05 | TZ-03_fune_2022_Q2 | TZ-03_Mpwapwa_fune_2022 | TZ-03_Mpwapwa_fune_2022_05 | TZ-03_Mpwapwa_fune_2022_Q2 |
| VBS82182 | TZ_DOM_3 | Joel Odero | Tanzania | Mbori | 2022 | 5 | -6.272 | 36.378 | F | 1236-VO-TZ-OKUMU-VMF00252 | 1.2 | 2 | TZA | Dodoma | TZ-03 | Mpwapwa | funestus | TZ-03_fune_2022 | TZ-03_fune_2022_05 | TZ-03_fune_2022_Q2 | TZ-03_Mpwapwa_fune_2022 | TZ-03_Mpwapwa_fune_2022_05 | TZ-03_Mpwapwa_fune_2022_Q2 |
| VBS82183 | TZ_DOM_4 | Joel Odero | Tanzania | Mbori | 2022 | 5 | -6.272 | 36.378 | F | 1236-VO-TZ-OKUMU-VMF00252 | 1.2 | 2 | TZA | Dodoma | TZ-03 | Mpwapwa | funestus | TZ-03_fune_2022 | TZ-03_fune_2022_05 | TZ-03_fune_2022_Q2 | TZ-03_Mpwapwa_fune_2022 | TZ-03_Mpwapwa_fune_2022_05 | TZ-03_Mpwapwa_fune_2022_Q2 |
| VBS82185 | TZ_DOM_6 | Joel Odero | Tanzania | Mbori | 2022 | 5 | -6.272 | 36.378 | F | 1236-VO-TZ-OKUMU-VMF00252 | 1.2 | 2 | TZA | Dodoma | TZ-03 | Mpwapwa | funestus | TZ-03_fune_2022 | TZ-03_fune_2022_05 | TZ-03_fune_2022_Q2 | TZ-03_Mpwapwa_fune_2022 | TZ-03_Mpwapwa_fune_2022_05 | TZ-03_Mpwapwa_fune_2022_Q2 |
| VBS82187 | TZ_DOM_8 | Joel Odero | Tanzania | Mbori | 2022 | 5 | -6.272 | 36.378 | F | 1236-VO-TZ-OKUMU-VMF00252 | 1.2 | 2 | TZA | Dodoma | TZ-03 | Mpwapwa | funestus | TZ-03_fune_2022 | TZ-03_fune_2022_05 | TZ-03_fune_2022_Q2 | TZ-03_Mpwapwa_fune_2022 | TZ-03_Mpwapwa_fune_2022_05 | TZ-03_Mpwapwa_fune_2022_Q2 |
| VBS82189 | TZ_DOM_11 | Joel Odero | Tanzania | Mbori | 2022 | 5 | -6.272 | 36.378 | F | 1236-VO-TZ-OKUMU-VMF00252 | 1.2 | 2 | TZA | Dodoma | TZ-03 | Mpwapwa | funestus | TZ-03_fune_2022 | TZ-03_fune_2022_05 | TZ-03_fune_2022_Q2 | TZ-03_Mpwapwa_fune_2022 | TZ-03_Mpwapwa_fune_2022_05 | TZ-03_Mpwapwa_fune_2022_Q2 |
| VBS82190 | TZ_DOM_12 | Joel Odero | Tanzania | Mbori | 2022 | 5 | -6.272 | 36.378 | F | 1236-VO-TZ-OKUMU-VMF00252 | 1.2 | 2 | TZA | Dodoma | TZ-03 | Mpwapwa | funestus | TZ-03_fune_2022 | TZ-03_fune_2022_05 | TZ-03_fune_2022_Q2 | TZ-03_Mpwapwa_fune_2022 | TZ-03_Mpwapwa_fune_2022_05 | TZ-03_Mpwapwa_fune_2022_Q2 |
| VBS82191 | TZ_DOM_13 | Joel Odero | Tanzania | Mbori | 2022 | 5 | -6.272 | 36.378 | F | 1236-VO-TZ-OKUMU-VMF00252 | 1.2 | 2 | TZA | Dodoma | TZ-03 | Mpwapwa | funestus | TZ-03_fune_2022 | TZ-03_fune_2022_05 | TZ-03_fune_2022_Q2 | TZ-03_Mpwapwa_fune_2022 | TZ-03_Mpwapwa_fune_2022_05 | TZ-03_Mpwapwa_fune_2022_Q2 |
| VBS82192 | TZ_DOM_14 | Joel Odero | Tanzania | Mbori | 2022 | 5 | -6.272 | 36.378 | F | 1236-VO-TZ-OKUMU-VMF00252 | 1.2 | 2 | TZA | Dodoma | TZ-03 | Mpwapwa | funestus | TZ-03_fune_2022 | TZ-03_fune_2022_05 | TZ-03_fune_2022_Q2 | TZ-03_Mpwapwa_fune_2022 | TZ-03_Mpwapwa_fune_2022_05 | TZ-03_Mpwapwa_fune_2022_Q2 |
| VBS82193 | TZ_DOM_15 | Joel Odero | Tanzania | Mbori | 2022 | 5 | -6.272 | 36.378 | F | 1236-VO-TZ-OKUMU-VMF00252 | 1.2 | 2 | TZA | Dodoma | TZ-03 | Mpwapwa | funestus | TZ-03_fune_2022 | TZ-03_fune_2022_05 | TZ-03_fune_2022_Q2 | TZ-03_Mpwapwa_fune_2022 | TZ-03_Mpwapwa_fune_2022_05 | TZ-03_Mpwapwa_fune_2022_Q2 |
| VBS82195 | TZ_DOM_17 | Joel Odero | Tanzania | Mbori | 2022 | 5 | -6.272 | 36.378 | F | 1236-VO-TZ-OKUMU-VMF00252 | 1.2 | 2 | TZA | Dodoma | TZ-03 | Mpwapwa | funestus | TZ-03_fune_2022 | TZ-03_fune_2022_05 | TZ-03_fune_2022_Q2 | TZ-03_Mpwapwa_fune_2022 | TZ-03_Mpwapwa_fune_2022_05 | TZ-03_Mpwapwa_fune_2022_Q2 |
| VBS82196 | TZ_DOM_18 | Joel Odero | Tanzania | Mbori | 2022 | 5 | -6.272 | 36.378 | F | 1236-VO-TZ-OKUMU-VMF00252 | 1.2 | 2 | TZA | Dodoma | TZ-03 | Mpwapwa | funestus | TZ-03_fune_2022 | TZ-03_fune_2022_05 | TZ-03_fune_2022_Q2 | TZ-03_Mpwapwa_fune_2022 | TZ-03_Mpwapwa_fune_2022_05 | TZ-03_Mpwapwa_fune_2022_Q2 |
| VBS82197 | TZ_DOM_19 | Joel Odero | Tanzania | Mbori | 2022 | 5 | -6.272 | 36.378 | F | 1236-VO-TZ-OKUMU-VMF00252 | 1.2 | 2 | TZA | Dodoma | TZ-03 | Mpwapwa | funestus | TZ-03_fune_2022 | TZ-03_fune_2022_05 | TZ-03_fune_2022_Q2 | TZ-03_Mpwapwa_fune_2022 | TZ-03_Mpwapwa_fune_2022_05 | TZ-03_Mpwapwa_fune_2022_Q2 |
| VBS82198 | TZ_DOM_20 | Joel Odero | Tanzania | Mbori | 2022 | 5 | -6.272 | 36.378 | F | 1236-VO-TZ-OKUMU-VMF00252 | 1.2 | 2 | TZA | Dodoma | TZ-03 | Mpwapwa | funestus | TZ-03_fune_2022 | TZ-03_fune_2022_05 | TZ-03_fune_2022_Q2 | TZ-03_Mpwapwa_fune_2022 | TZ-03_Mpwapwa_fune_2022_05 | TZ-03_Mpwapwa_fune_2022_Q2 |
| VBS82199 | TZ_BAG_1 | Joel Odero | Tanzania | Sengasenga | 2022 | 8 | -6.386 | 38.575 | F | 1236-VO-TZ-OKUMU-VMF00252 | 1.2 | 3 | TZA | Pwani | TZ-19 | Bagamoyo | funestus | TZ-19_fune_2022 | TZ-19_fune_2022_08 | TZ-19_fune_2022_Q3 | TZ-19_Bagamoyo_fune_2022 | TZ-19_Bagamoyo_fune_2022_08 | TZ-19_Bagamoyo_fune_2022_Q3 |
| VBS82200 | TZ_BAG_4 | Joel Odero | Tanzania | Sengasenga | 2022 | 8 | -6.386 | 38.575 | F | 1236-VO-TZ-OKUMU-VMF00252 | 1.2 | 3 | TZA | Pwani | TZ-19 | Bagamoyo | funestus | TZ-19_fune_2022 | TZ-19_fune_2022_08 | TZ-19_fune_2022_Q3 | TZ-19_Bagamoyo_fune_2022 | TZ-19_Bagamoyo_fune_2022_08 | TZ-19_Bagamoyo_fune_2022_Q3 |
| VBS82201 | TZ_LND_1 | Joel Odero | Tanzania | Mtualonga | 2022 | 9 | -10.218 | 39.492 | F | 1236-VO-TZ-OKUMU-VMF00252 | 1.2 | 3 | TZA | Lindi | TZ-12 | Lindi | funestus | TZ-12_fune_2022 | TZ-12_fune_2022_09 | TZ-12_fune_2022_Q3 | TZ-12_Lindi_fune_2022 | TZ-12_Lindi_fune_2022_09 | TZ-12_Lindi_fune_2022_Q3 |
| VBS82202 | TZ_LND_2 | Joel Odero | Tanzania | Mtualonga | 2022 | 9 | -10.218 | 39.492 | F | 1236-VO-TZ-OKUMU-VMF00252 | 1.2 | 3 | TZA | Lindi | TZ-12 | Lindi | funestus | TZ-12_fune_2022 | TZ-12_fune_2022_09 | TZ-12_fune_2022_Q3 | TZ-12_Lindi_fune_2022 | TZ-12_Lindi_fune_2022_09 | TZ-12_Lindi_fune_2022_Q3 |
| VBS82203 | TZ_LND_3 | Joel Odero | Tanzania | Mtualonga | 2022 | 9 | -10.218 | 39.492 | F | 1236-VO-TZ-OKUMU-VMF00252 | 1.2 | 3 | TZA | Lindi | TZ-12 | Lindi | funestus | TZ-12_fune_2022 | TZ-12_fune_2022_09 | TZ-12_fune_2022_Q3 | TZ-12_Lindi_fune_2022 | TZ-12_Lindi_fune_2022_09 | TZ-12_Lindi_fune_2022_Q3 |
| VBS82204 | TZ_LND_4 | Joel Odero | Tanzania | Mtualonga | 2022 | 9 | -10.218 | 39.492 | F | 1236-VO-TZ-OKUMU-VMF00252 | 1.2 | 3 | TZA | Lindi | TZ-12 | Lindi | funestus | TZ-12_fune_2022 | TZ-12_fune_2022_09 | TZ-12_fune_2022_Q3 | TZ-12_Lindi_fune_2022 | TZ-12_Lindi_fune_2022_09 | TZ-12_Lindi_fune_2022_Q3 |
| VBS82205 | TZ_LND_5 | Joel Odero | Tanzania | Mtualonga | 2022 | 9 | -10.218 | 39.492 | F | 1236-VO-TZ-OKUMU-VMF00252 | 1.2 | 3 | TZA | Lindi | TZ-12 | Lindi | funestus | TZ-12_fune_2022 | TZ-12_fune_2022_09 | TZ-12_fune_2022_Q3 | TZ-12_Lindi_fune_2022 | TZ-12_Lindi_fune_2022_09 | TZ-12_Lindi_fune_2022_Q3 |
| VBS82206 | TZ_LND_6 | Joel Odero | Tanzania | Mtualonga | 2022 | 9 | -10.218 | 39.492 | F | 1236-VO-TZ-OKUMU-VMF00252 | 1.2 | 3 | TZA | Lindi | TZ-12 | Lindi | funestus | TZ-12_fune_2022 | TZ-12_fune_2022_09 | TZ-12_fune_2022_Q3 | TZ-12_Lindi_fune_2022 | TZ-12_Lindi_fune_2022_09 | TZ-12_Lindi_fune_2022_Q3 |
| VBS82207 | TZ_LND_7 | Joel Odero | Tanzania | Mtualonga | 2022 | 9 | -10.218 | 39.492 | F | 1236-VO-TZ-OKUMU-VMF00252 | 1.2 | 3 | TZA | Lindi | TZ-12 | Lindi | funestus | TZ-12_fune_2022 | TZ-12_fune_2022_09 | TZ-12_fune_2022_Q3 | TZ-12_Lindi_fune_2022 | TZ-12_Lindi_fune_2022_09 | TZ-12_Lindi_fune_2022_Q3 |
| VBS82208 | TZ_LND_8 | Joel Odero | Tanzania | Mtualonga | 2022 | 9 | -10.218 | 39.492 | F | 1236-VO-TZ-OKUMU-VMF00252 | 1.2 | 3 | TZA | Lindi | TZ-12 | Lindi | funestus | TZ-12_fune_2022 | TZ-12_fune_2022_09 | TZ-12_fune_2022_Q3 | TZ-12_Lindi_fune_2022 | TZ-12_Lindi_fune_2022_09 | TZ-12_Lindi_fune_2022_Q3 |
| VBS82209 | TZ_LND_10 | Joel Odero | Tanzania | Mtualonga | 2022 | 9 | -10.218 | 39.492 | F | 1236-VO-TZ-OKUMU-VMF00252 | 1.2 | 3 | TZA | Lindi | TZ-12 | Lindi | funestus | TZ-12_fune_2022 | TZ-12_fune_2022_09 | TZ-12_fune_2022_Q3 | TZ-12_Lindi_fune_2022 | TZ-12_Lindi_fune_2022_09 | TZ-12_Lindi_fune_2022_Q3 |
| VBS82210 | TZ_LND_11 | Joel Odero | Tanzania | Mtualonga | 2022 | 9 | -10.218 | 39.492 | F | 1236-VO-TZ-OKUMU-VMF00252 | 1.2 | 3 | TZA | Lindi | TZ-12 | Lindi | funestus | TZ-12_fune_2022 | TZ-12_fune_2022_09 | TZ-12_fune_2022_Q3 | TZ-12_Lindi_fune_2022 | TZ-12_Lindi_fune_2022_09 | TZ-12_Lindi_fune_2022_Q3 |
| VBS82211 | TZ_LND_12 | Joel Odero | Tanzania | Mtualonga | 2022 | 9 | -10.218 | 39.492 | F | 1236-VO-TZ-OKUMU-VMF00252 | 1.2 | 3 | TZA | Lindi | TZ-12 | Lindi | funestus | TZ-12_fune_2022 | TZ-12_fune_2022_09 | TZ-12_fune_2022_Q3 | TZ-12_Lindi_fune_2022 | TZ-12_Lindi_fune_2022_09 | TZ-12_Lindi_fune_2022_Q3 |
| VBS82212 | TZ_LND_13 | Joel Odero | Tanzania | Mtualonga | 2022 | 9 | -10.218 | 39.492 | F | 1236-VO-TZ-OKUMU-VMF00252 | 1.2 | 3 | TZA | Lindi | TZ-12 | Lindi | funestus | TZ-12_fune_2022 | TZ-12_fune_2022_09 | TZ-12_fune_2022_Q3 | TZ-12_Lindi_fune_2022 | TZ-12_Lindi_fune_2022_09 | TZ-12_Lindi_fune_2022_Q3 |
| VBS82213 | TZ_LND_14 | Joel Odero | Tanzania | Mtualonga | 2022 | 9 | -10.218 | 39.492 | F | 1236-VO-TZ-OKUMU-VMF00252 | 1.2 | 3 | TZA | Lindi | TZ-12 | Lindi | funestus | TZ-12_fune_2022 | TZ-12_fune_2022_09 | TZ-12_fune_2022_Q3 | TZ-12_Lindi_fune_2022 | TZ-12_Lindi_fune_2022_09 | TZ-12_Lindi_fune_2022_Q3 |
| VBS82214 | TZ_LND_15 | Joel Odero | Tanzania | Mtualonga | 2022 | 9 | -10.218 | 39.492 | F | 1236-VO-TZ-OKUMU-VMF00252 | 1.2 | 3 | TZA | Lindi | TZ-12 | Lindi | funestus | TZ-12_fune_2022 | TZ-12_fune_2022_09 | TZ-12_fune_2022_Q3 | TZ-12_Lindi_fune_2022 | TZ-12_Lindi_fune_2022_09 | TZ-12_Lindi_fune_2022_Q3 |
| VBS82215 | TZ_LND_16 | Joel Odero | Tanzania | Mtualonga | 2022 | 9 | -10.218 | 39.492 | F | 1236-VO-TZ-OKUMU-VMF00252 | 1.2 | 3 | TZA | Lindi | TZ-12 | Lindi | funestus | TZ-12_fune_2022 | TZ-12_fune_2022_09 | TZ-12_fune_2022_Q3 | TZ-12_Lindi_fune_2022 | TZ-12_Lindi_fune_2022_09 | TZ-12_Lindi_fune_2022_Q3 |
| VBS82216 | TZ_LND_19 | Joel Odero | Tanzania | Mtualonga | 2022 | 9 | -10.218 | 39.492 | F | 1236-VO-TZ-OKUMU-VMF00252 | 1.2 | 3 | TZA | Lindi | TZ-12 | Lindi | funestus | TZ-12_fune_2022 | TZ-12_fune_2022_09 | TZ-12_fune_2022_Q3 | TZ-12_Lindi_fune_2022 | TZ-12_Lindi_fune_2022_09 | TZ-12_Lindi_fune_2022_Q3 |
| VBS82217 | TZ_TNG_1 | Joel Odero | Tanzania | Kwamwale | 2022 | 5 | -5.157 | 38.729 | F | 1236-VO-TZ-OKUMU-VMF00252 | 1.2 | 2 | TZA | Tanga | TZ-25 | Muheza | funestus | TZ-25_fune_2022 | TZ-25_fune_2022_05 | TZ-25_fune_2022_Q2 | TZ-25_Muheza_fune_2022 | TZ-25_Muheza_fune_2022_05 | TZ-25_Muheza_fune_2022_Q2 |
| VBS82218 | TZ_TNG_5 | Joel Odero | Tanzania | Kwamwale | 2022 | 5 | -5.157 | 38.729 | F | 1236-VO-TZ-OKUMU-VMF00252 | 1.2 | 2 | TZA | Tanga | TZ-25 | Muheza | funestus | TZ-25_fune_2022 | TZ-25_fune_2022_05 | TZ-25_fune_2022_Q2 | TZ-25_Muheza_fune_2022 | TZ-25_Muheza_fune_2022_05 | TZ-25_Muheza_fune_2022_Q2 |
| VBS82220 | TZ_TNG_8 | Joel Odero | Tanzania | Kwamwale | 2022 | 5 | -5.157 | 38.729 | F | 1236-VO-TZ-OKUMU-VMF00252 | 1.2 | 2 | TZA | Tanga | TZ-25 | Muheza | funestus | TZ-25_fune_2022 | TZ-25_fune_2022_05 | TZ-25_fune_2022_Q2 | TZ-25_Muheza_fune_2022 | TZ-25_Muheza_fune_2022_05 | TZ-25_Muheza_fune_2022_Q2 |
| VBS82221 | TZ_TNG_9 | Joel Odero | Tanzania | Kwamwale | 2022 | 5 | -5.157 | 38.729 | F | 1236-VO-TZ-OKUMU-VMF00252 | 1.2 | 2 | TZA | Tanga | TZ-25 | Muheza | funestus | TZ-25_fune_2022 | TZ-25_fune_2022_05 | TZ-25_fune_2022_Q2 | TZ-25_Muheza_fune_2022 | TZ-25_Muheza_fune_2022_05 | TZ-25_Muheza_fune_2022_Q2 |
| VBS82222 | TZ_TNG_10 | Joel Odero | Tanzania | Kwamwale | 2022 | 5 | -5.157 | 38.729 | F | 1236-VO-TZ-OKUMU-VMF00252 | 1.2 | 2 | TZA | Tanga | TZ-25 | Muheza | funestus | TZ-25_fune_2022 | TZ-25_fune_2022_05 | TZ-25_fune_2022_Q2 | TZ-25_Muheza_fune_2022 | TZ-25_Muheza_fune_2022_05 | TZ-25_Muheza_fune_2022_Q2 |
| VBS82223 | TZ_TNG_11 | Joel Odero | Tanzania | Kwamwale | 2022 | 5 | -5.157 | 38.729 | F | 1236-VO-TZ-OKUMU-VMF00252 | 1.2 | 2 | TZA | Tanga | TZ-25 | Muheza | funestus | TZ-25_fune_2022 | TZ-25_fune_2022_05 | TZ-25_fune_2022_Q2 | TZ-25_Muheza_fune_2022 | TZ-25_Muheza_fune_2022_05 | TZ-25_Muheza_fune_2022_Q2 |
| VBS82224 | TZ_TNG_12 | Joel Odero | Tanzania | Kwamwale | 2022 | 5 | -5.157 | 38.729 | F | 1236-VO-TZ-OKUMU-VMF00252 | 1.2 | 2 | TZA | Tanga | TZ-25 | Muheza | funestus | TZ-25_fune_2022 | TZ-25_fune_2022_05 | TZ-25_fune_2022_Q2 | TZ-25_Muheza_fune_2022 | TZ-25_Muheza_fune_2022_05 | TZ-25_Muheza_fune_2022_Q2 |
| VBS82225 | TZ_TNG_13 | Joel Odero | Tanzania | Kwamwale | 2022 | 5 | -5.157 | 38.729 | F | 1236-VO-TZ-OKUMU-VMF00252 | 1.2 | 2 | TZA | Tanga | TZ-25 | Muheza | funestus | TZ-25_fune_2022 | TZ-25_fune_2022_05 | TZ-25_fune_2022_Q2 | TZ-25_Muheza_fune_2022 | TZ-25_Muheza_fune_2022_05 | TZ-25_Muheza_fune_2022_Q2 |
| VBS82226 | TZ_TNG_14 | Joel Odero | Tanzania | Kwamwale | 2022 | 5 | -5.157 | 38.729 | F | 1236-VO-TZ-OKUMU-VMF00252 | 1.2 | 2 | TZA | Tanga | TZ-25 | Muheza | funestus | TZ-25_fune_2022 | TZ-25_fune_2022_05 | TZ-25_fune_2022_Q2 | TZ-25_Muheza_fune_2022 | TZ-25_Muheza_fune_2022_05 | TZ-25_Muheza_fune_2022_Q2 |
| VBS82227 | TZ_TNG_15 | Joel Odero | Tanzania | Kwamwale | 2022 | 5 | -5.157 | 38.729 | F | 1236-VO-TZ-OKUMU-VMF00252 | 1.2 | 2 | TZA | Tanga | TZ-25 | Muheza | funestus | TZ-25_fune_2022 | TZ-25_fune_2022_05 | TZ-25_fune_2022_Q2 | TZ-25_Muheza_fune_2022 | TZ-25_Muheza_fune_2022_05 | TZ-25_Muheza_fune_2022_Q2 |
| VBS82228 | TZ_TNG_16 | Joel Odero | Tanzania | Kwamwale | 2022 | 5 | -5.157 | 38.729 | F | 1236-VO-TZ-OKUMU-VMF00252 | 1.2 | 2 | TZA | Tanga | TZ-25 | Muheza | funestus | TZ-25_fune_2022 | TZ-25_fune_2022_05 | TZ-25_fune_2022_Q2 | TZ-25_Muheza_fune_2022 | TZ-25_Muheza_fune_2022_05 | TZ-25_Muheza_fune_2022_Q2 |
| VBS82229 | TZ_TNG_17 | Joel Odero | Tanzania | Kwamwale | 2022 | 5 | -5.157 | 38.729 | F | 1236-VO-TZ-OKUMU-VMF00252 | 1.2 | 2 | TZA | Tanga | TZ-25 | Muheza | funestus | TZ-25_fune_2022 | TZ-25_fune_2022_05 | TZ-25_fune_2022_Q2 | TZ-25_Muheza_fune_2022 | TZ-25_Muheza_fune_2022_05 | TZ-25_Muheza_fune_2022_Q2 |
| VBS82230 | TZ_TNG_18 | Joel Odero | Tanzania | Kwamwale | 2022 | 5 | -5.157 | 38.729 | F | 1236-VO-TZ-OKUMU-VMF00252 | 1.2 | 2 | TZA | Tanga | TZ-25 | Muheza | funestus | TZ-25_fune_2022 | TZ-25_fune_2022_05 | TZ-25_fune_2022_Q2 | TZ-25_Muheza_fune_2022 | TZ-25_Muheza_fune_2022_05 | TZ-25_Muheza_fune_2022_Q2 |
| VBS85333 | TZ_KAT_36 | Joel Odero | Tanzania | Vikonge | 2022 | 3 | -6.091 | 30.936 | F | 1236-VO-TZ-OKUMU-VMF00261 | 1.3 | 1 | TZA | Katavi | TZ-28 | Mpanda | funestus | TZ-28_fune_2022 | TZ-28_fune_2022_03 | TZ-28_fune_2022_Q1 | TZ-28_Mpanda_fune_2022 | TZ-28_Mpanda_fune_2022_03 | TZ-28_Mpanda_fune_2022_Q1 |
| VBS85334 | TZ_KAT_35 | Joel Odero | Tanzania | Vikonge | 2022 | 3 | -6.091 | 30.936 | F | 1236-VO-TZ-OKUMU-VMF00261 | 1.3 | 1 | TZA | Katavi | TZ-28 | Mpanda | funestus | TZ-28_fune_2022 | TZ-28_fune_2022_03 | TZ-28_fune_2022_Q1 | TZ-28_Mpanda_fune_2022 | TZ-28_Mpanda_fune_2022_03 | TZ-28_Mpanda_fune_2022_Q1 |
| VBS85335 | TZ_KAT_34 | Joel Odero | Tanzania | Vikonge | 2022 | 3 | -6.091 | 30.936 | F | 1236-VO-TZ-OKUMU-VMF00261 | 1.3 | 1 | TZA | Katavi | TZ-28 | Mpanda | funestus | TZ-28_fune_2022 | TZ-28_fune_2022_03 | TZ-28_fune_2022_Q1 | TZ-28_Mpanda_fune_2022 | TZ-28_Mpanda_fune_2022_03 | TZ-28_Mpanda_fune_2022_Q1 |
| VBS85336 | TZ_KAT_33 | Joel Odero | Tanzania | Vikonge | 2022 | 3 | -6.091 | 30.936 | F | 1236-VO-TZ-OKUMU-VMF00261 | 1.3 | 1 | TZA | Katavi | TZ-28 | Mpanda | funestus | TZ-28_fune_2022 | TZ-28_fune_2022_03 | TZ-28_fune_2022_Q1 | TZ-28_Mpanda_fune_2022 | TZ-28_Mpanda_fune_2022_03 | TZ-28_Mpanda_fune_2022_Q1 |
| VBS85337 | TZ_KAT_32 | Joel Odero | Tanzania | Vikonge | 2022 | 3 | -6.091 | 30.936 | F | 1236-VO-TZ-OKUMU-VMF00261 | 1.3 | 1 | TZA | Katavi | TZ-28 | Mpanda | funestus | TZ-28_fune_2022 | TZ-28_fune_2022_03 | TZ-28_fune_2022_Q1 | TZ-28_Mpanda_fune_2022 | TZ-28_Mpanda_fune_2022_03 | TZ-28_Mpanda_fune_2022_Q1 |
| VBS85338 | TZ_KAT_31 | Joel Odero | Tanzania | Vikonge | 2022 | 3 | -6.091 | 30.936 | F | 1236-VO-TZ-OKUMU-VMF00261 | 1.3 | 1 | TZA | Katavi | TZ-28 | Mpanda | funestus | TZ-28_fune_2022 | TZ-28_fune_2022_03 | TZ-28_fune_2022_Q1 | TZ-28_Mpanda_fune_2022 | TZ-28_Mpanda_fune_2022_03 | TZ-28_Mpanda_fune_2022_Q1 |
| VBS85339 | TZ_LND_34 | Joel Odero | Tanzania | Mtualonga | 2022 | 9 | -10.218 | 39.492 | F | 1236-VO-TZ-OKUMU-VMF00261 | 1.3 | 3 | TZA | Lindi | TZ-12 | Lindi | funestus | TZ-12_fune_2022 | TZ-12_fune_2022_09 | TZ-12_fune_2022_Q3 | TZ-12_Lindi_fune_2022 | TZ-12_Lindi_fune_2022_09 | TZ-12_Lindi_fune_2022_Q3 |
| VBS85340 | TZ_LND_33 | Joel Odero | Tanzania | Mtualonga | 2022 | 9 | -10.218 | 39.492 | F | 1236-VO-TZ-OKUMU-VMF00261 | 1.3 | 3 | TZA | Lindi | TZ-12 | Lindi | funestus | TZ-12_fune_2022 | TZ-12_fune_2022_09 | TZ-12_fune_2022_Q3 | TZ-12_Lindi_fune_2022 | TZ-12_Lindi_fune_2022_09 | TZ-12_Lindi_fune_2022_Q3 |
| VBS85341 | TZ_LND_32 | Joel Odero | Tanzania | Mtualonga | 2022 | 9 | -10.218 | 39.492 | F | 1236-VO-TZ-OKUMU-VMF00261 | 1.3 | 3 | TZA | Lindi | TZ-12 | Lindi | funestus | TZ-12_fune_2022 | TZ-12_fune_2022_09 | TZ-12_fune_2022_Q3 | TZ-12_Lindi_fune_2022 | TZ-12_Lindi_fune_2022_09 | TZ-12_Lindi_fune_2022_Q3 |
| VBS85342 | TZ_LND_31 | Joel Odero | Tanzania | Mtualonga | 2022 | 9 | -10.218 | 39.492 | F | 1236-VO-TZ-OKUMU-VMF00261 | 1.3 | 3 | TZA | Lindi | TZ-12 | Lindi | funestus | TZ-12_fune_2022 | TZ-12_fune_2022_09 | TZ-12_fune_2022_Q3 | TZ-12_Lindi_fune_2022 | TZ-12_Lindi_fune_2022_09 | TZ-12_Lindi_fune_2022_Q3 |
| VBS85343 | TZ_LND_30 | Joel Odero | Tanzania | Mtualonga | 2022 | 9 | -10.218 | 39.492 | F | 1236-VO-TZ-OKUMU-VMF00261 | 1.3 | 3 | TZA | Lindi | TZ-12 | Lindi | funestus | TZ-12_fune_2022 | TZ-12_fune_2022_09 | TZ-12_fune_2022_Q3 | TZ-12_Lindi_fune_2022 | TZ-12_Lindi_fune_2022_09 | TZ-12_Lindi_fune_2022_Q3 |
| VBS85344 | TZ_LND_29 | Joel Odero | Tanzania | Mtualonga | 2022 | 9 | -10.218 | 39.492 | F | 1236-VO-TZ-OKUMU-VMF00261 | 1.3 | 3 | TZA | Lindi | TZ-12 | Lindi | funestus | TZ-12_fune_2022 | TZ-12_fune_2022_09 | TZ-12_fune_2022_Q3 | TZ-12_Lindi_fune_2022 | TZ-12_Lindi_fune_2022_09 | TZ-12_Lindi_fune_2022_Q3 |
| VBS85345 | TZ_LND_28 | Joel Odero | Tanzania | Mtualonga | 2022 | 9 | -10.218 | 39.492 | F | 1236-VO-TZ-OKUMU-VMF00261 | 1.3 | 3 | TZA | Lindi | TZ-12 | Lindi | funestus | TZ-12_fune_2022 | TZ-12_fune_2022_09 | TZ-12_fune_2022_Q3 | TZ-12_Lindi_fune_2022 | TZ-12_Lindi_fune_2022_09 | TZ-12_Lindi_fune_2022_Q3 |
| VBS85346 | TZ_LND_27 | Joel Odero | Tanzania | Mtualonga | 2022 | 9 | -10.218 | 39.492 | F | 1236-VO-TZ-OKUMU-VMF00261 | 1.3 | 3 | TZA | Lindi | TZ-12 | Lindi | funestus | TZ-12_fune_2022 | TZ-12_fune_2022_09 | TZ-12_fune_2022_Q3 | TZ-12_Lindi_fune_2022 | TZ-12_Lindi_fune_2022_09 | TZ-12_Lindi_fune_2022_Q3 |
| VBS85347 | TZ_LND_26 | Joel Odero | Tanzania | Mtualonga | 2022 | 9 | -10.218 | 39.492 | F | 1236-VO-TZ-OKUMU-VMF00261 | 1.3 | 3 | TZA | Lindi | TZ-12 | Lindi | funestus | TZ-12_fune_2022 | TZ-12_fune_2022_09 | TZ-12_fune_2022_Q3 | TZ-12_Lindi_fune_2022 | TZ-12_Lindi_fune_2022_09 | TZ-12_Lindi_fune_2022_Q3 |
| VBS85348 | TZ_KAG_42 | Joel Odero | Tanzania | Byeju | 2022 | 1 | -1.08 | 31.376 | F | 1236-VO-TZ-OKUMU-VMF00261 | 1.3 | 1 | TZA | Kagera | TZ-05 | Missenyi | funestus | TZ-05_fune_2022 | TZ-05_fune_2022_01 | TZ-05_fune_2022_Q1 | TZ-05_Missenyi_fune_2022 | TZ-05_Missenyi_fune_2022_01 | TZ-05_Missenyi_fune_2022_Q1 |
| VBS85349 | TZ_KAG_41 | Joel Odero | Tanzania | Byeju | 2022 | 1 | -1.08 | 31.376 | F | 1236-VO-TZ-OKUMU-VMF00261 | 1.3 | 1 | TZA | Kagera | TZ-05 | Missenyi | funestus | TZ-05_fune_2022 | TZ-05_fune_2022_01 | TZ-05_fune_2022_Q1 | TZ-05_Missenyi_fune_2022 | TZ-05_Missenyi_fune_2022_01 | TZ-05_Missenyi_fune_2022_Q1 |
| VBS85350 | TZ_MTW_50 | Joel Odero | Tanzania | Mahurunga | 2022 | 10 | -10.538 | 40.279 | F | 1236-VO-TZ-OKUMU-VMF00261 | 1.3 | 4 | TZA | Mtwara | TZ-17 | Mtwara | funestus | TZ-17_fune_2022 | TZ-17_fune_2022_10 | TZ-17_fune_2022_Q4 | TZ-17_Mtwara_fune_2022 | TZ-17_Mtwara_fune_2022_10 | TZ-17_Mtwara_fune_2022_Q4 |
| VBS85351 | TZ_MTW_49 | Joel Odero | Tanzania | Mahurunga | 2022 | 10 | -10.538 | 40.279 | F | 1236-VO-TZ-OKUMU-VMF00261 | 1.3 | 4 | TZA | Mtwara | TZ-17 | Mtwara | funestus | TZ-17_fune_2022 | TZ-17_fune_2022_10 | TZ-17_fune_2022_Q4 | TZ-17_Mtwara_fune_2022 | TZ-17_Mtwara_fune_2022_10 | TZ-17_Mtwara_fune_2022_Q4 |
| VBS85352 | TZ_MTW_48 | Joel Odero | Tanzania | Mahurunga | 2022 | 10 | -10.538 | 40.279 | F | 1236-VO-TZ-OKUMU-VMF00261 | 1.3 | 4 | TZA | Mtwara | TZ-17 | Mtwara | funestus | TZ-17_fune_2022 | TZ-17_fune_2022_10 | TZ-17_fune_2022_Q4 | TZ-17_Mtwara_fune_2022 | TZ-17_Mtwara_fune_2022_10 | TZ-17_Mtwara_fune_2022_Q4 |
| VBS85353 | TZ_LND_25 | Joel Odero | Tanzania | Mtualonga | 2022 | 9 | -10.218 | 39.492 | F | 1236-VO-TZ-OKUMU-VMF00261 | 1.3 | 3 | TZA | Lindi | TZ-12 | Lindi | funestus | TZ-12_fune_2022 | TZ-12_fune_2022_09 | TZ-12_fune_2022_Q3 | TZ-12_Lindi_fune_2022 | TZ-12_Lindi_fune_2022_09 | TZ-12_Lindi_fune_2022_Q3 |
| VBS85354 | TZ_LND_24 | Joel Odero | Tanzania | Mtualonga | 2022 | 9 | -10.218 | 39.492 | F | 1236-VO-TZ-OKUMU-VMF00261 | 1.3 | 3 | TZA | Lindi | TZ-12 | Lindi | funestus | TZ-12_fune_2022 | TZ-12_fune_2022_09 | TZ-12_fune_2022_Q3 | TZ-12_Lindi_fune_2022 | TZ-12_Lindi_fune_2022_09 | TZ-12_Lindi_fune_2022_Q3 |

| | | | | | | | | | | | | | | | | | | | | | | | |
|---|---|---|---|---|---|---|---|---|---|---|---|---|---|---|---|---|---|---|---|---|---|---|---|
| VBS85355 | TZ_LND_23 | Joel Odero | Tanzania | Mtualonga | 2022 | 9 | -10.218 | 39.492 | F | 1236-VO-TZ-OKUMU-VMF00261 | 1.3 | 3 | TZA | Lindi | TZ-12 | Lindi | funestus | TZ-12_fune_2022 | TZ-12_fune_2022_09 | TZ-12_fune_2022_Q3 | TZ-12_Lindi_fune_2022 | TZ-12_Lindi_fune_2022_09 | TZ-12_Lindi_fune_2022_Q3 |
| VBS85356 | TZ_LND_22 | Joel Odero | Tanzania | Mtualonga | 2022 | 9 | -10.218 | 39.492 | F | 1236-VO-TZ-OKUMU-VMF00261 | 1.3 | 3 | TZA | Lindi | TZ-12 | Lindi | funestus | TZ-12_fune_2022 | TZ-12_fune_2022_09 | TZ-12_fune_2022_Q3 | TZ-12_Lindi_fune_2022 | TZ-12_Lindi_fune_2022_09 | TZ-12_Lindi_fune_2022_Q3 |
| VBS85357 | TZ_LND_21 | Joel Odero | Tanzania | Mtualonga | 2022 | 9 | -10.218 | 39.492 | F | 1236-VO-TZ-OKUMU-VMF00261 | 1.3 | 3 | TZA | Lindi | TZ-12 | Lindi | funestus | TZ-12_fune_2022 | TZ-12_fune_2022_09 | TZ-12_fune_2022_Q3 | TZ-12_Lindi_fune_2022 | TZ-12_Lindi_fune_2022_09 | TZ-12_Lindi_fune_2022_Q3 |
| VBS85358 | TZ_TNG_36 | Joel Odero | Tanzania | Kwamwale | 2022 | 5 | -5.157 | 38.729 | F | 1236-VO-TZ-OKUMU-VMF00261 | 1.3 | 2 | TZA | Tanga | TZ-25 | Muheza | funestus | TZ-25_fune_2022 | TZ-25_fune_2022_05 | TZ-25_fune_2022_Q2 | TZ-25_Muheza_fune_2022 | TZ-25_Muheza_fune_2022_05 | TZ-25_Muheza_fune_2022_Q2 |
| VBS85359 | TZ_TNG_35 | Joel Odero | Tanzania | Kwamwale | 2022 | 5 | -5.157 | 38.729 | F | 1236-VO-TZ-OKUMU-VMF00261 | 1.3 | 2 | TZA | Tanga | TZ-25 | Muheza | funestus | TZ-25_fune_2022 | TZ-25_fune_2022_05 | TZ-25_fune_2022_Q2 | TZ-25_Muheza_fune_2022 | TZ-25_Muheza_fune_2022_05 | TZ-25_Muheza_fune_2022_Q2 |
| VBS85360 | TZ_TNG_34 | Joel Odero | Tanzania | Kwamwale | 2022 | 5 | -5.157 | 38.729 | F | 1236-VO-TZ-OKUMU-VMF00261 | 1.3 | 2 | TZA | Tanga | TZ-25 | Muheza | funestus | TZ-25_fune_2022 | TZ-25_fune_2022_05 | TZ-25_fune_2022_Q2 | TZ-25_Muheza_fune_2022 | TZ-25_Muheza_fune_2022_05 | TZ-25_Muheza_fune_2022_Q2 |
| VBS85361 | TZ_TNG_33 | Joel Odero | Tanzania | Kwamwale | 2022 | 5 | -5.157 | 38.729 | F | 1236-VO-TZ-OKUMU-VMF00261 | 1.3 | 2 | TZA | Tanga | TZ-25 | Muheza | funestus | TZ-25_fune_2022 | TZ-25_fune_2022_05 | TZ-25_fune_2022_Q2 | TZ-25_Muheza_fune_2022 | TZ-25_Muheza_fune_2022_05 | TZ-25_Muheza_fune_2022_Q2 |
| VBS85362 | TZ_TNG_32 | Joel Odero | Tanzania | Kwamwale | 2022 | 5 | -5.157 | 38.729 | F | 1236-VO-TZ-OKUMU-VMF00261 | 1.3 | 2 | TZA | Tanga | TZ-25 | Muheza | funestus | TZ-25_fune_2022 | TZ-25_fune_2022_05 | TZ-25_fune_2022_Q2 | TZ-25_Muheza_fune_2022 | TZ-25_Muheza_fune_2022_05 | TZ-25_Muheza_fune_2022_Q2 |
| VBS85363 | TZ_TNG_31 | Joel Odero | Tanzania | Kwamwale | 2022 | 5 | -5.157 | 38.729 | F | 1236-VO-TZ-OKUMU-VMF00261 | 1.3 | 2 | TZA | Tanga | TZ-25 | Muheza | funestus | TZ-25_fune_2022 | TZ-25_fune_2022_05 | TZ-25_fune_2022_Q2 | TZ-25_Muheza_fune_2022 | TZ-25_Muheza_fune_2022_05 | TZ-25_Muheza_fune_2022_Q2 |
| VBS85364 | TZ_TNG_30 | Joel Odero | Tanzania | Kwamwale | 2022 | 5 | -5.157 | 38.729 | F | 1236-VO-TZ-OKUMU-VMF00261 | 1.3 | 2 | TZA | Tanga | TZ-25 | Muheza | funestus | TZ-25_fune_2022 | TZ-25_fune_2022_05 | TZ-25_fune_2022_Q2 | TZ-25_Muheza_fune_2022 | TZ-25_Muheza_fune_2022_05 | TZ-25_Muheza_fune_2022_Q2 |
| VBS85365 | TZ_TNG_29 | Joel Odero | Tanzania | Kwamwale | 2022 | 5 | -5.157 | 38.729 | F | 1236-VO-TZ-OKUMU-VMF00261 | 1.3 | 2 | TZA | Tanga | TZ-25 | Muheza | funestus | TZ-25_fune_2022 | TZ-25_fune_2022_05 | TZ-25_fune_2022_Q2 | TZ-25_Muheza_fune_2022 | TZ-25_Muheza_fune_2022_05 | TZ-25_Muheza_fune_2022_Q2 |
| VBS85366 | TZ_TNG_27 | Joel Odero | Tanzania | Kwamwale | 2022 | 5 | -5.157 | 38.729 | F | 1236-VO-TZ-OKUMU-VMF00261 | 1.3 | 2 | TZA | Tanga | TZ-25 | Muheza | funestus | TZ-25_fune_2022 | TZ-25_fune_2022_05 | TZ-25_fune_2022_Q2 | TZ-25_Muheza_fune_2022 | TZ-25_Muheza_fune_2022_05 | TZ-25_Muheza_fune_2022_Q2 |
| VBS85367 | TZ_TNG_26 | Joel Odero | Tanzania | Kwamwale | 2022 | 5 | -5.157 | 38.729 | F | 1236-VO-TZ-OKUMU-VMF00261 | 1.3 | 2 | TZA | Tanga | TZ-25 | Muheza | funestus | TZ-25_fune_2022 | TZ-25_fune_2022_05 | TZ-25_fune_2022_Q2 | TZ-25_Muheza_fune_2022 | TZ-25_Muheza_fune_2022_05 | TZ-25_Muheza_fune_2022_Q2 |
| VBS85368 | TZ_TNG_25 | Joel Odero | Tanzania | Kwamwale | 2022 | 5 | -5.157 | 38.729 | F | 1236-VO-TZ-OKUMU-VMF00261 | 1.3 | 2 | TZA | Tanga | TZ-25 | Muheza | funestus | TZ-25_fune_2022 | TZ-25_fune_2022_05 | TZ-25_fune_2022_Q2 | TZ-25_Muheza_fune_2022 | TZ-25_Muheza_fune_2022_05 | TZ-25_Muheza_fune_2022_Q2 |
| VBS85369 | TZ_TNG_24 | Joel Odero | Tanzania | Kwamwale | 2022 | 5 | -5.157 | 38.729 | F | 1236-VO-TZ-OKUMU-VMF00261 | 1.3 | 2 | TZA | Tanga | TZ-25 | Muheza | funestus | TZ-25_fune_2022 | TZ-25_fune_2022_05 | TZ-25_fune_2022_Q2 | TZ-25_Muheza_fune_2022 | TZ-25_Muheza_fune_2022_05 | TZ-25_Muheza_fune_2022_Q2 |
| VBS85370 | TZ_TNG_22 | Joel Odero | Tanzania | Kwamwale | 2022 | 5 | -5.157 | 38.729 | F | 1236-VO-TZ-OKUMU-VMF00261 | 1.3 | 2 | TZA | Tanga | TZ-25 | Muheza | funestus | TZ-25_fune_2022 | TZ-25_fune_2022_05 | TZ-25_fune_2022_Q2 | TZ-25_Muheza_fune_2022 | TZ-25_Muheza_fune_2022_05 | TZ-25_Muheza_fune_2022_Q2 |
| VBS85371 | TZ_TNG_21 | Joel Odero | Tanzania | Kwamwale | 2022 | 5 | -5.157 | 38.729 | F | 1236-VO-TZ-OKUMU-VMF00261 | 1.3 | 2 | TZA | Tanga | TZ-25 | Muheza | funestus | TZ-25_fune_2022 | TZ-25_fune_2022_05 | TZ-25_fune_2022_Q2 | TZ-25_Muheza_fune_2022 | TZ-25_Muheza_fune_2022_05 | TZ-25_Muheza_fune_2022_Q2 |
| VBS85372 | TZ_DOM_36 | Joel Odero | Tanzania | Mpwapwa | 2022 | 5 | -6.272 | 36.377 | F | 1236-VO-TZ-OKUMU-VMF00261 | 1.3 | 2 | TZA | Dodoma | TZ-03 | Mpwapwa | funestus | TZ-03_fune_2022 | TZ-03_fune_2022_05 | TZ-03_fune_2022_Q2 | TZ-03_Mpwapwa_fune_2022 | TZ-03_Mpwapwa_fune_2022_05 | TZ-03_Mpwapwa_fune_2022_Q2 |
| VBS85373 | TZ_DOM_35 | Joel Odero | Tanzania | Mpwapwa | 2022 | 5 | -6.272 | 36.377 | F | 1236-VO-TZ-OKUMU-VMF00261 | 1.3 | 2 | TZA | Dodoma | TZ-03 | Mpwapwa | funestus | TZ-03_fune_2022 | TZ-03_fune_2022_05 | TZ-03_fune_2022_Q2 | TZ-03_Mpwapwa_fune_2022 | TZ-03_Mpwapwa_fune_2022_05 | TZ-03_Mpwapwa_fune_2022_Q2 |
| VBS85374 | TZ_DOM_34 | Joel Odero | Tanzania | Mpwapwa | 2022 | 5 | -6.272 | 36.377 | F | 1236-VO-TZ-OKUMU-VMF00261 | 1.3 | 2 | TZA | Dodoma | TZ-03 | Mpwapwa | funestus | TZ-03_fune_2022 | TZ-03_fune_2022_05 | TZ-03_fune_2022_Q2 | TZ-03_Mpwapwa_fune_2022 | TZ-03_Mpwapwa_fune_2022_05 | TZ-03_Mpwapwa_fune_2022_Q2 |
| VBS85375 | TZ_DOM_33 | Joel Odero | Tanzania | Mpwapwa | 2022 | 5 | -6.272 | 36.377 | F | 1236-VO-TZ-OKUMU-VMF00261 | 1.3 | 2 | TZA | Dodoma | TZ-03 | Mpwapwa | funestus | TZ-03_fune_2022 | TZ-03_fune_2022_05 | TZ-03_fune_2022_Q2 | TZ-03_Mpwapwa_fune_2022 | TZ-03_Mpwapwa_fune_2022_05 | TZ-03_Mpwapwa_fune_2022_Q2 |
| VBS85376 | TZ_DOM_32 | Joel Odero | Tanzania | Mpwapwa | 2022 | 5 | -6.272 | 36.377 | F | 1236-VO-TZ-OKUMU-VMF00261 | 1.3 | 2 | TZA | Dodoma | TZ-03 | Mpwapwa | funestus | TZ-03_fune_2022 | TZ-03_fune_2022_05 | TZ-03_fune_2022_Q2 | TZ-03_Mpwapwa_fune_2022 | TZ-03_Mpwapwa_fune_2022_05 | TZ-03_Mpwapwa_fune_2022_Q2 |
| VBS85378 | TZ_DOM_29 | Joel Odero | Tanzania | Mpwapwa | 2022 | 5 | -6.272 | 36.377 | F | 1236-VO-TZ-OKUMU-VMF00261 | 1.3 | 2 | TZA | Dodoma | TZ-03 | Mpwapwa | funestus | TZ-03_fune_2022 | TZ-03_fune_2022_05 | TZ-03_fune_2022_Q2 | TZ-03_Mpwapwa_fune_2022 | TZ-03_Mpwapwa_fune_2022_05 | TZ-03_Mpwapwa_fune_2022_Q2 |
| VBS85379 | TZ_DOM_28 | Joel Odero | Tanzania | Mpwapwa | 2022 | 5 | -6.272 | 36.377 | F | 1236-VO-TZ-OKUMU-VMF00261 | 1.3 | 2 | TZA | Dodoma | TZ-03 | Mpwapwa | funestus | TZ-03_fune_2022 | TZ-03_fune_2022_05 | TZ-03_fune_2022_Q2 | TZ-03_Mpwapwa_fune_2022 | TZ-03_Mpwapwa_fune_2022_05 | TZ-03_Mpwapwa_fune_2022_Q2 |
| VBS85380 | TZ_DOM_27 | Joel Odero | Tanzania | Mpwapwa | 2022 | 5 | -6.272 | 36.377 | F | 1236-VO-TZ-OKUMU-VMF00261 | 1.3 | 2 | TZA | Dodoma | TZ-03 | Mpwapwa | funestus | TZ-03_fune_2022 | TZ-03_fune_2022_05 | TZ-03_fune_2022_Q2 | TZ-03_Mpwapwa_fune_2022 | TZ-03_Mpwapwa_fune_2022_05 | TZ-03_Mpwapwa_fune_2022_Q2 |
| VBS85381 | TZ_DOM_26 | Joel Odero | Tanzania | Mpwapwa | 2022 | 5 | -6.272 | 36.377 | F | 1236-VO-TZ-OKUMU-VMF00261 | 1.3 | 2 | TZA | Dodoma | TZ-03 | Mpwapwa | funestus | TZ-03_fune_2022 | TZ-03_fune_2022_05 | TZ-03_fune_2022_Q2 | TZ-03_Mpwapwa_fune_2022 | TZ-03_Mpwapwa_fune_2022_05 | TZ-03_Mpwapwa_fune_2022_Q2 |
| VBS85382 | TZ_DOM_25 | Joel Odero | Tanzania | Mpwapwa | 2022 | 5 | -6.272 | 36.377 | F | 1236-VO-TZ-OKUMU-VMF00261 | 1.3 | 2 | TZA | Dodoma | TZ-03 | Mpwapwa | funestus | TZ-03_fune_2022 | TZ-03_fune_2022_05 | TZ-03_fune_2022_Q2 | TZ-03_Mpwapwa_fune_2022 | TZ-03_Mpwapwa_fune_2022_05 | TZ-03_Mpwapwa_fune_2022_Q2 |
| VBS85384 | TZ_DOM_23 | Joel Odero | Tanzania | Mpwapwa | 2022 | 5 | -6.272 | 36.377 | F | 1236-VO-TZ-OKUMU-VMF00261 | 1.3 | 2 | TZA | Dodoma | TZ-03 | Mpwapwa | funestus | TZ-03_fune_2022 | TZ-03_fune_2022_05 | TZ-03_fune_2022_Q2 | TZ-03_Mpwapwa_fune_2022 | TZ-03_Mpwapwa_fune_2022_05 | TZ-03_Mpwapwa_fune_2022_Q2 |
| VBS85385 | TZ_DOM_22 | Joel Odero | Tanzania | Mpwapwa | 2022 | 5 | -6.272 | 36.377 | F | 1236-VO-TZ-OKUMU-VMF00261 | 1.3 | 2 | TZA | Dodoma | TZ-03 | Mpwapwa | funestus | TZ-03_fune_2022 | TZ-03_fune_2022_05 | TZ-03_fune_2022_Q2 | TZ-03_Mpwapwa_fune_2022 | TZ-03_Mpwapwa_fune_2022_05 | TZ-03_Mpwapwa_fune_2022_Q2 |
| VBS85386 | TZ_DOM_21 | Joel Odero | Tanzania | Mpwapwa | 2022 | 5 | -6.272 | 36.377 | F | 1236-VO-TZ-OKUMU-VMF00261 | 1.3 | 2 | TZA | Dodoma | TZ-03 | Mpwapwa | funestus | TZ-03_fune_2022 | TZ-03_fune_2022_05 | TZ-03_fune_2022_Q2 | TZ-03_Mpwapwa_fune_2022 | TZ-03_Mpwapwa_fune_2022_05 | TZ-03_Mpwapwa_fune_2022_Q2 |
| VBS85387 | TZ_RUV_44 | Joel Odero | Tanzania | Nandembo | 2022 | 3 | -10.967 | 37.23 | F | 1236-VO-TZ-OKUMU-VMF00261 | 1.3 | 1 | TZA | Ruvuma | TZ-21 | Tunduru | funestus | TZ-21_fune_2022 | TZ-21_fune_2022_03 | TZ-21_fune_2022_Q1 | TZ-21_Tunduru_fune_2022 | TZ-21_Tunduru_fune_2022_03 | TZ-21_Tunduru_fune_2022_Q1 |
| VBS85388 | TZ_RUV_43 | Joel Odero | Tanzania | Nandembo | 2022 | 3 | -10.967 | 37.23 | F | 1236-VO-TZ-OKUMU-VMF00261 | 1.3 | 1 | TZA | Ruvuma | TZ-21 | Tunduru | funestus | TZ-21_fune_2022 | TZ-21_fune_2022_03 | TZ-21_fune_2022_Q1 | TZ-21_Tunduru_fune_2022 | TZ-21_Tunduru_fune_2022_03 | TZ-21_Tunduru_fune_2022_Q1 |
| VBS85389 | TZ_RUV_42 | Joel Odero | Tanzania | Nandembo | 2022 | 3 | -10.967 | 37.23 | F | 1236-VO-TZ-OKUMU-VMF00261 | 1.3 | 1 | TZA | Ruvuma | TZ-21 | Tunduru | funestus | TZ-21_fune_2022 | TZ-21_fune_2022_03 | TZ-21_fune_2022_Q1 | TZ-21_Tunduru_fune_2022 | TZ-21_Tunduru_fune_2022_03 | TZ-21_Tunduru_fune_2022_Q1 |
| VBS85390 | TZ_RUV_41 | Joel Odero | Tanzania | Nandembo | 2022 | 3 | -10.967 | 37.23 | F | 1236-VO-TZ-OKUMU-VMF00261 | 1.3 | 1 | TZA | Ruvuma | TZ-21 | Tunduru | funestus | TZ-21_fune_2022 | TZ-21_fune_2022_03 | TZ-21_fune_2022_Q1 | TZ-21_Tunduru_fune_2022 | TZ-21_Tunduru_fune_2022_03 | TZ-21_Tunduru_fune_2022_Q1 |
| VBS85391 | TZ_RUV_40 | Joel Odero | Tanzania | Nandembo | 2022 | 3 | -10.967 | 37.23 | F | 1236-VO-TZ-OKUMU-VMF00261 | 1.3 | 1 | TZA | Ruvuma | TZ-21 | Tunduru | funestus | TZ-21_fune_2022 | TZ-21_fune_2022_03 | TZ-21_fune_2022_Q1 | TZ-21_Tunduru_fune_2022 | TZ-21_Tunduru_fune_2022_03 | TZ-21_Tunduru_fune_2022_Q1 |
| VBS85392 | TZ_RUV_39 | Joel Odero | Tanzania | Nandembo | 2022 | 3 | -10.967 | 37.23 | F | 1236-VO-TZ-OKUMU-VMF00261 | 1.3 | 1 | TZA | Ruvuma | TZ-21 | Tunduru | funestus | TZ-21_fune_2022 | TZ-21_fune_2022_03 | TZ-21_fune_2022_Q1 | TZ-21_Tunduru_fune_2022 | TZ-21_Tunduru_fune_2022_03 | TZ-21_Tunduru_fune_2022_Q1 |
| VBS85393 | TZ_RUV_38 | Joel Odero | Tanzania | Nandembo | 2022 | 3 | -10.967 | 37.23 | F | 1236-VO-TZ-OKUMU-VMF00261 | 1.3 | 1 | TZA | Ruvuma | TZ-21 | Tunduru | funestus | TZ-21_fune_2022 | TZ-21_fune_2022_03 | TZ-21_fune_2022_Q1 | TZ-21_Tunduru_fune_2022 | TZ-21_Tunduru_fune_2022_03 | TZ-21_Tunduru_fune_2022_Q1 |
| VBS85394 | TZ_RUV_37 | Joel Odero | Tanzania | Nandembo | 2022 | 3 | -10.967 | 37.23 | F | 1236-VO-TZ-OKUMU-VMF00261 | 1.3 | 1 | TZA | Ruvuma | TZ-21 | Tunduru | funestus | TZ-21_fune_2022 | TZ-21_fune_2022_03 | TZ-21_fune_2022_Q1 | TZ-21_Tunduru_fune_2022 | TZ-21_Tunduru_fune_2022_03 | TZ-21_Tunduru_fune_2022_Q1 |
| VBS85395 | TZ_RUV_36 | Joel Odero | Tanzania | Nandembo | 2022 | 3 | -10.967 | 37.23 | F | 1236-VO-TZ-OKUMU-VMF00261 | 1.3 | 1 | TZA | Ruvuma | TZ-21 | Tunduru | funestus | TZ-21_fune_2022 | TZ-21_fune_2022_03 | TZ-21_fune_2022_Q1 | TZ-21_Tunduru_fune_2022 | TZ-21_Tunduru_fune_2022_03 | TZ-21_Tunduru_fune_2022_Q1 |
| VBS85396 | TZ_RUV_35 | Joel Odero | Tanzania | Nandembo | 2022 | 3 | -10.967 | 37.23 | F | 1236-VO-TZ-OKUMU-VMF00261 | 1.3 | 1 | TZA | Ruvuma | TZ-21 | Tunduru | funestus | TZ-21_fune_2022 | TZ-21_fune_2022_03 | TZ-21_fune_2022_Q1 | TZ-21_Tunduru_fune_2022 | TZ-21_Tunduru_fune_2022_03 | TZ-21_Tunduru_fune_2022_Q1 |
| VBS85397 | TZ_RUV_34 | Joel Odero | Tanzania | Nandembo | 2022 | 3 | -10.967 | 37.23 | F | 1236-VO-TZ-OKUMU-VMF00261 | 1.3 | 1 | TZA | Ruvuma | TZ-21 | Tunduru | funestus | TZ-21_fune_2022 | TZ-21_fune_2022_03 | TZ-21_fune_2022_Q1 | TZ-21_Tunduru_fune_2022 | TZ-21_Tunduru_fune_2022_03 | TZ-21_Tunduru_fune_2022_Q1 |
| VBS85398 | TZ_RUV_33 | Joel Odero | Tanzania | Nandembo | 2022 | 3 | -10.967 | 37.23 | F | 1236-VO-TZ-OKUMU-VMF00261 | 1.3 | 1 | TZA | Ruvuma | TZ-21 | Tunduru | funestus | TZ-21_fune_2022 | TZ-21_fune_2022_03 | TZ-21_fune_2022_Q1 | TZ-21_Tunduru_fune_2022 | TZ-21_Tunduru_fune_2022_03 | TZ-21_Tunduru_fune_2022_Q1 |
| VBS85399 | TZ_RUV_32 | Joel Odero | Tanzania | Nandembo | 2022 | 3 | -10.967 | 37.23 | F | 1236-VO-TZ-OKUMU-VMF00261 | 1.3 | 1 | TZA | Ruvuma | TZ-21 | Tunduru | funestus | TZ-21_fune_2022 | TZ-21_fune_2022_03 | TZ-21_fune_2022_Q1 | TZ-21_Tunduru_fune_2022 | TZ-21_Tunduru_fune_2022_03 | TZ-21_Tunduru_fune_2022_Q1 |
| VBS85400 | TZ_RUV_31 | Joel Odero | Tanzania | Nandembo | 2022 | 3 | -10.967 | 37.23 | F | 1236-VO-TZ-OKUMU-VMF00261 | 1.3 | 1 | TZA | Ruvuma | TZ-21 | Tunduru | funestus | TZ-21_fune_2022 | TZ-21_fune_2022_03 | TZ-21_fune_2022_Q1 | TZ-21_Tunduru_fune_2022 | TZ-21_Tunduru_fune_2022_03 | TZ-21_Tunduru_fune_2022_Q1 |
| VBS85401 | TZ_RUV_30 | Joel Odero | Tanzania | Nandembo | 2022 | 3 | -10.967 | 37.23 | F | 1236-VO-TZ-OKUMU-VMF00261 | 1.3 | 1 | TZA | Ruvuma | TZ-21 | Tunduru | funestus | TZ-21_fune_2022 | TZ-21_fune_2022_03 | TZ-21_fune_2022_Q1 | TZ-21_Tunduru_fune_2022 | TZ-21_Tunduru_fune_2022_03 | TZ-21_Tunduru_fune_2022_Q1 |
| VBS85402 | TZ_RUV_29 | Joel Odero | Tanzania | Nandembo | 2022 | 3 | -10.967 | 37.23 | F | 1236-VO-TZ-OKUMU-VMF00261 | 1.3 | 1 | TZA | Ruvuma | TZ-21 | Tunduru | funestus | TZ-21_fune_2022 | TZ-21_fune_2022_03 | TZ-21_fune_2022_Q1 | TZ-21_Tunduru_fune_2022 | TZ-21_Tunduru_fune_2022_03 | TZ-21_Tunduru_fune_2022_Q1 |
| VBS85403 | TZ_RUV_28 | Joel Odero | Tanzania | Nandembo | 2022 | 3 | -10.967 | 37.23 | F | 1236-VO-TZ-OKUMU-VMF00261 | 1.3 | 1 | TZA | Ruvuma | TZ-21 | Tunduru | funestus | TZ-21_fune_2022 | TZ-21_fune_2022_03 | TZ-21_fune_2022_Q1 | TZ-21_Tunduru_fune_2022 | TZ-21_Tunduru_fune_2022_03 | TZ-21_Tunduru_fune_2022_Q1 |
| VBS85404 | TZ_RUV_24 | Joel Odero | Tanzania | Nandembo | 2022 | 3 | -10.967 | 37.23 | F | 1236-VO-TZ-OKUMU-VMF00261 | 1.3 | 1 | TZA | Ruvuma | TZ-21 | Tunduru | funestus | TZ-21_fune_2022 | TZ-21_fune_2022_03 | TZ-21_fune_2022_Q1 | TZ-21_Tunduru_fune_2022 | TZ-21_Tunduru_fune_2022_03 | TZ-21_Tunduru_fune_2022_Q1 |
| VBS85405 | TZ_RUV_23 | Joel Odero | Tanzania | Nandembo | 2022 | 3 | -10.967 | 37.23 | F | 1236-VO-TZ-OKUMU-VMF00261 | 1.3 | 1 | TZA | Ruvuma | TZ-21 | Tunduru | funestus | TZ-21_fune_2022 | TZ-21_fune_2022_03 | TZ-21_fune_2022_Q1 | TZ-21_Tunduru_fune_2022 | TZ-21_Tunduru_fune_2022_03 | TZ-21_Tunduru_fune_2022_Q1 |
| VBS85406 | TZ_RUV_22 | Joel Odero | Tanzania | Nandembo | 2022 | 3 | -10.967 | 37.23 | F | 1236-VO-TZ-OKUMU-VMF00261 | 1.3 | 1 | TZA | Ruvuma | TZ-21 | Tunduru | funestus | TZ-21_fune_2022 | TZ-21_fune_2022_03 | TZ-21_fune_2022_Q1 | TZ-21_Tunduru_fune_2022 | TZ-21_Tunduru_fune_2022_03 | TZ-21_Tunduru_fune_2022_Q1 |
| VBS85407 | TZ_RUV_21 | Joel Odero | Tanzania | Nandembo | 2022 | 3 | -10.967 | 37.23 | F | 1236-VO-TZ-OKUMU-VMF00261 | 1.3 | 1 | TZA | Ruvuma | TZ-21 | Tunduru | funestus | TZ-21_fune_2022 | TZ-21_fune_2022_03 | TZ-21_fune_2022_Q1 | TZ-21_Tunduru_fune_2022 | TZ-21_Tunduru_fune_2022_03 | TZ-21_Tunduru_fune_2022_Q1 |
| VBS85408 | TZ_KAG_64 | Joel Odero | Tanzania | Byeju | 2022 | 1 | -1.08 | 31.376 | F | 1236-VO-TZ-OKUMU-VMF00261 | 1.3 | 1 | TZA | Kagera | TZ-05 | Missenyi | funestus | TZ-05_fune_2022 | TZ-05_fune_2022_01 | TZ-05_fune_2022_Q1 | TZ-05_Missenyi_fune_2022 | TZ-05_Missenyi_fune_2022_01 | TZ-05_Missenyi_fune_2022_Q1 |
| VBS85409 | TZ_KAG_63 | Joel Odero | Tanzania | Byeju | 2022 | 1 | -1.08 | 31.376 | F | 1236-VO-TZ-OKUMU-VMF00261 | 1.3 | 1 | TZA | Kagera | TZ-05 | Missenyi | funestus | TZ-05_fune_2022 | TZ-05_fune_2022_01 | TZ-05_fune_2022_Q1 | TZ-05_Missenyi_fune_2022 | TZ-05_Missenyi_fune_2022_01 | TZ-05_Missenyi_fune_2022_Q1 |
| VBS85410 | TZ_KAG_62 | Joel Odero | Tanzania | Byeju | 2022 | 1 | -1.08 | 31.376 | F | 1236-VO-TZ-OKUMU-VMF00261 | 1.3 | 1 | TZA | Kagera | TZ-05 | Missenyi | funestus | TZ-05_fune_2022 | TZ-05_fune_2022_01 | TZ-05_fune_2022_Q1 | TZ-05_Missenyi_fune_2022 | TZ-05_Missenyi_fune_2022_01 | TZ-05_Missenyi_fune_2022_Q1 |
| VBS85411 | TZ_KAG_61 | Joel Odero | Tanzania | Byeju | 2022 | 1 | -1.08 | 31.376 | F | 1236-VO-TZ-OKUMU-VMF00261 | 1.3 | 1 | TZA | Kagera | TZ-05 | Missenyi | funestus | TZ-05_fune_2022 | TZ-05_fune_2022_01 | TZ-05_fune_2022_Q1 | TZ-05_Missenyi_fune_2022 | TZ-05_Missenyi_fune_2022_01 | TZ-05_Missenyi_fune_2022_Q1 |
| VBS85412 | TZ_KAG_60 | Joel Odero | Tanzania | Byeju | 2022 | 1 | -1.08 | 31.376 | F | 1236-VO-TZ-OKUMU-VMF00261 | 1.3 | 1 | TZA | Kagera | TZ-05 | Missenyi | funestus | TZ-05_fune_2022 | TZ-05_fune_2022_01 | TZ-05_fune_2022_Q1 | TZ-05_Missenyi_fune_2022 | TZ-05_Missenyi_fune_2022_01 | TZ-05_Missenyi_fune_2022_Q1 |
| VBS85413 | TZ_KAG_59 | Joel Odero | Tanzania | Byeju | 2022 | 1 | -1.08 | 31.376 | F | 1236-VO-TZ-OKUMU-VMF00261 | 1.3 | 1 | TZA | Kagera | TZ-05 | Missenyi | funestus | TZ-05_fune_2022 | TZ-05_fune_2022_01 | TZ-05_fune_2022_Q1 | TZ-05_Missenyi_fune_2022 | TZ-05_Missenyi_fune_2022_01 | TZ-05_Missenyi_fune_2022_Q1 |
| VBS85414 | TZ_KAG_58 | Joel Odero | Tanzania | Byeju | 2022 | 1 | -1.08 | 31.376 | F | 1236-VO-TZ-OKUMU-VMF00261 | 1.3 | 1 | TZA | Kagera | TZ-05 | Missenyi | funestus | TZ-05_fune_2022 | TZ-05_fune_2022_01 | TZ-05_fune_2022_Q1 | TZ-05_Missenyi_fune_2022 | TZ-05_Missenyi_fune_2022_01 | TZ-05_Missenyi_fune_2022_Q1 |
| VBS85415 | TZ_KAG_57 | Joel Odero | Tanzania | Byeju | 2022 | 1 | -1.08 | 31.376 | F | 1236-VO-TZ-OKUMU-VMF00261 | 1.3 | 1 | TZA | Kagera | TZ-05 | Missenyi | funestus | TZ-05_fune_2022 | TZ-05_fune_2022_01 | TZ-05_fune_2022_Q1 | TZ-05_Missenyi_fune_2022 | TZ-05_Missenyi_fune_2022_01 | TZ-05_Missenyi_fune_2022_Q1 |
| VBS85416 | TZ_KAG_56 | Joel Odero | Tanzania | Byeju | 2022 | 1 | -1.08 | 31.376 | F | 1236-VO-TZ-OKUMU-VMF00261 | 1.3 | 1 | TZA | Kagera | TZ-05 | Missenyi | funestus | TZ-05_fune_2022 | TZ-05_fune_2022_01 | TZ-05_fune_2022_Q1 | TZ-05_Missenyi_fune_2022 | TZ-05_Missenyi_fune_2022_01 | TZ-05_Missenyi_fune_2022_Q1 |
| VBS85417 | TZ_KAG_55 | Joel Odero | Tanzania | Byeju | 2022 | 1 | -1.08 | 31.376 | F | 1236-VO-TZ-OKUMU-VMF00261 | 1.3 | 1 | TZA | Kagera | TZ-05 | Missenyi | funestus | TZ-05_fune_2022 | TZ-05_fune_2022_01 | TZ-05_fune_2022_Q1 | TZ-05_Missenyi_fune_2022 | TZ-05_Missenyi_fune_2022_01 | TZ-05_Missenyi_fune_2022_Q1 |
| VBS85419 | TZ_RUV_27 | Joel Odero | Tanzania | Nandembo | 2022 | 3 | -10.967 | 37.23 | F | 1236-VO-TZ-OKUMU-VMF00261 | 1.3 | 1 | TZA | Ruvuma | TZ-21 | Tunduru | funestus | TZ-21_fune_2022 | TZ-21_fune_2022_03 | TZ-21_fune_2022_Q1 | TZ-21_Tunduru_fune_2022 | TZ-21_Tunduru_fune_2022_03 | TZ-21_Tunduru_fune_2022_Q1 |
| VBS85420 | TZ_RUV_26 | Joel Odero | Tanzania | Nandembo | 2022 | 3 | -10.967 | 37.23 | F | 1236-VO-TZ-OKUMU-VMF00261 | 1.3 | 1 | TZA | Ruvuma | TZ-21 | Tunduru | funestus | TZ-21_fune_2022 | TZ-21_fune_2022_03 | TZ-21_fune_2022_Q1 | TZ-21_Tunduru_fune_2022 | TZ-21_Tunduru_fune_2022_03 | TZ-21_Tunduru_fune_2022_Q1 |
| VBS85421 | TZ_RUV_25 | Joel Odero | Tanzania | Nandembo | 2022 | 3 | -10.967 | 37.23 | F | 1236-VO-TZ-OKUMU-VMF00261 | 1.3 | 1 | TZA | Ruvuma | TZ-21 | Tunduru | funestus | TZ-21_fune_2022 | TZ-21_fune_2022_03 | TZ-21_fune_2022_Q1 | TZ-21_Tunduru_fune_2022 | TZ-21_Tunduru_fune_2022_03 | TZ-21_Tunduru_fune_2022_Q1 |
| VBS85422 | TZ_KAG_53 | Joel Odero | Tanzania | Byeju | 2022 | 1 | -1.08 | 31.376 | F | 1236-VO-TZ-OKUMU-VMF00261 | 1.3 | 1 | TZA | Kagera | TZ-05 | Missenyi | funestus | TZ-05_fune_2022 | TZ-05_fune_2022_01 | TZ-05_fune_2022_Q1 | TZ-05_Missenyi_fune_2022 | TZ-05_Missenyi_fune_2022_01 | TZ-05_Missenyi_fune_2022_Q1 |
| VBS85423 | TZ_KAG_52 | Joel Odero | Tanzania | Byeju | 2022 | 1 | -1.08 | 31.376 | F | 1236-VO-TZ-OKUMU-VMF00261 | 1.3 | 1 | TZA | Kagera | TZ-05 | Missenyi | funestus | TZ-05_fune_2022 | TZ-05_fune_2022_01 | TZ-05_fune_2022_Q1 | TZ-05_Missenyi_fune_2022 | TZ-05_Missenyi_fune_2022_01 | TZ-05_Missenyi_fune_2022_Q1 |
| VBS85425 | TZ_MTW_42 | Joel Odero | Tanzania | Mahurunga | 2022 | 10 | -10.538 | 40.279 | F | 1236-VO-TZ-OKUMU-VMF00261 | 1.3 | 4 | TZA | Mtwara | TZ-17 | Mtwara | funestus | TZ-17_fune_2022 | TZ-17_fune_2022_10 | TZ-17_fune_2022_Q4 | TZ-17_Mtwara_fune_2022 | TZ-17_Mtwara_fune_2022_10 | TZ-17_Mtwara_fune_2022_Q4 |
| VBS85426 | TZ_MTW_37 | Joel Odero | Tanzania | Mahurunga | 2022 | 10 | -10.538 | 40.279 | F | 1236-VO-TZ-OKUMU-VMF00261 | 1.3 | 4 | TZA | Mtwara | TZ-17 | Mtwara | funestus | TZ-17_fune_2022 | TZ-17_fune_2022_10 | TZ-17_fune_2022_Q4 | TZ-17_Mtwara_fune_2022 | TZ-17_Mtwara_fune_2022_10 | TZ-17_Mtwara_fune_2022_Q4 |
| VBS85427 | TZ_MTW_36 | Joel Odero | Tanzania | Mahurunga | 2022 | 10 | -10.538 | 40.279 | F | 1236-VO-TZ-OKUMU-VMF00261 | 1.3 | 4 | TZA | Mtwara | TZ-17 | Mtwara | funestus | TZ-17_fune_2022 | TZ-17_fune_2022_10 | TZ-17_fune_2022_Q4 | TZ-17_Mtwara_fune_2022 | TZ-17_Mtwara_fune_2022_10 | TZ-17_Mtwara_fune_2022_Q4 |
| VBS85428 | TZ_MTW_30 | Joel Odero | Tanzania | Mahurunga | 2022 | 10 | -10.538 | 40.279 | F | 1236-VO-TZ-OKUMU-VMF00261 | 1.3 | 4 | TZA | Mtwara | TZ-17 | Mtwara | funestus | TZ-17_fune_2022 | TZ-17_fune_2022_10 | TZ-17_fune_2022_Q4 | TZ-17_Mtwara_fune_2022 | TZ-17_Mtwara_fune_2022_10 | TZ-17_Mtwara_fune_2022_Q4 |
| VBS85429 | TZ_MTW_29 | Joel Odero | Tanzania | Mahurunga | 2022 | 10 | -10.538 | 40.279 | F | 1236-VO-TZ-OKUMU-VMF00261 | 1.3 | 4 | TZA | Mtwara | TZ-17 | Mtwara | funestus | TZ-17_fune_2022 | TZ-17_fune_2022_10 | TZ-17_fune_2022_Q4 | TZ-17_Mtwara_fune_2022 | TZ-17_Mtwara_fune_2022_10 | TZ-17_Mtwara_fune_2022_Q4 |
| VBS85430 | TZ_MTW_28 | Joel Odero | Tanzania | Mahurunga | 2022 | 10 | -10.538 | 40.279 | F | 1236-VO-TZ-OKUMU-VMF00261 | 1.3 | 4 | TZA | Mtwara | TZ-17 | Mtwara | funestus | TZ-17_fune_2022 | TZ-17_fune_2022_10 | TZ-17_fune_2022_Q4 | TZ-17_Mtwara_fune_2022 | TZ-17_Mtwara_fune_2022_10 | TZ-17_Mtwara_fune_2022_Q4 |
| VBS85431 | TZ_MTW_26 | Joel Odero | Tanzania | Mahurunga | 2022 | 10 | -10.538 | 40.279 | F | 1236-VO-TZ-OKUMU-VMF00261 | 1.3 | 4 | TZA | Mtwara | TZ-17 | Mtwara | funestus | TZ-17_fune_2022 | TZ-17_fune_2022_10 | TZ-17_fune_2022_Q4 | TZ-17_Mtwara_fune_2022 | TZ-17_Mtwara_fune_2022_10 | TZ-17_Mtwara_fune_2022_Q4 |
| VBS85432 | TZ_MTW_25 | Joel Odero | Tanzania | Mahurunga | 2022 | 10 | -10.538 | 40.279 | F | 1236-VO-TZ-OKUMU-VMF00261 | 1.3 | 4 | TZA | Mtwara | TZ-17 | Mtwara | funestus | TZ-17_fune_2022 | TZ-17_fune_2022_10 | TZ-17_fune_2022_Q4 | TZ-17_Mtwara_fune_2022 | TZ-17_Mtwara_fune_2022_10 | TZ-17_Mtwara_fune_2022_Q4 |
| VBS85433 | TZ_MOR_54 | Joel Odero | Tanzania | Ikwambi | 2023 | 3 | -7.984 | 36.818 | F | 1236-VO-TZ-OKUMU-VMF00261 | 1.3 | 1 | TZA | Morogoro | TZ-16 | Kilombero | funestus | TZ-16_fune_2023 | TZ-16_fune_2023_03 | TZ-16_fune_2023_Q1 | TZ-16_Kilombero_fune_2023 | TZ-16_Kilombero_fune_2023_03 | TZ-16_Kilombero_fune_2023_Q1 |
| VBS85434 | TZ_MOR_53 | Joel Odero | Tanzania | Ikwambi | 2023 | 3 | -7.984 | 36.818 | F | 1236-VO-TZ-OKUMU-VMF00261 | 1.3 | 1 | TZA | Morogoro | TZ-16 | Kilombero | funestus | TZ-16_fune_2023 | TZ-16_fune_2023_03 | TZ-16_fune_2023_Q1 | TZ-16_Kilombero_fune_2023 | TZ-16_Kilombero_fune_2023_03 | TZ-16_Kilombero_fune_2023_Q1 |
| VBS85435 | TZ_MOR_52 | Joel Odero | Tanzania | Ikwambi | 2023 | 3 | -7.984 | 36.818 | F | 1236-VO-TZ-OKUMU-VMF00261 | 1.3 | 1 | TZA | Morogoro | TZ-16 | Kilombero | funestus | TZ-16_fune_2023 | TZ-16_fune_2023_03 | TZ-16_fune_2023_Q1 | TZ-16_Kilombero_fune_2023 | TZ-16_Kilombero_fune_2023_03 | TZ-16_Kilombero_fune_2023_Q1 |
| VBS85436 | TZ_MOR_51 | Joel Odero | Tanzania | Ikwambi | 2023 | 3 | -7.984 | 36.818 | F | 1236-VO-TZ-OKUMU-VMF00261 | 1.3 | 1 | TZA | Morogoro | TZ-16 | Kilombero | funestus | TZ-16_fune_2023 | TZ-16_fune_2023_03 | TZ-16_fune_2023_Q1 | TZ-16_Kilombero_fune_2023 | TZ-16_Kilombero_fune_2023_03 | TZ-16_Kilombero_fune_2023_Q1 |
| VBS85437 | TZ_MOR_50 | Joel Odero | Tanzania | Ikwambi | 2023 | 3 | -7.984 | 36.818 | F | 1236-VO-TZ-OKUMU-VMF00261 | 1.3 | 1 | TZA | Morogoro | TZ-16 | Kilombero | funestus | TZ-16_fune_2023 | TZ-16_fune_2023_03 | TZ-16_fune_2023_Q1 | TZ-16_Kilombero_fune_2023 | TZ-16_Kilombero_fune_2023_03 | TZ-16_Kilombero_fune_2023_Q1 |
| VBS85438 | TZ_MOR_49 | Joel Odero | Tanzania | Ikwambi | 2023 | 3 | -7.984 | 36.818 | F | 1236-VO-TZ-OKUMU-VMF00261 | 1.3 | 1 | TZA | Morogoro | TZ-16 | Kilombero | funestus | TZ-16_fune_2023 | TZ-16_fune_2023_03 | TZ-16_fune_2023_Q1 | TZ-16_Kilombero_fune_2023 | TZ-16_Kilombero_fune_2023_03 | TZ-16_Kilombero_fune_2023_Q1 |
| VBS85439 | TZ_MOR_48 | Joel Odero | Tanzania | Ikwambi | 2023 | 3 | -7.984 | 36.818 | F | 1236-VO-TZ-OKUMU-VMF00261 | 1.3 | 1 | TZA | Morogoro | TZ-16 | Kilombero | funestus | TZ-16_fune_2023 | TZ-16_fune_2023_03 | TZ-16_fune_2023_Q1 | TZ-16_Kilombero_fune_2023 | TZ-16_Kilombero_fune_2023_03 | TZ-16_Kilombero_fune_2023_Q1 |
| VBS85440 | TZ_MOR_47 | Joel Odero | Tanzania | Ikwambi | 2023 | 3 | -7.984 | 36.818 | F | 1236-VO-TZ-OKUMU-VMF00261 | 1.3 | 1 | TZA | Morogoro | TZ-16 | Kilombero | funestus | TZ-16_fune_2023 | TZ-16_fune_2023_03 | TZ-16_fune_2023_Q1 | TZ-16_Kilombero_fune_2023 | TZ-16_Kilombero_fune_2023_03 | TZ-16_Kilombero_fune_2023_Q1 |
| VBS85441 | TZ_MOR_45 | Joel Odero | Tanzania | Ikwambi | 2023 | 3 | -7.984 | 36.818 | F | 1236-VO-TZ-OKUMU-VMF00261 | 1.3 | 1 | TZA | Morogoro | TZ-16 | Kilombero | funestus | TZ-16_fune_2023 | TZ-16_fune_2023_03 | TZ-16_fune_2023_Q1 | TZ-16_Kilombero_fune_2023 | TZ-16_Kilombero_fune_2023_03 | TZ-16_Kilombero_fune_2023_Q1 |
| VBS85442 | TZ_MOR_44 | Joel Odero | Tanzania | Ikwambi | 2023 | 3 | -7.984 | 36.818 | F | 1236-VO-TZ-OKUMU-VMF00261 | 1.3 | 1 | TZA | Morogoro | TZ-16 | Kilombero | funestus | TZ-16_fune_2023 | TZ-16_fune_2023_03 | TZ-16_fune_2023_Q1 | TZ-16_Kilombero_fune_2023 | TZ-16_Kilombero_fune_2023_03 | TZ-16_Kilombero_fune_2023_Q1 |
| VBS85443 | TZ_MOR_43 | Joel Odero | Tanzania | Ikwambi | 2023 | 3 | -7.984 | 36.818 | F | 1236-VO-TZ-OKUMU-VMF00261 | 1.3 | 1 | TZA | Morogoro | TZ-16 | Kilombero | funestus | TZ-16_fune_2023 | TZ-16_fune_2023_03 | TZ-16_fune_2023_Q1 | TZ-16_Kilombero_fune_2023 | TZ-16_Kilombero_fune_2023_03 | TZ-16_Kilombero_fune_2023_Q1 |
| VBS85444 | TZ_MOR_42 | Joel Odero | Tanzania | Ikwambi | 2023 | 3 | -7.984 | 36.818 | F | 1236-VO-TZ-OKUMU-VMF00261 | 1.3 | 1 | TZA | Morogoro | TZ-16 | Kilombero | funestus | TZ-16_fune_2023 | TZ-16_fune_2023_03 | TZ-16_fune_2023_Q1 | TZ-16_Kilombero_fune_2023 | TZ-16_Kilombero_fune_2023_03 | TZ-16_Kilombero_fune_2023_Q1 |

| | | | | | | | | | | | | | | | | | | | | | | | |
|---|---|---|---|---|---|---|---|---|---|---|---|---|---|---|---|---|---|---|---|---|---|---|---|
| VBS85445 | TZ_MOR_41 | Joel Odero | Tanzania | Ikwambi | 2023 | 3 | -7.984 | 36.818 | F | 1236-VO-TZ-OKUMU-VMF00261 | 1.3 | 1 | TZA | Morogoro | TZ-16 | Kilombero | funestus | TZ-16_fune_2023 | TZ-16_fune_2023_03 | TZ-16_fune_2023_Q1 | TZ-16_Kilombero_fune_2023 | TZ-16_Kilombero_fune_2023_03 | TZ-16_Kilombero_fune_2023_Q1 |
| VBS85446 | TZ_MOR_40 | Joel Odero | Tanzania | Ikwambi | 2023 | 3 | -7.984 | 36.818 | F | 1236-VO-TZ-OKUMU-VMF00261 | 1.3 | 1 | TZA | Morogoro | TZ-16 | Kilombero | funestus | TZ-16_fune_2023 | TZ-16_fune_2023_03 | TZ-16_fune_2023_Q1 | TZ-16_Kilombero_fune_2023 | TZ-16_Kilombero_fune_2023_03 | TZ-16_Kilombero_fune_2023_Q1 |
| VBS85447 | TZ_MOR_39 | Joel Odero | Tanzania | Ikwambi | 2023 | 3 | -7.984 | 36.818 | F | 1236-VO-TZ-OKUMU-VMF00261 | 1.3 | 1 | TZA | Morogoro | TZ-16 | Kilombero | funestus | TZ-16_fune_2023 | TZ-16_fune_2023_03 | TZ-16_fune_2023_Q1 | TZ-16_Kilombero_fune_2023 | TZ-16_Kilombero_fune_2023_03 | TZ-16_Kilombero_fune_2023_Q1 |
| VBS85448 | TZ_MOR_38 | Joel Odero | Tanzania | Ikwambi | 2023 | 3 | -7.984 | 36.818 | F | 1236-VO-TZ-OKUMU-VMF00261 | 1.3 | 1 | TZA | Morogoro | TZ-16 | Kilombero | funestus | TZ-16_fune_2023 | TZ-16_fune_2023_03 | TZ-16_fune_2023_Q1 | TZ-16_Kilombero_fune_2023 | TZ-16_Kilombero_fune_2023_03 | TZ-16_Kilombero_fune_2023_Q1 |
| VBS85449 | TZ_MOR_37 | Joel Odero | Tanzania | Ikwambi | 2023 | 3 | -7.984 | 36.818 | F | 1236-VO-TZ-OKUMU-VMF00261 | 1.3 | 1 | TZA | Morogoro | TZ-16 | Kilombero | funestus | TZ-16_fune_2023 | TZ-16_fune_2023_03 | TZ-16_fune_2023_Q1 | TZ-16_Kilombero_fune_2023 | TZ-16_Kilombero_fune_2023_03 | TZ-16_Kilombero_fune_2023_Q1 |
| VBS85450 | TZ_MOR_36 | Joel Odero | Tanzania | Ikwambi | 2023 | 3 | -7.984 | 36.818 | F | 1236-VO-TZ-OKUMU-VMF00261 | 1.3 | 1 | TZA | Morogoro | TZ-16 | Kilombero | funestus | TZ-16_fune_2023 | TZ-16_fune_2023_03 | TZ-16_fune_2023_Q1 | TZ-16_Kilombero_fune_2023 | TZ-16_Kilombero_fune_2023_03 | TZ-16_Kilombero_fune_2023_Q1 |
| VBS85451 | TZ_MOR_35 | Joel Odero | Tanzania | Ikwambi | 2023 | 3 | -7.984 | 36.818 | F | 1236-VO-TZ-OKUMU-VMF00261 | 1.3 | 1 | TZA | Morogoro | TZ-16 | Kilombero | funestus | TZ-16_fune_2023 | TZ-16_fune_2023_03 | TZ-16_fune_2023_Q1 | TZ-16_Kilombero_fune_2023 | TZ-16_Kilombero_fune_2023_03 | TZ-16_Kilombero_fune_2023_Q1 |
| VBS85452 | TZ_MOR_34 | Joel Odero | Tanzania | Ikwambi | 2023 | 3 | -7.984 | 36.818 | F | 1236-VO-TZ-OKUMU-VMF00261 | 1.3 | 1 | TZA | Morogoro | TZ-16 | Kilombero | funestus | TZ-16_fune_2023 | TZ-16_fune_2023_03 | TZ-16_fune_2023_Q1 | TZ-16_Kilombero_fune_2023 | TZ-16_Kilombero_fune_2023_03 | TZ-16_Kilombero_fune_2023_Q1 |
| VBS85453 | TZ_MOR_33 | Joel Odero | Tanzania | Ikwambi | 2023 | 3 | -7.984 | 36.818 | F | 1236-VO-TZ-OKUMU-VMF00261 | 1.3 | 1 | TZA | Morogoro | TZ-16 | Kilombero | funestus | TZ-16_fune_2023 | TZ-16_fune_2023_03 | TZ-16_fune_2023_Q1 | TZ-16_Kilombero_fune_2023 | TZ-16_Kilombero_fune_2023_03 | TZ-16_Kilombero_fune_2023_Q1 |
| VBS85454 | TZ_MOR_32 | Joel Odero | Tanzania | Ikwambi | 2023 | 3 | -7.984 | 36.818 | F | 1236-VO-TZ-OKUMU-VMF00261 | 1.3 | 1 | TZA | Morogoro | TZ-16 | Kilombero | funestus | TZ-16_fune_2023 | TZ-16_fune_2023_03 | TZ-16_fune_2023_Q1 | TZ-16_Kilombero_fune_2023 | TZ-16_Kilombero_fune_2023_03 | TZ-16_Kilombero_fune_2023_Q1 |
| VBS85455 | TZ_MOR_31 | Joel Odero | Tanzania | Ikwambi | 2023 | 3 | -7.984 | 36.818 | F | 1236-VO-TZ-OKUMU-VMF00261 | 1.3 | 1 | TZA | Morogoro | TZ-16 | Kilombero | funestus | TZ-16_fune_2023 | TZ-16_fune_2023_03 | TZ-16_fune_2023_Q1 | TZ-16_Kilombero_fune_2023 | TZ-16_Kilombero_fune_2023_03 | TZ-16_Kilombero_fune_2023_Q1 |
| VBS85456 | TZ_KAT_54 | Joel Odero | Tanzania | Vikonge | 2022 | 3 | -6.091 | 30.936 | F | 1236-VO-TZ-OKUMU-VMF00261 | 1.3 | 1 | TZA | Katavi | TZ-28 | Mpanda | funestus | TZ-28_fune_2022 | TZ-28_fune_2022_03 | TZ-28_fune_2022_Q1 | TZ-28_Mpanda_fune_2022 | TZ-28_Mpanda_fune_2022_03 | TZ-28_Mpanda_fune_2022_Q1 |
| VBS85457 | TZ_KAT_53 | Joel Odero | Tanzania | Vikonge | 2022 | 3 | -6.091 | 30.936 | F | 1236-VO-TZ-OKUMU-VMF00261 | 1.3 | 1 | TZA | Katavi | TZ-28 | Mpanda | funestus | TZ-28_fune_2022 | TZ-28_fune_2022_03 | TZ-28_fune_2022_Q1 | TZ-28_Mpanda_fune_2022 | TZ-28_Mpanda_fune_2022_03 | TZ-28_Mpanda_fune_2022_Q1 |
| VBS85458 | TZ_KAT_52 | Joel Odero | Tanzania | Vikonge | 2022 | 3 | -6.091 | 30.936 | F | 1236-VO-TZ-OKUMU-VMF00261 | 1.3 | 1 | TZA | Katavi | TZ-28 | Mpanda | funestus | TZ-28_fune_2022 | TZ-28_fune_2022_03 | TZ-28_fune_2022_Q1 | TZ-28_Mpanda_fune_2022 | TZ-28_Mpanda_fune_2022_03 | TZ-28_Mpanda_fune_2022_Q1 |
| VBS85459 | TZ_KAT_51 | Joel Odero | Tanzania | Vikonge | 2022 | 3 | -6.091 | 30.936 | F | 1236-VO-TZ-OKUMU-VMF00261 | 1.3 | 1 | TZA | Katavi | TZ-28 | Mpanda | funestus | TZ-28_fune_2022 | TZ-28_fune_2022_03 | TZ-28_fune_2022_Q1 | TZ-28_Mpanda_fune_2022 | TZ-28_Mpanda_fune_2022_03 | TZ-28_Mpanda_fune_2022_Q1 |
| VBS85460 | TZ_KAT_50 | Joel Odero | Tanzania | Vikonge | 2022 | 3 | -6.091 | 30.936 | F | 1236-VO-TZ-OKUMU-VMF00261 | 1.3 | 1 | TZA | Katavi | TZ-28 | Mpanda | funestus | TZ-28_fune_2022 | TZ-28_fune_2022_03 | TZ-28_fune_2022_Q1 | TZ-28_Mpanda_fune_2022 | TZ-28_Mpanda_fune_2022_03 | TZ-28_Mpanda_fune_2022_Q1 |
| VBS85461 | TZ_KAT_49 | Joel Odero | Tanzania | Vikonge | 2022 | 3 | -6.091 | 30.936 | F | 1236-VO-TZ-OKUMU-VMF00261 | 1.3 | 1 | TZA | Katavi | TZ-28 | Mpanda | funestus | TZ-28_fune_2022 | TZ-28_fune_2022_03 | TZ-28_fune_2022_Q1 | TZ-28_Mpanda_fune_2022 | TZ-28_Mpanda_fune_2022_03 | TZ-28_Mpanda_fune_2022_Q1 |
| VBS85462 | TZ_KAT_48 | Joel Odero | Tanzania | Vikonge | 2022 | 3 | -6.091 | 30.936 | F | 1236-VO-TZ-OKUMU-VMF00261 | 1.3 | 1 | TZA | Katavi | TZ-28 | Mpanda | funestus | TZ-28_fune_2022 | TZ-28_fune_2022_03 | TZ-28_fune_2022_Q1 | TZ-28_Mpanda_fune_2022 | TZ-28_Mpanda_fune_2022_03 | TZ-28_Mpanda_fune_2022_Q1 |
| VBS85463 | TZ_KAT_47 | Joel Odero | Tanzania | Vikonge | 2022 | 3 | -6.091 | 30.936 | F | 1236-VO-TZ-OKUMU-VMF00261 | 1.3 | 1 | TZA | Katavi | TZ-28 | Mpanda | funestus | TZ-28_fune_2022 | TZ-28_fune_2022_03 | TZ-28_fune_2022_Q1 | TZ-28_Mpanda_fune_2022 | TZ-28_Mpanda_fune_2022_03 | TZ-28_Mpanda_fune_2022_Q1 |
| VBS85464 | TZ_KAT_46 | Joel Odero | Tanzania | Vikonge | 2022 | 3 | -6.091 | 30.936 | F | 1236-VO-TZ-OKUMU-VMF00261 | 1.3 | 1 | TZA | Katavi | TZ-28 | Mpanda | funestus | TZ-28_fune_2022 | TZ-28_fune_2022_03 | TZ-28_fune_2022_Q1 | TZ-28_Mpanda_fune_2022 | TZ-28_Mpanda_fune_2022_03 | TZ-28_Mpanda_fune_2022_Q1 |
| VBS85465 | TZ_KAT_45 | Joel Odero | Tanzania | Vikonge | 2022 | 3 | -6.091 | 30.936 | F | 1236-VO-TZ-OKUMU-VMF00261 | 1.3 | 1 | TZA | Katavi | TZ-28 | Mpanda | funestus | TZ-28_fune_2022 | TZ-28_fune_2022_03 | TZ-28_fune_2022_Q1 | TZ-28_Mpanda_fune_2022 | TZ-28_Mpanda_fune_2022_03 | TZ-28_Mpanda_fune_2022_Q1 |
| VBS85466 | TZ_KAT_44 | Joel Odero | Tanzania | Vikonge | 2022 | 3 | -6.091 | 30.936 | F | 1236-VO-TZ-OKUMU-VMF00261 | 1.3 | 1 | TZA | Katavi | TZ-28 | Mpanda | funestus | TZ-28_fune_2022 | TZ-28_fune_2022_03 | TZ-28_fune_2022_Q1 | TZ-28_Mpanda_fune_2022 | TZ-28_Mpanda_fune_2022_03 | TZ-28_Mpanda_fune_2022_Q1 |
| VBS85467 | TZ_KAT_43 | Joel Odero | Tanzania | Vikonge | 2022 | 3 | -6.091 | 30.936 | F | 1236-VO-TZ-OKUMU-VMF00261 | 1.3 | 1 | TZA | Katavi | TZ-28 | Mpanda | funestus | TZ-28_fune_2022 | TZ-28_fune_2022_03 | TZ-28_fune_2022_Q1 | TZ-28_Mpanda_fune_2022 | TZ-28_Mpanda_fune_2022_03 | TZ-28_Mpanda_fune_2022_Q1 |
| VBS85468 | TZ_KAT_42 | Joel Odero | Tanzania | Vikonge | 2022 | 3 | -6.091 | 30.936 | F | 1236-VO-TZ-OKUMU-VMF00261 | 1.3 | 1 | TZA | Katavi | TZ-28 | Mpanda | funestus | TZ-28_fune_2022 | TZ-28_fune_2022_03 | TZ-28_fune_2022_Q1 | TZ-28_Mpanda_fune_2022 | TZ-28_Mpanda_fune_2022_03 | TZ-28_Mpanda_fune_2022_Q1 |
| VBS85469 | TZ_KAT_41 | Joel Odero | Tanzania | Vikonge | 2022 | 3 | -6.091 | 30.936 | F | 1236-VO-TZ-OKUMU-VMF00261 | 1.3 | 1 | TZA | Katavi | TZ-28 | Mpanda | funestus | TZ-28_fune_2022 | TZ-28_fune_2022_03 | TZ-28_fune_2022_Q1 | TZ-28_Mpanda_fune_2022 | TZ-28_Mpanda_fune_2022_03 | TZ-28_Mpanda_fune_2022_Q1 |
| VBS85470 | TZ_KAT_40 | Joel Odero | Tanzania | Vikonge | 2022 | 3 | -6.091 | 30.936 | F | 1236-VO-TZ-OKUMU-VMF00261 | 1.3 | 1 | TZA | Katavi | TZ-28 | Mpanda | funestus | TZ-28_fune_2022 | TZ-28_fune_2022_03 | TZ-28_fune_2022_Q1 | TZ-28_Mpanda_fune_2022 | TZ-28_Mpanda_fune_2022_03 | TZ-28_Mpanda_fune_2022_Q1 |
| VBS85471 | TZ_KAT_39 | Joel Odero | Tanzania | Vikonge | 2022 | 3 | -6.091 | 30.936 | F | 1236-VO-TZ-OKUMU-VMF00261 | 1.3 | 1 | TZA | Katavi | TZ-28 | Mpanda | funestus | TZ-28_fune_2022 | TZ-28_fune_2022_03 | TZ-28_fune_2022_Q1 | TZ-28_Mpanda_fune_2022 | TZ-28_Mpanda_fune_2022_03 | TZ-28_Mpanda_fune_2022_Q1 |
| VBS85472 | TZ_KAT_38 | Joel Odero | Tanzania | Vikonge | 2022 | 3 | -6.091 | 30.936 | F | 1236-VO-TZ-OKUMU-VMF00261 | 1.3 | 1 | TZA | Katavi | TZ-28 | Mpanda | funestus | TZ-28_fune_2022 | TZ-28_fune_2022_03 | TZ-28_fune_2022_Q1 | TZ-28_Mpanda_fune_2022 | TZ-28_Mpanda_fune_2022_03 | TZ-28_Mpanda_fune_2022_Q1 |
| VBS85473 | TZ_KAT_37 | Joel Odero | Tanzania | Vikonge | 2022 | 3 | -6.091 | 30.936 | F | 1236-VO-TZ-OKUMU-VMF00261 | 1.3 | 1 | TZA | Katavi | TZ-28 | Mpanda | funestus | TZ-28_fune_2022 | TZ-28_fune_2022_03 | TZ-28_fune_2022_Q1 | TZ-28_Mpanda_fune_2022 | TZ-28_Mpanda_fune_2022_03 | TZ-28_Mpanda_fune_2022_Q1 |
| VBS85474 | TZ_KAG_50 | Joel Odero | Tanzania | Byeju | 2022 | 1 | -1.08 | 31.376 | F | 1236-VO-TZ-OKUMU-VMF00261 | 1.3 | 1 | TZA | Kagera | TZ-05 | Missenyi | funestus | TZ-05_fune_2022 | TZ-05_fune_2022_01 | TZ-05_fune_2022_Q1 | TZ-05_Missenyi_fune_2022 | TZ-05_Missenyi_fune_2022_01 | TZ-05_Missenyi_fune_2022_Q1 |
| VBS85475 | TZ_KAG_49 | Joel Odero | Tanzania | Byeju | 2022 | 1 | -1.08 | 31.376 | F | 1236-VO-TZ-OKUMU-VMF00261 | 1.3 | 1 | TZA | Kagera | TZ-05 | Missenyi | funestus | TZ-05_fune_2022 | TZ-05_fune_2022_01 | TZ-05_fune_2022_Q1 | TZ-05_Missenyi_fune_2022 | TZ-05_Missenyi_fune_2022_01 | TZ-05_Missenyi_fune_2022_Q1 |
| VBS85476 | TZ_KAG_48 | Joel Odero | Tanzania | Byeju | 2022 | 1 | -1.08 | 31.376 | F | 1236-VO-TZ-OKUMU-VMF00261 | 1.3 | 1 | TZA | Kagera | TZ-05 | Missenyi | funestus | TZ-05_fune_2022 | TZ-05_fune_2022_01 | TZ-05_fune_2022_Q1 | TZ-05_Missenyi_fune_2022 | TZ-05_Missenyi_fune_2022_01 | TZ-05_Missenyi_fune_2022_Q1 |
| VBS85477 | TZ_KAG_47 | Joel Odero | Tanzania | Byeju | 2022 | 1 | -1.08 | 31.376 | F | 1236-VO-TZ-OKUMU-VMF00261 | 1.3 | 1 | TZA | Kagera | TZ-05 | Missenyi | funestus | TZ-05_fune_2022 | TZ-05_fune_2022_01 | TZ-05_fune_2022_Q1 | TZ-05_Missenyi_fune_2022 | TZ-05_Missenyi_fune_2022_01 | TZ-05_Missenyi_fune_2022_Q1 |
| VBS85478 | TZ_KAG_46 | Joel Odero | Tanzania | Byeju | 2022 | 1 | -1.08 | 31.376 | F | 1236-VO-TZ-OKUMU-VMF00261 | 1.3 | 1 | TZA | Kagera | TZ-05 | Missenyi | funestus | TZ-05_fune_2022 | TZ-05_fune_2022_01 | TZ-05_fune_2022_Q1 | TZ-05_Missenyi_fune_2022 | TZ-05_Missenyi_fune_2022_01 | TZ-05_Missenyi_fune_2022_Q1 |
| VBS85479 | TZ_KAG_45 | Joel Odero | Tanzania | Byeju | 2022 | 1 | -1.08 | 31.376 | F | 1236-VO-TZ-OKUMU-VMF00261 | 1.3 | 1 | TZA | Kagera | TZ-05 | Missenyi | funestus | TZ-05_fune_2022 | TZ-05_fune_2022_01 | TZ-05_fune_2022_Q1 | TZ-05_Missenyi_fune_2022 | TZ-05_Missenyi_fune_2022_01 | TZ-05_Missenyi_fune_2022_Q1 |
| VBS85480 | TZ_KAG_44 | Joel Odero | Tanzania | Byeju | 2022 | 1 | -1.08 | 31.376 | F | 1236-VO-TZ-OKUMU-VMF00261 | 1.3 | 1 | TZA | Kagera | TZ-05 | Missenyi | funestus | TZ-05_fune_2022 | TZ-05_fune_2022_01 | TZ-05_fune_2022_Q1 | TZ-05_Missenyi_fune_2022 | TZ-05_Missenyi_fune_2022_01 | TZ-05_Missenyi_fune_2022_Q1 |
| VBS85481 | TZ_KAG_43 | Joel Odero | Tanzania | Byeju | 2022 | 1 | -1.08 | 31.376 | F | 1236-VO-TZ-OKUMU-VMF00261 | 1.3 | 1 | TZA | Kagera | TZ-05 | Missenyi | funestus | TZ-05_fune_2022 | TZ-05_fune_2022_01 | TZ-05_fune_2022_Q1 | TZ-05_Missenyi_fune_2022 | TZ-05_Missenyi_fune_2022_01 | TZ-05_Missenyi_fune_2022_Q1 |
| VBS85482 | TZ_MTW_45 | Joel Odero | Tanzania | Mahurunga | 2022 | 10 | -10.538 | 40.279 | F | 1236-VO-TZ-OKUMU-VMF00261 | 1.3 | 4 | TZA | Mtwara | TZ-17 | Mtwara | funestus | TZ-17_fune_2022 | TZ-17_fune_2022_10 | TZ-17_fune_2022_Q4 | TZ-17_Mtwara_fune_2022 | TZ-17_Mtwara_fune_2022_10 | TZ-17_Mtwara_fune_2022_Q4 |
| VBS85483 | TZ_MTW_47 | Joel Odero | Tanzania | Mahurunga | 2022 | 10 | -10.538 | 40.279 | F | 1236-VO-TZ-OKUMU-VMF00261 | 1.3 | 4 | TZA | Mtwara | TZ-17 | Mtwara | funestus | TZ-17_fune_2022 | TZ-17_fune_2022_10 | TZ-17_fune_2022_Q4 | TZ-17_Mtwara_fune_2022 | TZ-17_Mtwara_fune_2022_10 | TZ-17_Mtwara_fune_2022_Q4 |
| VBS85484 | TZ_MTW_46 | Joel Odero | Tanzania | Mahurunga | 2022 | 10 | -10.538 | 40.279 | F | 1236-VO-TZ-OKUMU-VMF00261 | 1.3 | 4 | TZA | Mtwara | TZ-17 | Mtwara | funestus | TZ-17_fune_2022 | TZ-17_fune_2022_10 | TZ-17_fune_2022_Q4 | TZ-17_Mtwara_fune_2022 | TZ-17_Mtwara_fune_2022_10 | TZ-17_Mtwara_fune_2022_Q4 |
| VBS85485 | TZ_MOR_58 | Joel Odero | Tanzania | Ikwambi | 2023 | 3 | -7.984 | 36.818 | F | 1236-VO-TZ-OKUMU-VMF00261 | 1.3 | 1 | TZA | Morogoro | TZ-16 | Kilombero | funestus | TZ-16_fune_2023 | TZ-16_fune_2023_03 | TZ-16_fune_2023_Q1 | TZ-16_Kilombero_fune_2023 | TZ-16_Kilombero_fune_2023_03 | TZ-16_Kilombero_fune_2023_Q1 |
| VBS85486 | TZ_MOR_57 | Joel Odero | Tanzania | Ikwambi | 2023 | 3 | -7.984 | 36.818 | F | 1236-VO-TZ-OKUMU-VMF00261 | 1.3 | 1 | TZA | Morogoro | TZ-16 | Kilombero | funestus | TZ-16_fune_2023 | TZ-16_fune_2023_03 | TZ-16_fune_2023_Q1 | TZ-16_Kilombero_fune_2023 | TZ-16_Kilombero_fune_2023_03 | TZ-16_Kilombero_fune_2023_Q1 |
| VBS85487 | TZ_MOR_56 | Joel Odero | Tanzania | Ikwambi | 2023 | 3 | -7.984 | 36.818 | F | 1236-VO-TZ-OKUMU-VMF00261 | 1.3 | 1 | TZA | Morogoro | TZ-16 | Kilombero | funestus | TZ-16_fune_2023 | TZ-16_fune_2023_03 | TZ-16_fune_2023_Q1 | TZ-16_Kilombero_fune_2023 | TZ-16_Kilombero_fune_2023_03 | TZ-16_Kilombero_fune_2023_Q1 |
| VBS85488 | TZ_MOR_55 | Joel Odero | Tanzania | Ikwambi | 2023 | 3 | -7.984 | 36.818 | F | 1236-VO-TZ-OKUMU-VMF00261 | 1.3 | 1 | TZA | Morogoro | TZ-16 | Kilombero | funestus | TZ-16_fune_2023 | TZ-16_fune_2023_03 | TZ-16_fune_2023_Q1 | TZ-16_Kilombero_fune_2023 | TZ-16_Kilombero_fune_2023_03 | TZ-16_Kilombero_fune_2023_Q1 |
| VBS85489 | TZ_TNG_41 | Joel Odero | Tanzania | Kwamwale | 2022 | 5 | -5.157 | 38.729 | F | 1236-VO-TZ-OKUMU-VMF00261 | 1.3 | 2 | TZA | Tanga | TZ-25 | Muheza | funestus | TZ-25_fune_2022 | TZ-25_fune_2022_05 | TZ-25_fune_2022_Q2 | TZ-25_Muheza_fune_2022 | TZ-25_Muheza_fune_2022_05 | TZ-25_Muheza_fune_2022_Q2 |
| VBS85490 | TZ_TNG_40 | Joel Odero | Tanzania | Kwamwale | 2022 | 5 | -5.157 | 38.729 | F | 1236-VO-TZ-OKUMU-VMF00261 | 1.3 | 2 | TZA | Tanga | TZ-25 | Muheza | funestus | TZ-25_fune_2022 | TZ-25_fune_2022_05 | TZ-25_fune_2022_Q2 | TZ-25_Muheza_fune_2022 | TZ-25_Muheza_fune_2022_05 | TZ-25_Muheza_fune_2022_Q2 |
| VBS85491 | TZ_TNG_39 | Joel Odero | Tanzania | Kwamwale | 2022 | 5 | -5.157 | 38.729 | F | 1236-VO-TZ-OKUMU-VMF00261 | 1.3 | 2 | TZA | Tanga | TZ-25 | Muheza | funestus | TZ-25_fune_2022 | TZ-25_fune_2022_05 | TZ-25_fune_2022_Q2 | TZ-25_Muheza_fune_2022 | TZ-25_Muheza_fune_2022_05 | TZ-25_Muheza_fune_2022_Q2 |
| VBS85492 | TZ_TNG_38 | Joel Odero | Tanzania | Kwamwale | 2022 | 5 | -5.157 | 38.729 | F | 1236-VO-TZ-OKUMU-VMF00261 | 1.3 | 2 | TZA | Tanga | TZ-25 | Muheza | funestus | TZ-25_fune_2022 | TZ-25_fune_2022_05 | TZ-25_fune_2022_Q2 | TZ-25_Muheza_fune_2022 | TZ-25_Muheza_fune_2022_05 | TZ-25_Muheza_fune_2022_Q2 |
| VBS85493 | TZ_TNG_37 | Joel Odero | Tanzania | Kwamwale | 2022 | 5 | -5.157 | 38.729 | F | 1236-VO-TZ-OKUMU-VMF00261 | 1.3 | 2 | TZA | Tanga | TZ-25 | Muheza | funestus | TZ-25_fune_2022 | TZ-25_fune_2022_05 | TZ-25_fune_2022_Q2 | TZ-25_Muheza_fune_2022 | TZ-25_Muheza_fune_2022_05 | TZ-25_Muheza_fune_2022_Q2 |
| VBS85494 | TZ_MTW_69 | Joel Odero | Tanzania | Mahurunga | 2022 | 10 | -10.538 | 40.279 | F | 1236-VO-TZ-OKUMU-VMF00261 | 1.3 | 4 | TZA | Mtwara | TZ-17 | Mtwara | funestus | TZ-17_fune_2022 | TZ-17_fune_2022_10 | TZ-17_fune_2022_Q4 | TZ-17_Mtwara_fune_2022 | TZ-17_Mtwara_fune_2022_10 | TZ-17_Mtwara_fune_2022_Q4 |
| VBS85495 | TZ_MTW_68 | Joel Odero | Tanzania | Mahurunga | 2022 | 10 | -10.538 | 40.279 | F | 1236-VO-TZ-OKUMU-VMF00261 | 1.3 | 4 | TZA | Mtwara | TZ-17 | Mtwara | funestus | TZ-17_fune_2022 | TZ-17_fune_2022_10 | TZ-17_fune_2022_Q4 | TZ-17_Mtwara_fune_2022 | TZ-17_Mtwara_fune_2022_10 | TZ-17_Mtwara_fune_2022_Q4 |
| VBS85496 | TZ_MTW_64 | Joel Odero | Tanzania | Mahurunga | 2022 | 10 | -10.538 | 40.279 | F | 1236-VO-TZ-OKUMU-VMF00261 | 1.3 | 4 | TZA | Mtwara | TZ-17 | Mtwara | funestus | TZ-17_fune_2022 | TZ-17_fune_2022_10 | TZ-17_fune_2022_Q4 | TZ-17_Mtwara_fune_2022 | TZ-17_Mtwara_fune_2022_10 | TZ-17_Mtwara_fune_2022_Q4 |
| VBS85497 | TZ_MTW_61 | Joel Odero | Tanzania | Mahurunga | 2022 | 10 | -10.538 | 40.279 | F | 1236-VO-TZ-OKUMU-VMF00261 | 1.3 | 4 | TZA | Mtwara | TZ-17 | Mtwara | funestus | TZ-17_fune_2022 | TZ-17_fune_2022_10 | TZ-17_fune_2022_Q4 | TZ-17_Mtwara_fune_2022 | TZ-17_Mtwara_fune_2022_10 | TZ-17_Mtwara_fune_2022_Q4 |
| VBS85498 | TZ_MTW_60 | Joel Odero | Tanzania | Mahurunga | 2022 | 10 | -10.538 | 40.279 | F | 1236-VO-TZ-OKUMU-VMF00261 | 1.3 | 4 | TZA | Mtwara | TZ-17 | Mtwara | funestus | TZ-17_fune_2022 | TZ-17_fune_2022_10 | TZ-17_fune_2022_Q4 | TZ-17_Mtwara_fune_2022 | TZ-17_Mtwara_fune_2022_10 | TZ-17_Mtwara_fune_2022_Q4 |
| VBS85499 | TZ_MTW_55 | Joel Odero | Tanzania | Mahurunga | 2022 | 10 | -10.538 | 40.279 | F | 1236-VO-TZ-OKUMU-VMF00261 | 1.3 | 4 | TZA | Mtwara | TZ-17 | Mtwara | funestus | TZ-17_fune_2022 | TZ-17_fune_2022_10 | TZ-17_fune_2022_Q4 | TZ-17_Mtwara_fune_2022 | TZ-17_Mtwara_fune_2022_10 | TZ-17_Mtwara_fune_2022_Q4 |
| VBS85500 | TZ_MTW_52 | Joel Odero | Tanzania | Mahurunga | 2022 | 10 | -10.538 | 40.279 | F | 1236-VO-TZ-OKUMU-VMF00261 | 1.3 | 4 | TZA | Mtwara | TZ-17 | Mtwara | funestus | TZ-17_fune_2022 | TZ-17_fune_2022_10 | TZ-17_fune_2022_Q4 | TZ-17_Mtwara_fune_2022 | TZ-17_Mtwara_fune_2022_10 | TZ-17_Mtwara_fune_2022_Q4 |
| VBS85501 | TZ_MTW_51 | Joel Odero | Tanzania | Mahurunga | 2022 | 10 | -10.538 | 40.279 | F | 1236-VO-TZ-OKUMU-VMF00261 | 1.3 | 4 | TZA | Mtwara | TZ-17 | Mtwara | funestus | TZ-17_fune_2022 | TZ-17_fune_2022_10 | TZ-17_fune_2022_Q4 | TZ-17_Mtwara_fune_2022 | TZ-17_Mtwara_fune_2022_10 | TZ-17_Mtwara_fune_2022_Q4 |
| VBS85502 | TZ_MWA_39 | Joel Odero | Tanzania | Nyangomango | 2023 | 3 | -3.05 | 32.8 | F | 1236-VO-TZ-OKUMU-VMF00261 | 1.3 | 1 | TZA | Mwanza | TZ-18 | Misungwi | funestus | TZ-18_fune_2023 | TZ-18_fune_2023_03 | TZ-18_fune_2023_Q1 | TZ-18_Misungwi_fune_2023 | TZ-18_Misungwi_fune_2023_03 | TZ-18_Misungwi_fune_2023_Q1 |
| VBS85503 | TZ_MWA_38 | Joel Odero | Tanzania | Nyangomango | 2023 | 3 | -3.05 | 32.8 | F | 1236-VO-TZ-OKUMU-VMF00261 | 1.3 | 1 | TZA | Mwanza | TZ-18 | Misungwi | funestus | TZ-18_fune_2023 | TZ-18_fune_2023_03 | TZ-18_fune_2023_Q1 | TZ-18_Misungwi_fune_2023 | TZ-18_Misungwi_fune_2023_03 | TZ-18_Misungwi_fune_2023_Q1 |
| VBS85504 | TZ_MWA_37 | Joel Odero | Tanzania | Nyangomango | 2023 | 3 | -3.05 | 32.8 | F | 1236-VO-TZ-OKUMU-VMF00261 | 1.3 | 1 | TZA | Mwanza | TZ-18 | Misungwi | funestus | TZ-18_fune_2023 | TZ-18_fune_2023_03 | TZ-18_fune_2023_Q1 | TZ-18_Misungwi_fune_2023 | TZ-18_Misungwi_fune_2023_03 | TZ-18_Misungwi_fune_2023_Q1 |
| VBS85506 | TZ_MWA_33 | Joel Odero | Tanzania | Nyangomango | 2023 | 3 | -3.05 | 32.8 | F | 1236-VO-TZ-OKUMU-VMF00261 | 1.3 | 1 | TZA | Mwanza | TZ-18 | Misungwi | funestus | TZ-18_fune_2023 | TZ-18_fune_2023_03 | TZ-18_fune_2023_Q1 | TZ-18_Misungwi_fune_2023 | TZ-18_Misungwi_fune_2023_03 | TZ-18_Misungwi_fune_2023_Q1 |
| VBS85508 | TZ_MWA_31 | Joel Odero | Tanzania | Nyangomango | 2023 | 3 | -3.05 | 32.8 | F | 1236-VO-TZ-OKUMU-VMF00261 | 1.3 | 1 | TZA | Mwanza | TZ-18 | Misungwi | funestus | TZ-18_fune_2023 | TZ-18_fune_2023_03 | TZ-18_fune_2023_Q1 | TZ-18_Misungwi_fune_2023 | TZ-18_Misungwi_fune_2023_03 | TZ-18_Misungwi_fune_2023_Q1 |
| VBS85509 | TZ_MWA_29 | Joel Odero | Tanzania | Nyangomango | 2023 | 3 | -3.05 | 32.8 | F | 1236-VO-TZ-OKUMU-VMF00261 | 1.3 | 1 | TZA | Mwanza | TZ-18 | Misungwi | funestus | TZ-18_fune_2023 | TZ-18_fune_2023_03 | TZ-18_fune_2023_Q1 | TZ-18_Misungwi_fune_2023 | TZ-18_Misungwi_fune_2023_03 | TZ-18_Misungwi_fune_2023_Q1 |
| VBS85510 | TZ_MWA_28 | Joel Odero | Tanzania | Nyangomango | 2023 | 3 | -3.05 | 32.8 | F | 1236-VO-TZ-OKUMU-VMF00261 | 1.3 | 1 | TZA | Mwanza | TZ-18 | Misungwi | funestus | TZ-18_fune_2023 | TZ-18_fune_2023_03 | TZ-18_fune_2023_Q1 | TZ-18_Misungwi_fune_2023 | TZ-18_Misungwi_fune_2023_03 | TZ-18_Misungwi_fune_2023_Q1 |
| VBS85511 | TZ_MWA_27 | Joel Odero | Tanzania | Nyangomango | 2023 | 3 | -3.05 | 32.8 | F | 1236-VO-TZ-OKUMU-VMF00261 | 1.3 | 1 | TZA | Mwanza | TZ-18 | Misungwi | funestus | TZ-18_fune_2023 | TZ-18_fune_2023_03 | TZ-18_fune_2023_Q1 | TZ-18_Misungwi_fune_2023 | TZ-18_Misungwi_fune_2023_03 | TZ-18_Misungwi_fune_2023_Q1 |
| VBS85512 | TZ_MWA_26 | Joel Odero | Tanzania | Nyangomango | 2023 | 3 | -3.05 | 32.8 | F | 1236-VO-TZ-OKUMU-VMF00261 | 1.3 | 1 | TZA | Mwanza | TZ-18 | Misungwi | funestus | TZ-18_fune_2023 | TZ-18_fune_2023_03 | TZ-18_fune_2023_Q1 | TZ-18_Misungwi_fune_2023 | TZ-18_Misungwi_fune_2023_03 | TZ-18_Misungwi_fune_2023_Q1 |
| VBS85513 | TZ_MWA_25 | Joel Odero | Tanzania | Nyangomango | 2023 | 3 | -3.05 | 32.8 | F | 1236-VO-TZ-OKUMU-VMF00261 | 1.3 | 1 | TZA | Mwanza | TZ-18 | Misungwi | funestus | TZ-18_fune_2023 | TZ-18_fune_2023_03 | TZ-18_fune_2023_Q1 | TZ-18_Misungwi_fune_2023 | TZ-18_Misungwi_fune_2023_03 | TZ-18_Misungwi_fune_2023_Q1 |
| VBS85514 | TZ_MWA_24 | Joel Odero | Tanzania | Nyangomango | 2023 | 3 | -3.05 | 32.8 | F | 1236-VO-TZ-OKUMU-VMF00261 | 1.3 | 1 | TZA | Mwanza | TZ-18 | Misungwi | funestus | TZ-18_fune_2023 | TZ-18_fune_2023_03 | TZ-18_fune_2023_Q1 | TZ-18_Misungwi_fune_2023 | TZ-18_Misungwi_fune_2023_03 | TZ-18_Misungwi_fune_2023_Q1 |
| VBS85515 | TZ_MWA_23 | Joel Odero | Tanzania | Nyangomango | 2023 | 3 | -3.05 | 32.8 | F | 1236-VO-TZ-OKUMU-VMF00261 | 1.3 | 1 | TZA | Mwanza | TZ-18 | Misungwi | funestus | TZ-18_fune_2023 | TZ-18_fune_2023_03 | TZ-18_fune_2023_Q1 | TZ-18_Misungwi_fune_2023 | TZ-18_Misungwi_fune_2023_03 | TZ-18_Misungwi_fune_2023_Q1 |
| VBS85516 | TZ_MWA_22 | Joel Odero | Tanzania | Nyangomango | 2023 | 3 | -3.05 | 32.8 | F | 1236-VO-TZ-OKUMU-VMF00261 | 1.3 | 1 | TZA | Mwanza | TZ-18 | Misungwi | funestus | TZ-18_fune_2023 | TZ-18_fune_2023_03 | TZ-18_fune_2023_Q1 | TZ-18_Misungwi_fune_2023 | TZ-18_Misungwi_fune_2023_03 | TZ-18_Misungwi_fune_2023_Q1 |
| VBS85517 | TZ_MWA_21 | Joel Odero | Tanzania | Nyangomango | 2023 | 3 | -3.05 | 32.8 | F | 1236-VO-TZ-OKUMU-VMF00261 | 1.3 | 1 | TZA | Mwanza | TZ-18 | Misungwi | funestus | TZ-18_fune_2023 | TZ-18_fune_2023_03 | TZ-18_fune_2023_Q1 | TZ-18_Misungwi_fune_2023 | TZ-18_Misungwi_fune_2023_03 | TZ-18_Misungwi_fune_2023_Q1 |
| VBS85518 | TZ_MWA_20 | Joel Odero | Tanzania | Nyangomango | 2023 | 3 | -3.05 | 32.8 | F | 1236-VO-TZ-OKUMU-VMF00261 | 1.3 | 1 | TZA | Mwanza | TZ-18 | Misungwi | funestus | TZ-18_fune_2023 | TZ-18_fune_2023_03 | TZ-18_fune_2023_Q1 | TZ-18_Misungwi_fune_2023 | TZ-18_Misungwi_fune_2023_03 | TZ-18_Misungwi_fune_2023_Q1 |
| VBS85519 | TZ_MWA_19 | Joel Odero | Tanzania | Nyangomango | 2023 | 3 | -3.05 | 32.8 | F | 1236-VO-TZ-OKUMU-VMF00261 | 1.3 | 1 | TZA | Mwanza | TZ-18 | Misungwi | funestus | TZ-18_fune_2023 | TZ-18_fune_2023_03 | TZ-18_fune_2023_Q1 | TZ-18_Misungwi_fune_2023 | TZ-18_Misungwi_fune_2023_03 | TZ-18_Misungwi_fune_2023_Q1 |
| VBS85521 | TZ_MWA_17 | Joel Odero | Tanzania | Nyangomango | 2023 | 3 | -3.05 | 32.8 | F | 1236-VO-TZ-OKUMU-VMF00261 | 1.3 | 1 | TZA | Mwanza | TZ-18 | Misungwi | funestus | TZ-18_fune_2023 | TZ-18_fune_2023_03 | TZ-18_fune_2023_Q1 | TZ-18_Misungwi_fune_2023 | TZ-18_Misungwi_fune_2023_03 | TZ-18_Misungwi_fune_2023_Q1 |
| VBS85522 | TZ_MWA_16 | Joel Odero | Tanzania | Nyangomango | 2023 | 3 | -3.05 | 32.8 | F | 1236-VO-TZ-OKUMU-VMF00261 | 1.3 | 1 | TZA | Mwanza | TZ-18 | Misungwi | funestus | TZ-18_fune_2023 | TZ-18_fune_2023_03 | TZ-18_fune_2023_Q1 | TZ-18_Misungwi_fune_2023 | TZ-18_Misungwi_fune_2023_03 | TZ-18_Misungwi_fune_2023_Q1 |
| VBS85523 | TZ_MWA_14 | Joel Odero | Tanzania | Nyangomango | 2023 | 3 | -3.05 | 32.8 | F | 1236-VO-TZ-OKUMU-VMF00261 | 1.3 | 1 | TZA | Mwanza | TZ-18 | Misungwi | funestus | TZ-18_fune_2023 | TZ-18_fune_2023_03 | TZ-18_fune_2023_Q1 | TZ-18_Misungwi_fune_2023 | TZ-18_Misungwi_fune_2023_03 | TZ-18_Misungwi_fune_2023_Q1 |
| VBS85524 | TZ_MWA_13 | Joel Odero | Tanzania | Nyangomango | 2023 | 3 | -3.05 | 32.8 | F | 1236-VO-TZ-OKUMU-VMF00261 | 1.3 | 1 | TZA | Mwanza | TZ-18 | Misungwi | funestus | TZ-18_fune_2023 | TZ-18_fune_2023_03 | TZ-18_fune_2023_Q1 | TZ-18_Misungwi_fune_2023 | TZ-18_Misungwi_fune_2023_03 | TZ-18_Misungwi_fune_2023_Q1 |
| VBS85525 | TZ_MWA_12 | Joel Odero | Tanzania | Nyangomango | 2023 | 3 | -3.05 | 32.8 | F | 1236-VO-TZ-OKUMU-VMF00261 | 1.3 | 1 | TZA | Mwanza | TZ-18 | Misungwi | funestus | TZ-18_fune_2023 | TZ-18_fune_2023_03 | TZ-18_fune_2023_Q1 | TZ-18_Misungwi_fune_2023 | TZ-18_Misungwi_fune_2023_03 | TZ-18_Misungwi_fune_2023_Q1 |
| VBS85526 | TZ_MWA_11 | Joel Odero | Tanzania | Nyangomango | 2023 | 3 | -3.05 | 32.8 | F | 1236-VO-TZ-OKUMU-VMF00261 | 1.3 | 1 | TZA | Mwanza | TZ-18 | Misungwi | funestus | TZ-18_fune_2023 | TZ-18_fune_2023_03 | TZ-18_fune_2023_Q1 | TZ-18_Misungwi_fune_2023 | TZ-18_Misungwi_fune_2023_03 | TZ-18_Misungwi_fune_2023_Q1 |
| VBS85527 | TZ_MWA_10 | Joel Odero | Tanzania | Nyangomango | 2023 | 3 | -3.05 | 32.8 | F | 1236-VO-TZ-OKUMU-VMF00261 | 1.3 | 1 | TZA | Mwanza | TZ-18 | Misungwi | funestus | TZ-18_fune_2023 | TZ-18_fune_2023_03 | TZ-18_fune_2023_Q1 | TZ-18_Misungwi_fune_2023 | TZ-18_Misungwi_fune_2023_03 | TZ-18_Misungwi_fune_2023_Q1 |
| VBS85528 | TZ_MWA_6 | Joel Odero | Tanzania | Nyangomango | 2023 | 3 | -3.05 | 32.8 | F | 1236-VO-TZ-OKUMU-VMF00261 | 1.3 | 1 | TZA | Mwanza | TZ-18 | Misungwi | funestus | TZ-18_fune_2023 | TZ-18_fune_2023_03 | TZ-18_fune_2023_Q1 | TZ-18_Misungwi_fune_2023 | TZ-18_Misungwi_fune_2023_03 | TZ-18_Misungwi_fune_2023_Q1 |
| VBS85529 | TZ_MWA_5 | Joel Odero | Tanzania | Nyangomango | 2023 | 3 | -3.05 | 32.8 | F | 1236-VO-TZ-OKUMU-VMF00261 | 1.3 | 1 | TZA | Mwanza | TZ-18 | Misungwi | funestus | TZ-18_fune_2023 | TZ-18_fune_2023_03 | TZ-18_fune_2023_Q1 | TZ-18_Misungwi_fune_2023 | TZ-18_Misungwi_fune_2023_03 | TZ-18_Misungwi_fune_2023_Q1 |
| VBS85530 | TZ_MWA_3 | Joel Odero | Tanzania | Nyangomango | 2023 | 3 | -3.05 | 32.8 | F | 1236-VO-TZ-OKUMU-VMF00261 | 1.3 | 1 | TZA | Mwanza | TZ-18 | Misungwi | funestus | TZ-18_fune_2023 | TZ-18_fune_2023_03 | TZ-18_fune_2023_Q1 | TZ-18_Misungwi_fune_2023 | TZ-18_Misungwi_fune_2023_03 | TZ-18_Misungwi_fune_2023_Q1 |
| VBS85531 | TZ_MWA_2 | Joel Odero | Tanzania | Nyangomango | 2023 | 3 | -3.05 | 32.8 | F | 1236-VO-TZ-OKUMU-VMF00261 | 1.3 | 1 | TZA | Mwanza | TZ-18 | Misungwi | funestus | TZ-18_fune_2023 | TZ-18_fune_2023_03 | TZ-18_fune_2023_Q1 | TZ-18_Misungwi_fune_2023 | TZ-18_Misungwi_fune_2023_03 | TZ-18_Misungwi_fune_2023_Q1 |
| VBS85532 | TZ_MWA_1 | Joel Odero | Tanzania | Nyangomango | 2023 | 3 | -3.05 | 32.8 | F | 1236-VO-TZ-OKUMU-VMF00261 | 1.3 | 1 | TZA | Mwanza | TZ-18 | Misungwi | funestus | TZ-18_fune_2023 | TZ-18_fune_2023_03 | TZ-18_fune_2023_Q1 | TZ-18_Misungwi_fune_2023 | TZ-18_Misungwi_fune_2023_03 | TZ-18_Misungwi_fune_2023_Q1 |

**Supplementary Table 2: Samples location and number used in the G123 selection scans. Only locations with n>20**

| Admin1_Name | Sample No. |
|---|---|
| Dodoma | 32 |
| Kagera | 32 |
| Katavi | 33 |
| Kigoma | 1 |
| Lindi | 30 |
| Morogoro | 64 |
| Mtwara | 32 |
| Mwanza | 28 |
| Pwani | 12 |
| Ruvuma | 37 |
| Tanga | 33 |

**Supplementary Table 3: List of primer sets and cycling conditions used to amplify Anopheles funestus VGSC gene**

| | |
|---|---|
| kdrL976F_FWD | 5'- TGTGCGGTGAATGGATCGAA-3' |
| kdrL976F_REV | 5'-CGCTTCAGCGATCTTGTTGG-3' |
| kdrP1842S-FWI | 5'-CTACCCGGGAAATTGTGGCT-3' |
| kdrP1842S_RE' | 5'-TGCCACCATCGTTTCCGTTA-3' |

Cycling conditions

| | | |
|---|---|---|
| 94 °C | 5min | |
| 94 °C | 1min | |
| 58 °C | 30sec | 35 cycles |
| 72 °C | 30sec | |
| 72 °C | 10min | |
| 4 °C | hold | |
